# Human land-use impacts viral diversity and abundance in a New Zealand river

**DOI:** 10.1101/2022.01.04.474996

**Authors:** Rebecca French, Justine Charon, Callum Le Lay, Chris Muller, Edward C. Holmes

## Abstract

Although water-borne viruses have important implications for the health of humans and other animals, little is known about the impact of human land-use on viral diversity and evolution in water systems such as rivers. We used metagenomic next-generation sequencing to compare the diversity and abundance of viruses at sampling sites along a single river in New Zealand that differed in human land use impact, ranging from pristine to urban. From this we identified 504 putative virus species, of which 97% were novel. Many of the novel viruses were highly divergent, and likely included a new subfamily within the *Parvoviridae*. We identified at least 63 virus species that may infect vertebrates – most likely fish and water birds – from the *Astroviridae, Birnaviridae, Parvoviridae* and *Picornaviridae*. No putative human viruses were detected. Importantly, we observed differences in the composition of viral communities at sites impacted by human land-use (farming and urban) compared to native forest sites (pristine). At the viral species level, the urban sites had higher diversity (327 virus species) than the farming (n=150) and pristine sites (n=119), and more viruses were shared between the urban and farming sites (n=76) than between the pristine and farming or urban sites (n=24). The two farming sites had a lower viral abundance across all host types, while the pristine sites had a higher abundance of viruses associated with animals, plants and fungi. We also identified viruses linked to agriculture and human impact at the river sampling sites in farming and urban areas that were not present at the native forest sites. Overall, our study shows that human land-use can impact viral communities in rivers, such that further work is needed to reduce the impact of intensive farming and urbanization on water systems.

## 1. Introduction

As viruses likely infect all life forms, and often at high abundance, they can be considered an integral part of global ecosystems (Zhang et al. 2018; French and Holmes 2020; Sommers et al. 2021). Until recently, however, there has been a strong bias toward studying viruses in the context of individual disease-causing pathogens, particularly in humans, domestic animals and plants (Zhang et al. 2018). Although understandable, such a bias limits our understanding of their ecology and evolution, how viral abundance and diversity might be shaped by anthropogenic activities, and their role at the ecosystem scale (French and Holmes 2020; Sommers et al. 2021). Clearly, a better understanding of these processes will enable virus evolution and disease emergence to be placed in its true ecological context. As most viruses do not cause disease in their hosts (Roossinck 2015), characterising non-pathogenic viruses will greatly expand our understanding of the composition of the global virosphere.

Metagenomic next-generation sequencing (mNGS) enables the entire virome of a sample to be characterised in an unbiased manner, giving studies of RNA virus diversity and evolution a new perspective (Zhang et al. 2018; Wolf et al. 2020; Nayfach et al. 2021). In particular, mNGS enables the comparison of viral abundance and diversity between groups (animal populations, environments etc.) that was previously not possible on large scales. To date, however, most metagenomic studies of viromes have focused on describing viral diversity without placing it in an appropriate ecological context (Zhang et al. 2018; French and Holmes 2020; Sommers et al. 2021).

Rivers collect water from the land they flow through. As such, their microbial community necessarily reflects the ecological properties of this adjacent land (Van Rossum et al. 2015).

Run-off from farmland, urban areas and sewage discharge directly introduce human and livestock-infecting microbes into rivers, sometimes causing water-borne disease (Ferguson et al. 2003; Alegbeleye and Sant’Ana 2020). For bacteria, it is well understood that human activity on land impacts the environment within rivers, in turn affecting bacterial abundance and diversity (Van Rossum et al. 2015; Chen et al. 2018; Phiri et al. 2020; Qiu et al. 2020). However, even though some water-borne viruses have important implications for human health, such as enteroviruses (Amvrosieva et al. 2001), hepatitis E virus (Sedyaningsih-Mamahit et al. 2002; Martolia et al. 2009) and norovirus (Jack et al. 2013; Sekwadi et al. 2018), we know little about how human land-use impacts viral abundance and diversity in rivers. A study of an agricultural river basin in Ontario, Canada, found that higher levels of human viruses and coliphages were associated with greater upstream human land development (Jones et al. 2017). Similarly, land use in Singapore was the main driver of viral community structure in reservoirs used for potable water supplies and recreational activities (Gu et al. 2018). However, because such comparisons involved different catchments that are likely to have contrasting viral communities, it may not be possible to isolate the effect of human activity on viral ecology. To date there has been no study directly comparing the viral ecology of river water flowing through different land-use types within the same river catchment.

New Zealand freshwater communities have been isolated since New Zealand split from Gondwanaland approximately eighty million years ago (Mortimer et al. 2019). Freshwater communities within New Zealand are also generally isolated from each other, with little opportunity for non-migratory species to colonize new water catchments (Burridge and Waters 2020). This is reflected in the evolution of freshwater plant, vertebrate and invertebrate species. For example, there are high levels of endemism within New Zealand non-migratory galaxiid fish, with many species are found only in one water catchment (Dunn et al. 2018; Burridge and Waters 2020). It might therefore be expected that the freshwater communities of New Zealand would similarly contain many highly divergent viruses and locally unique viruses that have co-evolved with their isolated hosts. In contrast, human activity has had a large impact on New Zealand freshwater communities, including run-off from intensive agriculture and urbanization and the introduction of invasive species such as rainbow trout (*Oncorhynchus mykiss*). It is likely that these changes would also have affected the viral community in the rivers, introducing viruses associated with plants and animals grown for food, as well as viruses that infect humans.

Very little is known about the viral ecology of New Zealand rivers, with research generally limited to targeted testing for known pathogens. Two river sites – the Waikato River in the North Island and the Oreti River in the South Island – that supply drinking water to urban populations have been screened for enteric viruses, with positive results in 97% of samples (Williamson et al. 2011). In the Manawatū region, the Manawatū and Pohangina rivers and Turitea creek have been screened for plant viruses, with three tombusviruses detected (Mukherjee 2011; Mukherjee et al. 2012). New variants of Tobacco mosaic virus and Tomato mosaic virus were also identified (Mukherjee 2011), and human polyomaviruses have been found in the Matai river in Nelson (Kirs et al. 2011). *Sclerotinia sclerotiorum* Hypovirulence-Associated Virus-1 (which infects a fungus often found in agricultural plants) was detected in a Christchurch river using metagenomic sequencing of DNA from sediment samples (Kraberger et al. 2013). To our knowledge, viral meta-transcriptomics has not yet been performed on a river system in New Zealand.

The core aim of this study was to compare the viral (particularly RNA virus) abundance and diversity between sites with differing human land use impact in a New Zealand river catchment and from this determine how virome ecology and evolution are shaped by human activity. Accordingly, six sites on the Manawatū River, North Island, were selected based on their differing land use types. Two sites were at the edge of the Ruahine forest park, containing water that has only flowed through pristine native forest (pristine sites, denoted P1 and P2). Two sites contain water that has flowed through farmland (farmland sites, at least 25 km for F1 and 50 km for F2). The final two sites have flowed first through pristine native forest, then farmland and finally urbanized areas (urban sites) – Feilding and Palmerston North (named U1 and U2, respectively). Water samples were taken at these sites and subjected to total RNA sequencing (i.e., meta-transcriptomics).

## 2. Methods

### 2.1 The Manawatū River

The Manawatū River is a 180 km river located in the North Island of New Zealand (Figure 1). Importantly, it flows through three very different land-use types, allowing a direct comparison between them. The river begins in the Ruahine forest park that encompasses the Ruahine mountain range (Department of Conservation 2021b). The park is dominated by native vegetation, including podocarp forest at lower altitudes and sub-alpine shrubland and tussock grasslands at higher altitudes. Between the 1800s and 1970s there was considerable forest clearing and logging, but since 1976 the area has been protected as a forest park with no farming or logging (Department of Conservation 2021a). There is little to no human habitation or activity in these ranges, with the exception of recreational hikers, hunters and rangers. A variety of endemic New Zealand animals inhabit these ranges, including parrots (kakariki *Cyanoramphus novaezelandiae*, kaka *Nestor meridionalis*), ducks (whio *Hymenolaimus malacorhynchos*), New Zealand long-tailed bats (*Chalinolobus tuberculatus*) and large carnivorous land snails (*Powelliphanta marchanti*). Introduced pest species are also found there, including red deer (*Cervus elaphus*), feral pigs (*Sus scrofa)* and goats (*Capra hircus)*, brush tailed possums (*Trichosurus vulpecula*), stoats (*Mustela erminea*) and rainbow trout (*Oncorhynchus mykiss*) (Department of Conservation 2021b).

**Figure 1.**
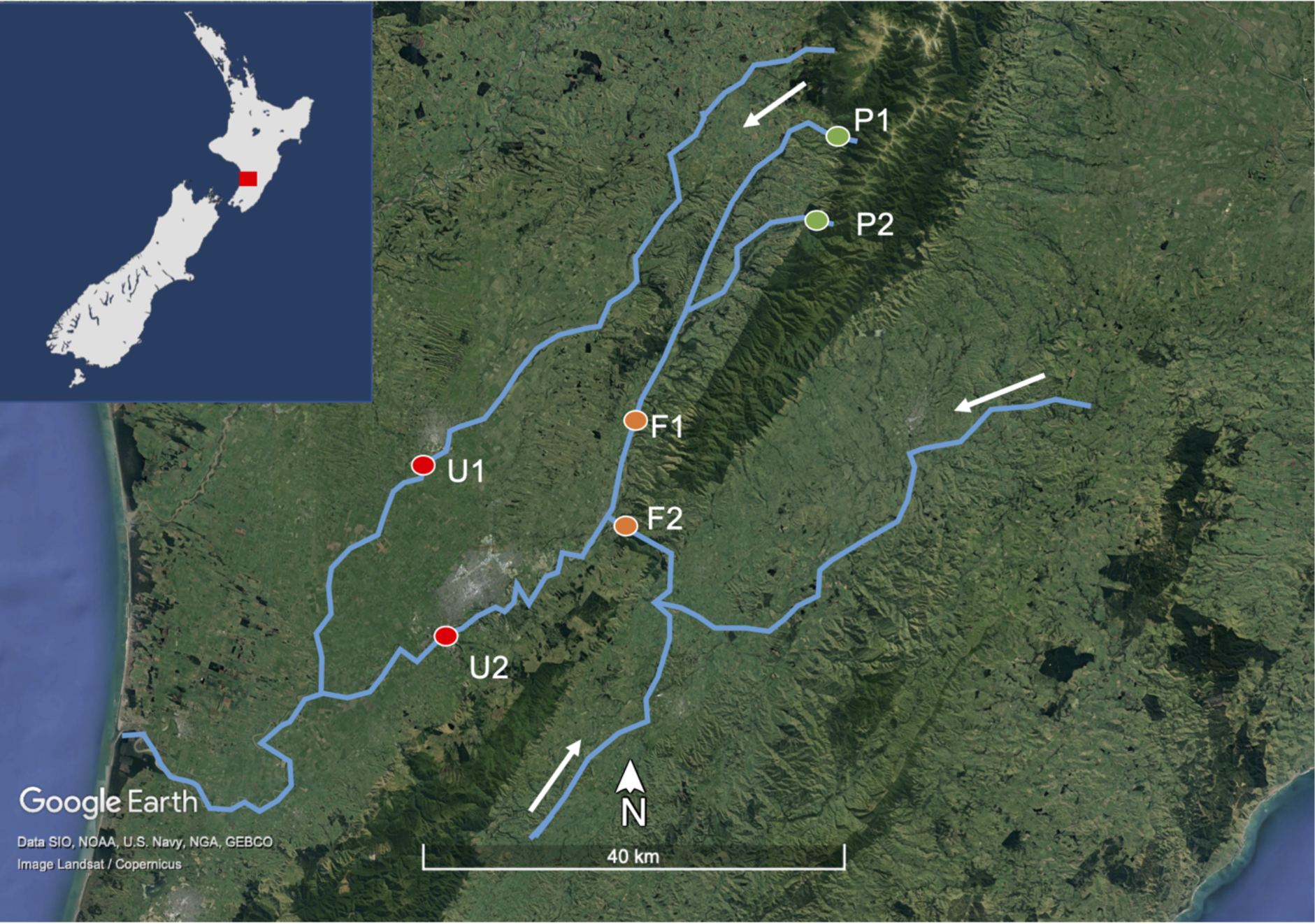
Map of the Manawatū River catchment in blue and the six sampling sites as coloured circles, with the inset showing the river catchment location on the North Island of New Zealand as a red square. P1 and P2 are ‘pristine’ sites having only flowed through native bush (seen as dark green areas on the satellite image). F1 and F2 are ‘farming’ sites flowing through intensive agricultural land (light green on the satellite image). U1 and U2 are ‘urban’ sites, flowing through two urban areas – Feilding and Palmerston North (grey on the satellite image). The water flows from the east and south towards the sea on the west coast, as indicated by the white arrows. Satellite image created using Google Earth.

After flowing through the Ruahine Forest Park, the river passes through intensively farmed areas, primarily consisting of sheep, beef and dairy farming. This land use is known to impact the river, with the Manawatū region having among the highest nitrogen and phosphorus concentrations (nutrients associated with pastoral agriculture) in New Zealand (Roygard et al. 2012). It then flows through two urban centres: Feilding (town, population 17,050), and Palmerston North (city, population 81,500), before flowing out into the Tasman Sea. Both these urban centres discharge treated wastewater into the Manawatū River, and in some years this negatively impacts aquatic life through discharge of nutrients and a corresponding increase in periphyton cover (Hamill 2012).

### 2.2 Sample collection

Two-litre (L) water samples were collected at each of the six sites in the Manawatū River catchment (Figure 1). For consistency, all samples were collected on the same day (13^th^ July 2019). At each site, 1 L was collected from the water’s edge and 1 L was collected 1.2 metres from the bank in the main flow of the river using a sampling pole, with the aim of obtaining a representative sample of the river water. In the case of P2, which was less than 1.2 metres wide, the second sample was taken by hand in the main flow of the river. These samples were combined to obtain 2 L of water per site and 12 L in total. Once collected, the water samples were kept at approximately 4°C using icepacks until processing.

At each site, a separate 250 ml sample was collected from the main flow of the river to measure additional variables (temperature, salinity, conductivity, pH, total dissolved solids, turbidity). These were measured using a multiparameter tester (Waterproof PCSTestr 35, Thermo Scientific) and a Turbidimetre (2100P Turbidimetre, Hach). Temperature was measured on-site and the remaining variables were measured in the laboratory.

### 2.3 Sample processing and sequencing

Filtering of all samples was completed within thirty hours of sample collection. Samples with a large amount of silt (farming and urban sites) were first filtered through a glass fibre filter, of 47 mm diameter and 0.7 μm pore size (Microscience). All samples were then filtered through polyether sulphone (PES) membrane filters, of 47mm diameter and 0.2μm pore size (Microscience).

Samples were concentrated from 2 L to ∼100 µL in two steps using tangential ultra-filtration and ultra-centrifugation. The samples were first concentrated from 2 L down to 40 mL using the vivaflow 200 (Sartorius). The samples were then concentrated from 40 mL to ∼2 mL using vivaspin 20 ultrafiltration units (Sartorius), and further concentrated to ∼100 µL using vivaspin 2 ultrafiltration units (Sartorius). All units had polyether sulphone filters with a molecular weight cut-off of 10 kDa. Samples were stored at -80°C until nucleic acid extraction.

RNA and DNA were extracted from the concentrated water samples using the AllPrep® PowerViral® DNA/RNA Kit. DNA was removed by DNase digestion (Qiagen RNase-Free DNase I Set), then in the same column the RNA was concentrated to 15 µL using the MN NucleoSpin RNA Clean-up XS (Macherey-Nagel). The same process was conducted on 2 x 200 µL of sterile water to create two blank control libraries. cDNA libraries were prepared using the SMARTer® Universal Low Input RNA Kit for Sequencing (Takara Bio), without rRNA depletion. Libraries were sequenced on the Illumina Novaseq platform (150bp, paired end sequencing). The corresponding sequencing data have been deposited in the Sequence Read Archive (SRA) under accession numbers SRR17234948-53. The trimmed alignment fasta files used to infer the phylogenetic trees are available at https://github.com/RKFrench/Viral-Diversity-NZ-River.

### 2.4 Quality control, assembly and virus identification

TruSeq3 adapters were trimmed using Trimmomatic (0.38) (Bolger et al. 2014). Bases below a quality score of five were trimmed with a sliding window approach (window size of four). Bases at the beginning and end of the reads were similarly excluded if below a quality of three. SMART adapters were trimmed using bbduk in BBtools (bbmap 37.98) (Bushnell 2018). Sequences below an average quality of ten were removed.

Sequence reads were assembled *de novo* using Trinity (2.5.1) (Grabherr et al. 2011), with a kmer size of 32 and a minimum contig length of 300. BLASTN (BLAST+ 2.9.0) and Diamond BLASTX (Diamond 0.9.32) were used to identify viruses by comparing the assembled contigs to the NCBI nucleotide database (nt) and non-redundant protein database (nr) (Camacho et al. 2009; Buchfink et al. 2021). Contigs with hits to viruses were retained.

To avoid false-positives, sequence similarity cut-off values of 1E-5 and 1E-10 e-value were used for the nt and nr databases, respectively. Virus abundances were estimated using RSEM (1.3.0), allowing us to determine the expected count according to the Expectation-Maximization algorithm for each contig (Li and Dewey 2011). This was expressed as the percentage of the total number of reads in each library. Eukaryotic and prokaryotic diversity was characterized using CCMetagen (v 1.2.4) and the NCBI nucleotide database (nt) (Clausen et al. 2018; Marcelino et al. 2020).

### 2.5 Evolutionary and ecological analysis

Using the nucleotide sequences identified as viral replication proteins (i.e., as identified by BLAST), the getorf program from EMBOSS (6.6.0) was used to find and extract open reading frames and translate them into amino acid sequences using the standard genetic code with a minimum size of 100 amino acids (Rice et al. 2000). Amino acid sequences were aligned using the E-INS-i algorithm in MAFFT (7.402) (Katoh and Standley 2013) and trimmed using Trimal, (1.4.1) (Capella-Gutiérrez et al. 2009) with a gap threshold of 0.9 and at least 20% of the sequence conserved. See Supplementary Table S1 for more details. Maximum likelihood phylogenetic trees for each virus family were then estimated using IQ-TREE (1.6.12) (Nguyen et al. 2015), with the best fit substitution model determined by the program and employing 1000 bootstrap replications to assess node robustness. Any sequences with >95% amino acid similarity to each other or known species were assumed to represent the same virus species, with only one representative of each then included in the phylogenetic analysis. All novel viruses identified were given names that include the word ‘flumine’ (Latin for ‘of the river’) to convey where the virus was found.

APE (5.4) and ggtree (2.4.1) were used to visualize the phylogenetic trees and produce figures (Paradis and Schliep 2019; Yu 2020). Alpha diversity (i.e., diversity within each sample) was analysed using richness (number of viral families), Shannon index and the Shannon effective number of species (ENS). The Shannon index reflects the number of taxa and the evenness of the taxa abundances. The Shannon ENS is the effective number of taxa present in the community if the abundances were equal (Hill 1973). The beta diversity (i.e., diversity across land-use types) was analysed using a principal co-ordinate analysis with a Bray–Curtis dissimilarity matrix, presented as an ordination plot. Alpha and Beta diversity analyses were conducted using Phyloseq (v1.34.0) in R (v 4.0.5) (McMurdie and Holmes 2013; R Core Team 2021). Other graphs were generated using ggplot2 (Wickham 2016) and venneuler (Wilkinson 2011).

### 2.6 Identifying possible reagent contamination

Any virus found in the blank negative control libraries (i.e., a sterile water and reagent mix) was assumed to have resulted from contamination likely associated with laboratory reagents. Accordingly, these viruses were removed from the river sample libraries and excluded from all analyses. Additionally, any viruses that fell into the same clades as those found in blank libraries were conservatively assumed to be contaminants (Porter et al. 2021) and similarly removed. These included six circo-like viruses in the *Circoviridae* (single-strand DNA viruses) and 19 tombus-like viruses from the *Tombusviridae* (single-strand, positive-sense RNA viruses).

## 3. Results

We characterized the RNA viromes from six Manawatū River water samples using total RNA sequencing.

### 3.1 Water measurements

Our water measurements indicated that the two pristine sites had a different abiotic environment from the farming and urban sites, while the farming and urban sites were similar to each other (Figure 2). Specifically, the pristine sites had lower salinity, pH, total dissolved solids, turbidity, conductivity and temperature than the farming and urban sites. The farming and urban sites also had a larger variation between sites, with the exception of pH and temperature.

**Figure 2.**
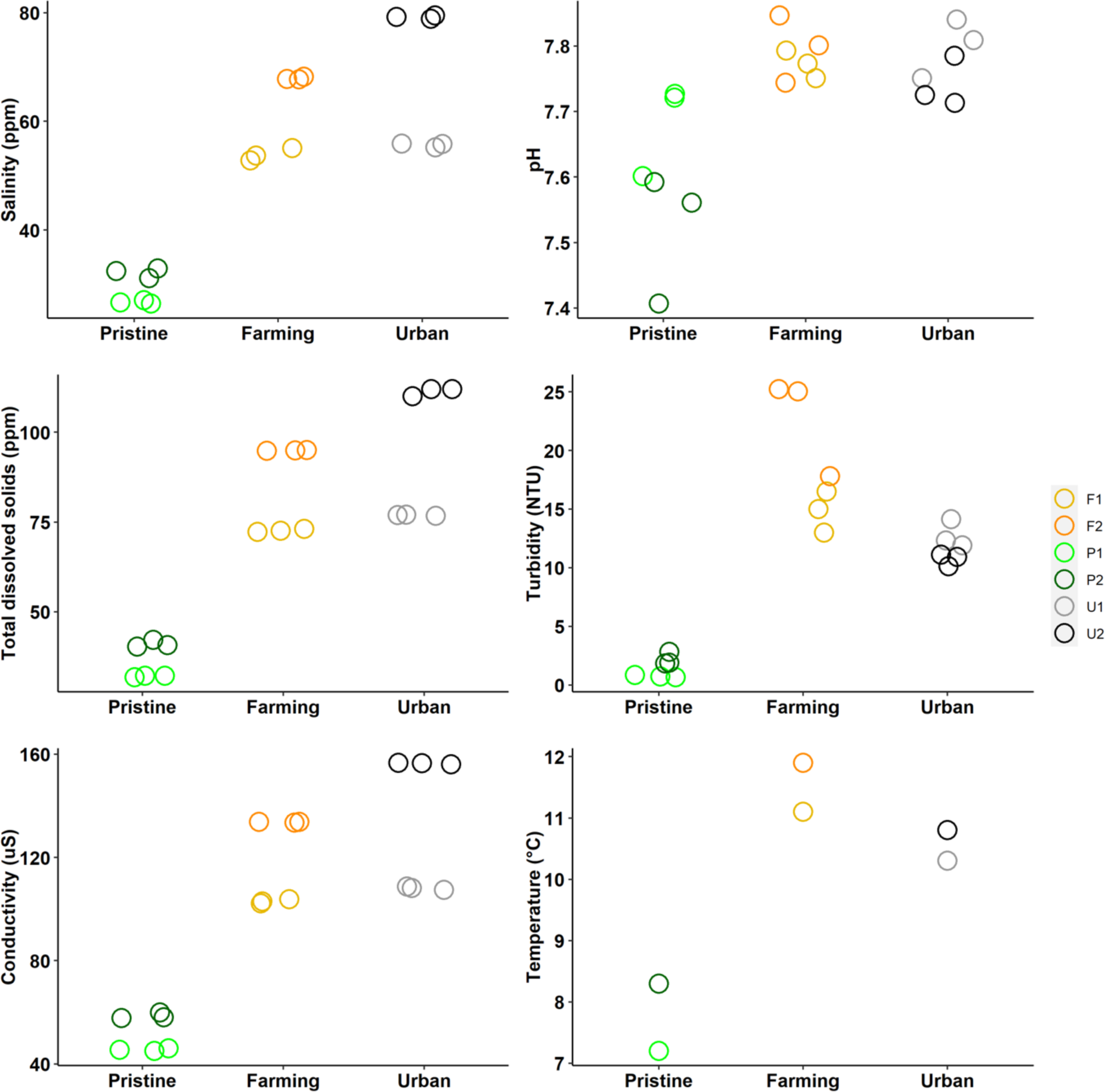
Measurements of salinity, pH, total dissolved solids, turbidity, conductivity and temperature at the six different sites on the Manawatū River, grouped into the different site types (pristine, farming and urban). For all measurements except temperature there are three values per sampling site which have been jittered to reduce overplotting.

### 3.2 Virus identification

The six sequencing libraries generated had an average of 90 million reads per library, and on average 0.7% of these were derived from viruses. This is within the usual range found in faecal samples, cloacal swabs and invertebrate tissue, but higher than commonly observed in vertebrates (Zhang et al. 2018; Campbell et al. 2020; Le Lay et al. 2020; Mahar et al. 2020; Wille et al. 2020; Wille et al. 2021) in studies using similar metagenomic techniques. However, it was lower than observed in urban streams in Ecuador (Guerrero-Latorre et al. 2018). P2 and U2 had the highest number of reads and the highest number of viral reads (Figure 3). Notably, the two pristine sites had the highest percentage of viral reads, at 1.31 and 1.35%. F1, F2 and U1 all had lower total reads, total viral reads and percentage viral reads. Analysis of eukaryotic and prokaryotic diversity showed that all samples primarily consisted of bacteria (accounting for 71-86% of assembled contigs) followed by eukaryotes (5-19%).

**Figure 3.**
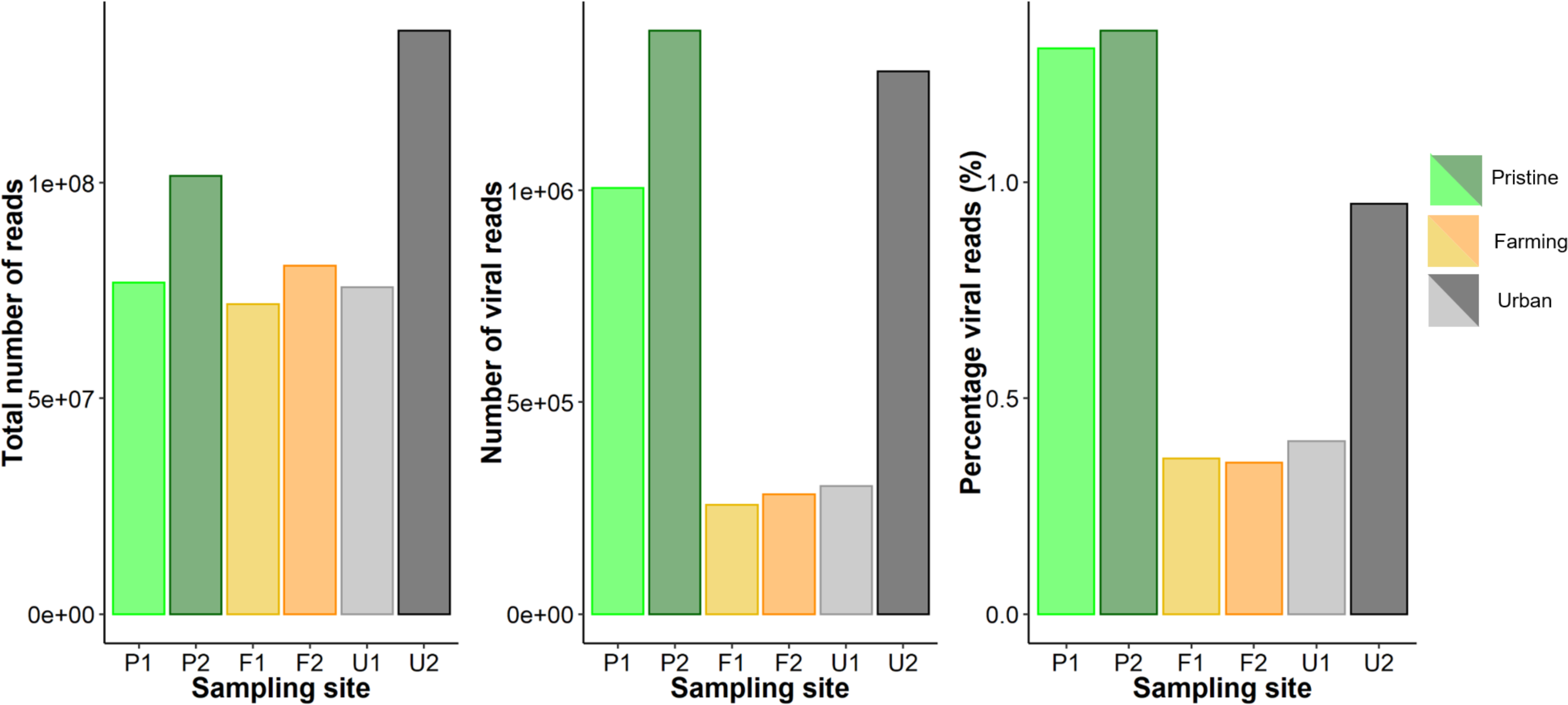
Read counts and the percentage of viral reads (%) for libraries from six sites on the Manawatū River.

In total, we identified 504 putative virus species from 27 viral families, of which 491 (97%) were novel using a cut-off of 95% amino acid similarity, primarily in replication-associated proteins (although these await formal verification by the International Committee on Taxonomy of Viruses). These included multiple members of the *Nodaviridae* (n=74 novel viruses), *Tombusviridae* (n=64) and *Dicistroviridae* (n=61). If a more conservative <90% amino acid sequence similarity is used to define a novel virus species, then the samples analysed here contain 470 novel viruses. We also detected previously described viruses (i.e., with more than 95% amino acid similarity to viruses detected previously), including *Sclerotinia sclerotiorum* hypovirulence associated DNA virus 1 (*Genomoviridae*), White clover mosaic virus (*Alphaflexiviridae)*, Rhopalosiphum padi virus (*Dicistroviridae),* Norway luteo-like virus 4 (*Luteoviridae),* Carnation Italian ringspot virus *(Tombusviridae)* and Pepper mild mottle virus (*Virgaviridae)*. Importantly, we identified at least 63 virus species from the *Astroviridae, Birnaviridae, Parvoviridae* and *Picornaviridae* that may infect vertebrates. No likely human viruses were detected. Below we describe, in more detail, those families with high virus diversity in our study and those containing viruses that may infect vertebrates.

### 3.3 High Phylogenetic Diversity Families

#### 3.3.1 Tombusviridae

We identified a high diversity and abundance of novel *Tombusviridae,* a family of single-strand positive-sense RNA viruses that infect plants (Sit and Lommel 2015). Of the viruses identified 18 fell into the subfamily *Procedovirinae*, found in all land-use types. Of these, one virus in U2 clustered within the genus *Betacarmovirus* and was most closely related to Cardamine chlorotic fleck virus (Skotnicki *et al*. 1993), although with only 45% amino acid similarity. Another, found in U2 and F2, belonged to the genus *Gammacarmovirus* and is most closely related to Melon necrotic spot virus with 71-73% amino acid similarity (Riviere and Rochon 1990). We also found Carnation Italian Ringspot virus (genus *Tombusvirus*) at both farming sites, with 98-99% amino acid similarity. The *Procedovirinae* also contain a clade of closely related novel tombus-like viruses that appear basal to any currently described genera and were found at the pristine sites. Another clade was found in the urban and farming sites and appears to fall within the subfamily *Regressovirinae*.

#### 3.3.2 Dicistroviridae

We similarly found a high diversity of viruses belonging to the *Dicistroviridae*, a family of single-strand positive-sense RNA viruses commonly associated with arthropods (Valles et al. 2017). Thirty-nine viruses were part of a highly divergent clade that fell basal to the three currently recognized genera, to which it exhibited less than 50% amino acid similarity. There were also 15 newly identified viruses that fell into the genus *Cripavirus* found across all the land-use types. In addition, we detected two previously described cripaviruses - Rhopalosiphum padi virus (97-100% amino acid similarity) and Cricket paralysis virus (96-97%), both only at the urban site U2.

#### 3.3.3 Nodaviridae

We identified 13 novel viruses from the genus *Alphanodavirus* of the *Nodaviridae* (single-strand positive-sense RNA viruses). This included Black beetle virus (*Alphanodavirus;* 95-98% amino acid similarity) in all farming and urban sites, but not at the pristine sites.

Although we did not find any viruses belonging to the genus *Betanodavirus*, the only other genus of *Nodaviridae*, we did identify 61 other novel viruses from a divergent clade that fell outside of the *Alphanodavirus* and *Betanodavirus* genera. Many of these were most closely related to Barns Ness serrated wrack noda-like virus 2 isolated from marine algae (Waldron et al. 2018), although with less than 50% amino acid similarity.

### 3.4 Vertebrate-infecting families

#### 3.4.1 Astroviridae

The *Astroviridae* are a family of single-stranded positive-sense RNA viruses that infect mammals and birds (Lukashov and Goudsmit 2002). We identified 28 novel astroviruses, including three new species within the *Bastrovirus* clade that were found in the farming and urban sites (Figure 4). These three viruses were most closely related to a bastrovirus found in sewage in Brazil (Dos Anjos et al. 2017), although with only 57-67% amino acid similarity. Twenty-four novel viruses fell into a divergent clade outside of the genus *Avastrovirus* and the *Bastrovirus* clade: these were most closely related to ‘*Astroviridae* sp.’ viruses found in metagenomic studies of grassland soil in California, USA (Starr et al. 2019). The viruses found on our study that fell into this clade (denoted Flumine astrovirus 1-24) all had less than 62% amino acid similarity with the soil viruses. Notably, they were found across all our river sites, with a higher diversity at the farming and urban sites (n=11 and 21 species, respectively) than the pristine sites (n=4). However, flumine astrovirus 3 found at a pristine site had the highest abundance, representing 0.04% of the total reads in the library.

**Figure 4.**
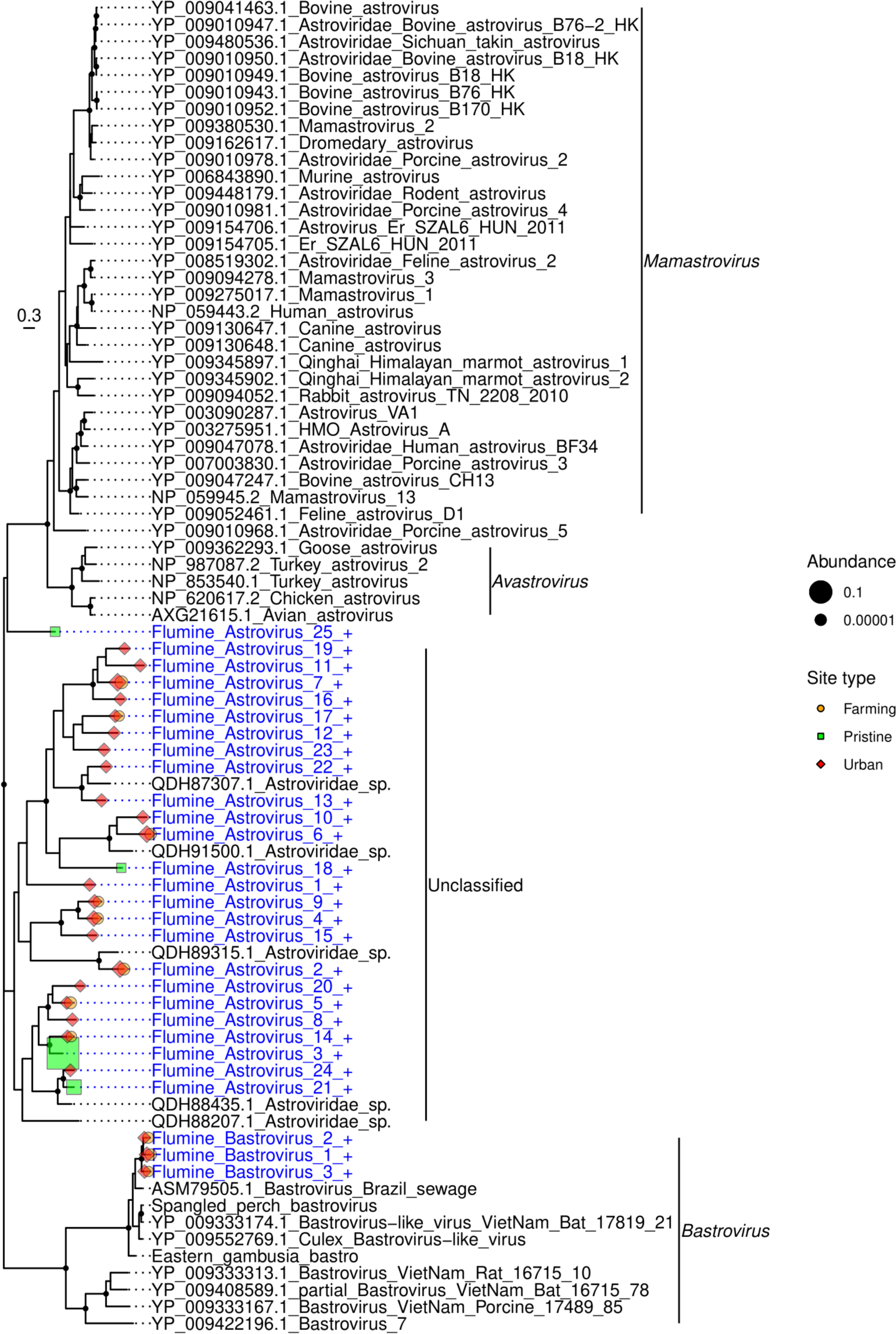
Phylogeny of the *Astroviridae* based on the non-structural polyprotein sequence (alignment length of 733 amino acids). Viruses obtained in this study are shown in blue and have a ’+’ after their name. Related viruses are shown in black. Virus abundance is expressed as the percentage of the total number of reads and represented by the size of each coloured symbol. The colour of each symbol refers to the site type. Black circles on nodes show bootstrap support values greater than 90%. Branches are scaled according to the number of amino acid substitutions per site, shown in the scale bar. The tree is midpoint rooted.

#### 3.4.2 Birnaviridae

The *Birnaviridae* are a family of double-stranded RNA viruses that infect fish, birds and insects. We identified one virus from this family at a pristine site (P1), that was most closely related (although with only 30% amino acid similarity) to blotched snakehead virus (Da Costa et al. 2003) and Lates calcarifer birnavirus (Chen et al. 2019), both of which are associated with fish.

#### 3.4.3 Parvoviridae

The *Parvoviridae* are a family of small double-stranded DNA viruses that infect both vertebrates and invertebrates. We identified 21 parvoviruses: all were present at the Feilding urban site (U1), with one (Flumine parvovirus 17) found at both P1 and U1. Most (n=18) of these viruses fell into a distinct clade, separate from other previously described subfamily and genera (Figure 5). This clade had less than 18% amino acid similarity with other parvoviruses, but up to 81% similarity (average of 41%) with each other. A new subfamily *Hamaparvovirinae* was created in 2020 that exhibits less than 20% amino acid sequence identity with other parvoviruses (Pénzes et al. 2020). This is comparable to the level of sequence similarity observed in the novel clade identified here, indicating this may also represent a new subfamily that we have tentatively called the *Flumenparvovirinae*. The remaining three viruses fell into the subfamily *Densovirinae*. These viruses most likely infect invertebrates as they were most closely related to Planococcus citri densovirus (Thao et al. 2001) (76% amino acid identity) and Blattella germanica densovirus (Kapelinskaya et al. 2011) (55-61%) that infect mealy bugs and cockroaches, respectively.

**Figure 5.**
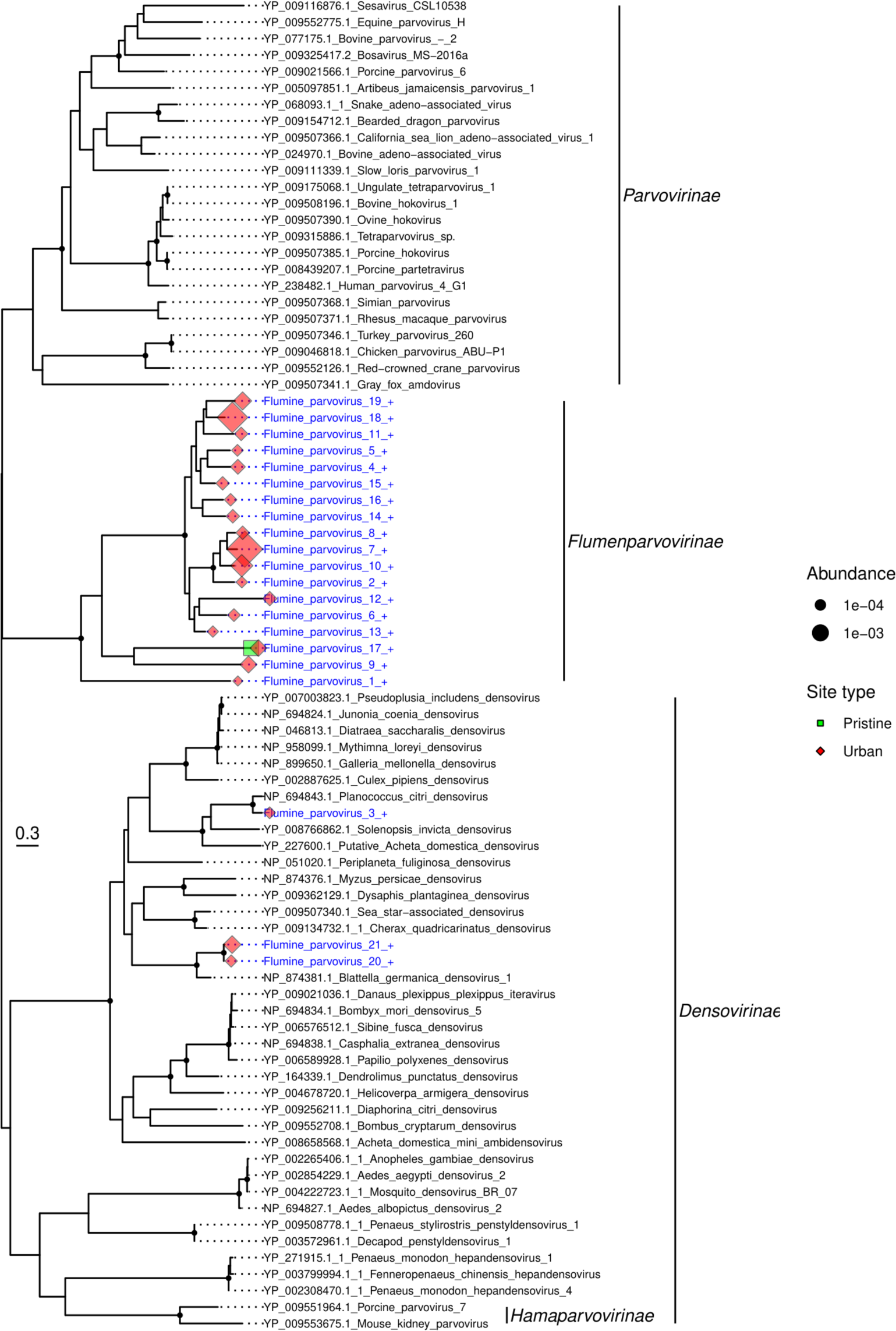
Phylogeny of the *Parvoviridae* based on the non-structural protein sequence (alignment length of 452 amino acids). Viruses obtained in this study are shown in blue and have a ’+’ after their name. Related viruses are in black. Virus abundance is expressed as the percentage of the total number of reads and represented by the size of each coloured symbol. The colour of each symbol refers to the site type. Black circles on nodes show bootstrap support greater than 90%. Branches are scaled according to the number of amino acid substitutions per site, shown in the scale bar. The tree is midpoint rooted.

#### 3.4.4 Picornaviridae

The *Picornaviridae* are a large family of single-stranded RNA viruses that infect both vertebrates and invertebrates. We identified 15 novel picornaviruses, most (n=11) from the urban sites, although they were also identified at both farming and pristine sites. The pristine sites had the lowest diversity of picornaviruses (n=2), but the highest abundance. Eight of the picornaviruses fell into a clade with fur seal picorna-like virus (Krumbholz et al. 2017) and Ampivirus A1 associated with the smooth newt *Lissotriton vulgaris* (Reuter et al. 2015). This clade is most closely related to a cluster of genera *Tremovirus, Harkavirus* and *Hepatovirus* found in birds and mammals.

#### 3.4.5 Genomoviridae

The *Genomoviridae* are a family of single-stranded DNA viruses. Although they are commonly associated with fungi, they have also identified them in mammals and birds (Varsani and Krupovic 2021). With the exception of one virus species (Flumine genomovirus 1), members of the *Genomoviridae* were only found at the urban sites. Nine of the 14 viruses were most closely related to viruses found in sewage or faeces. Interestingly, a number of these viruses are closely related to viruses previously documented in New Zealand. Flumine genomovirus 11 fell into a clade of five sewage-derived gemycircularviruses, exhibiting 85% amino acid similarity to its closest match Sewage-associated gemycircularvirus-7a sampled from a sewage oxidation pond in Christchurch, New Zealand (Kraberger et al. 2015). Similarly, Flumine genomovirus 8 and 9 share 85% and 86 % amino acid similarity, respectively, with Sewage associated gemycircularvirus 3 also isolated from a New Zealand oxidation pond (Kraberger et al. 2015), while Flumine genomovirus 12, 4 and 6 share 87%, 76% and 82% amino acid similarity, respectively, with faeces-associated gemycircularvirus 21 isolated from llama faeces in New Zealand (Steel et al. 2016). Finally, the single *Genomoviridae* virus found in the pristine sites (Flumine genomovirus 1) was most closely related (81% amino acid similarity) to a virus isolated from minnow tissue in the USA.

### 3.5 Host relationships and patterns of virus diversity

We next characterized each viral family according to their usual assigned host (as identified in previous studies) and used this to visualize patterns of virus abundance and diversity (Figure 6). This revealed that the two farming sites had a lower viral abundance across all host types, while the two pristine sites had a higher abundance of viruses with animal, plant/fungi (such as the *Astroviridae* and *Tombusviridae*) and unknown hosts. In turn, U1 had a very high abundance of prokaryote-infecting viruses but a lower abundance of all other virus types (Figure 6). Virus abundance was highest for those viral families where the host was unknown, accounting for 2.35% of reads. Plant-infecting viruses were the next-most abundant (1%), followed by animal-infecting viruses (0.56%). At the family level, we found a high diversity of plant- and animal-infecting viruses (18 and 17 families, respectively).

**Figure 6.**
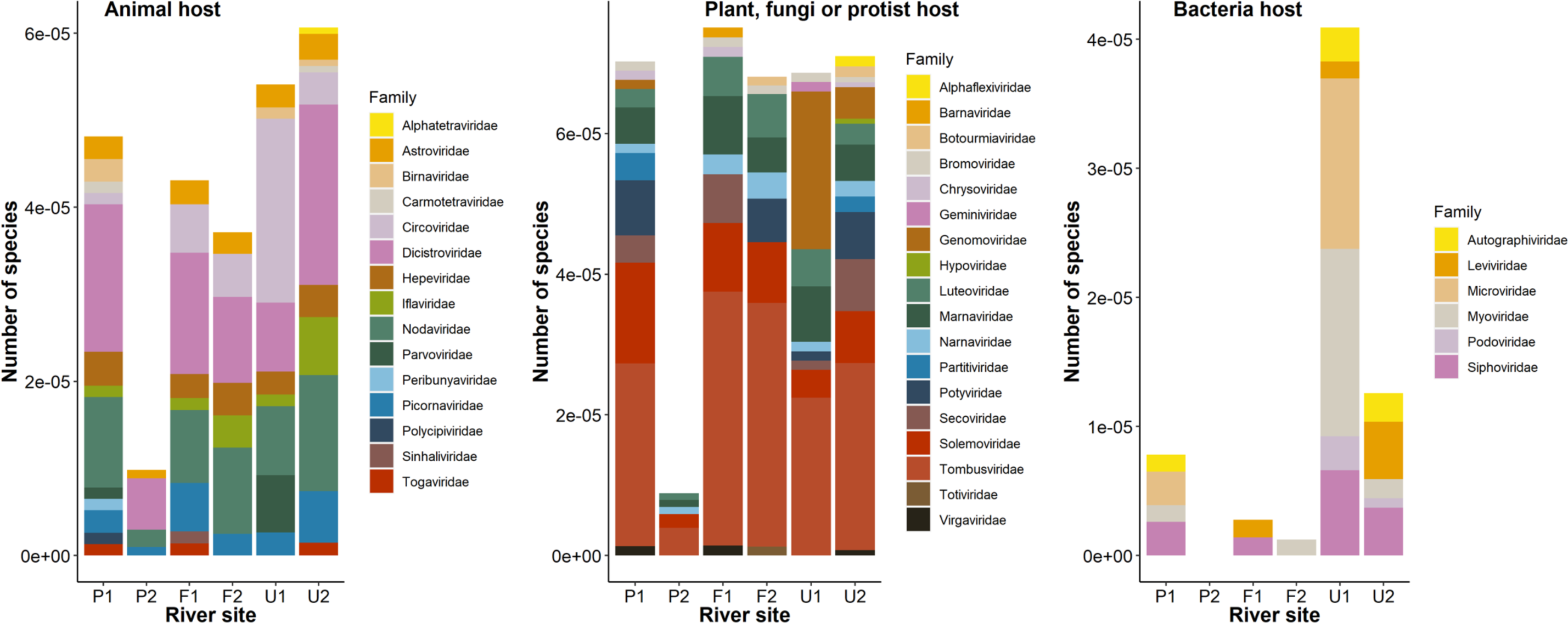
Virus abundance (as a percentage of the total number of reads) of viral families from each site on the Manawatū River. Virus families are divided into each panel depending on their usual host - animal, plant/fungi, prokaryote and unknown. Unclassified viruses did not have a classification according to the current NCBI taxonomy.

There was a surprisingly low abundance and diversity of known prokaryote infecting viruses: these were the least abundant (0.096%), and the least diverse (six families) group. All sites had a proportionally high abundance of *Nodaviridae, Tombusviridae* and ‘unclassified *Picornavirales’* compared to other viral families.

### 3.6 Alpha and beta virus diversity

Sites P1 and U2 had the highest richness, manifesting as 38 and 40 virus families, respectively, while P2 had the lowest at 16 (Table 1). U1 had higher Shannon and Shannon ENS values than any other site, but only the third highest richness. The two pristine sites had very different levels of diversity, with P1 having much higher richness (with more than twice the number of viral families), Shannon and Shannon ENS than P2. The two farming sites had a similar richness, Shannon and Shannon ENS to each other, and the two urban sites were also similar with respect to richness (with U1 having 20% fewer virus families than U2).

**Table 1.** The richness, Shannon, and Shannon effective number of species (ENS) values for each site on the Manawatū River. These were calculated at a family level and include unclassified virus groups such as ‘unclassified Riboviria’.

|  | Richness | Shannon | Shannon ENS |
| --- | --- | --- | --- |
| P1 | 38 | 2.13 | 8.39 |
| P2 | 16 | 1.88 | 6.54 |
| F1 | 28 | 2.21 | 9.07 |
| F2 | 24 | 2.11 | 8.28 |
| U1 | 32 | 2.52 | 12.45 |
| U2 | 40 | 2.00 | 7.39 |

At the level of virus family, we used principal co-ordinate analysis to examine the differences between viral communities, examining ‘intra-type’ (i.e., comparing sites with the same land-use type) and ‘inter-type’ (comparing sites across different land-use types) differences (Figure 7). Accordingly, the pristine and farming sites had high intra-type similarity and low inter-type similarity, such that their viral communities were more similar within a land-use type than to viral communities from different land-use types (Figure 7). The two pristine sites displayed the most similar viral communities. In contrast, the two urban sites differed markedly from each other and all other sites and hence had a very different viral community, both from each other and to that at any other site (i.e., both intra- and inter-type). The urban site U2 (Palmerston North) was closer to F1 and F2 than to U1, and hence had a viral community that was more similar to the farming sites than to the other urban site. At the viral species level, the urban sites had a higher diversity (n=327 species) than the farming (n=150) and pristine sites (n=119). There were many more viruses shared between the urban and farming sites (n=76) than between the pristine and farming or urban sites (n=24). Finally, only eight of 504 species were found in all land-use types, indicative of a relatively high level of local differentiation.

**Figure 7.**
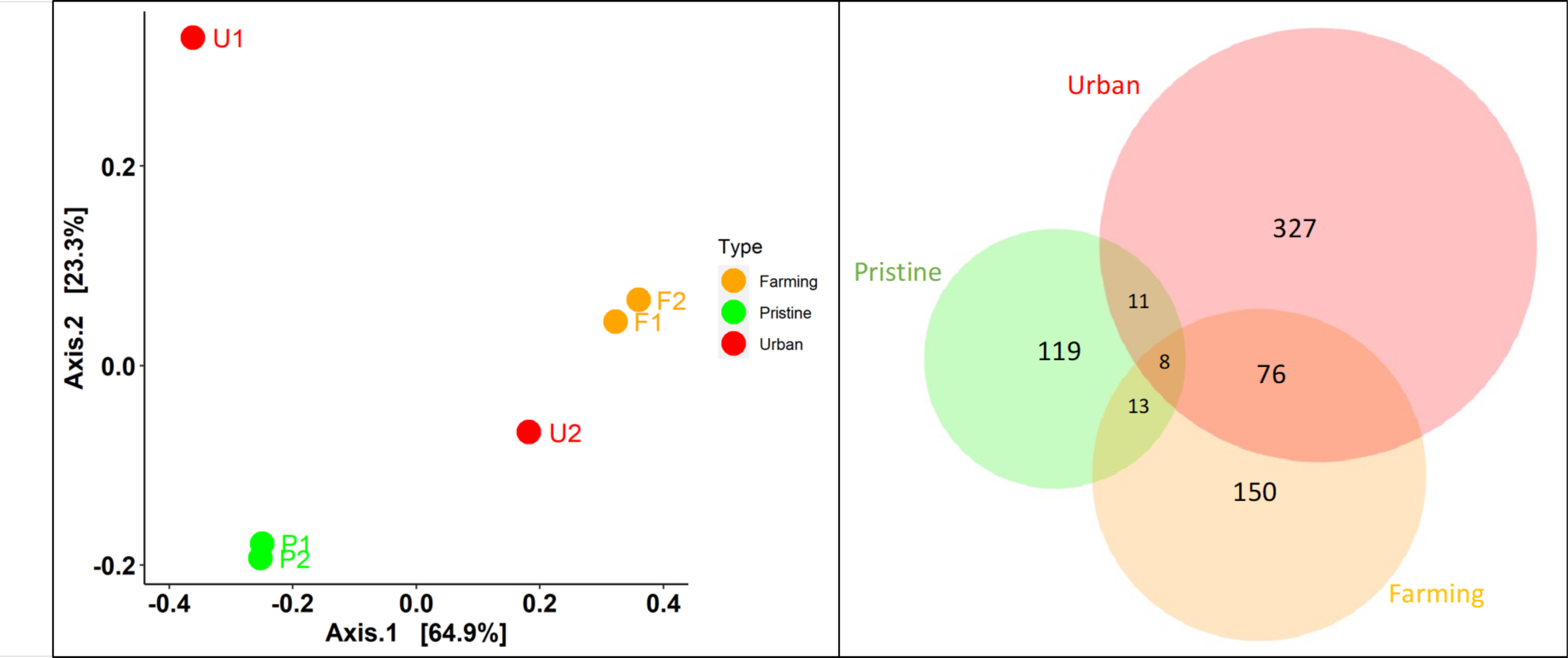
Viral community similarities and differences at the taxonomic family level (left) and species level (right). Left - A Principal Coordinates Analysis (PCoA) applying the Bray– Curtis dissimilarity matrix for viral abundance and virus family diversity, showing the relative similarity/differences in viral community between sites with differing land-use types – pristine (green), farming (orange), and urban (red). Points closer to one another are more similar in virome structure than those further away. Right – a Venn diagram showing the number of virus species shared between the three land-use types (with an amino acid similarity >95%).

## 4. Discussion

Using a mNGS approach we identified a high diversity of novel and highly divergent viruses in a single New Zealand river system and revealed differences of virome composition in the river between sampling sites associated with different land-use types. In total, we observed 504 putative virus species, of which 491 (97%) were potentially novel and including a new subfamily within the *Parvoviridae*. Notably, there were considerable differences in the viral community structure between the land-use types. In particular, the two pristine sites had a higher abundance of viruses that infect animals, plants and fungi, while at the viral species level, the urban sites had the highest virome diversity. In addition, there were many more viruses shared between the urban and farming sites (n=76) than between the pristine and farming or urban sites (n=24).

The abiotic environment within the Manawatū River system differed considerably between sampling sites that had different land-use types, with the pristine sites having lower measurements of all variables. Total dissolved solids (TDS), salinity, conductivity and turbidity represent different ways of measuring the presence of dissolved solids and ions in the water, and generally the lower they are then the higher the water quality (Davies-Colley 2013). Increases in total dissolved solids can adversely affect plants and animals in freshwater environments due to changes in osmotic conditions, making this an important indicator of the health of the freshwater ecosystem (Chapman et al. 2000). These measures are all elevated in effluents, and their presence in freshwater can therefore indicate contamination from fertilizers, urban run-off, and animal and human waste (Chapman et al. 2000; de Sousa et al. 2014). They can also naturally be elevated due to changes in climate and differing geology, which impacts the amount of dissolved substance from the weathering of rocks (Davies-Colley 2013). However, as the sites in this study were all in the same river catchment and within 50km of each other, the climate and geology are unlikely to be markedly different. Water temperature was also lower in the pristine sites than in the farming and urban sites, likely due to the pristine sites being at higher altitude, closer to the Ruahine Ranges (a source of snow melt), and also the thicker vegetation cover blocking solar radiation. The pristine sites had a pH closer to neutral, which may be related to differences in dissolved solids from the surrounding soils (Baldisserotto 2011). All these indicators show that the riverine environment was very different between the pristine and farming/urban sites, but not substantially different between the farming and urban sites.

A key observation of our study was the high diversity of viruses in the Manawatū River system, with an average of 30 virus families detected per site. Indeed, we found a very high diversity of novel viruses, and only a small number (n=13) described previously, again indicating that only a tiny fraction of the virosphere has been described to date. Environments such as freshwater have received relatively little viral metagenomic research, although our study suggests that they may have highly diverse viromes. In particular, we show that isolated New Zealand freshwater environments can harbour many novel and highly divergent viruses. Perhaps of most note, we have identified a novel clade of parvoviruses that was phylogenetically distinct from other previously described subfamily and genera, and which may be a new subfamily of the *Parvoviridae*. This may in part reflect New Zealand’s distinctive ecological history, including high levels of endemic species (Walker et al. 2021). These novel viruses may therefore be naturally present in this water catchment and reflect the ‘undisturbed’ diversity of the river, rather than the result of human land-use. A high diversity of bacteria have also been found in New Zealand freshwater (Phiri et al. 2021). Indeed, it is notable that the pristine sites had the highest abundance of viruses that infect animals, plants and fungi, and had little overlap (in terms of species shared) with the farming and urban sites (Figures 6 and 7). This may reflect a higher abundance of native flora and fauna in these pristine sections of the river that are surrounded by native forest. The two pristine sites had very different levels of diversity, with P1 having more than twice the number of viral families than P2, despite P2 having a higher number of viral reads. This may be because the P2 stream was smaller in volume with a smaller catchment, with less opportunity for a high diversity of terrestrial viruses to enter the river from the surrounding land.

Virus families that infect fungi, plants and algae were found in particularly high abundance, with the highest being the *Tombusviridae* associated with plants. This presumably reflects a high abundance of plant and algae matter in the water, including both aquatic plants living in the river and terrestrial plants from the surrounding land. However, there was a surprisingly low abundance and diversity of viruses associated with prokaryotes. This may be because we filtered out bacteria prior to RNA extraction (using PES membrane filters of 0.2μm pore size) which would also have removed any lysogenic phages. However, a previous study of urban streams in Ecuador using a similar filtering system found that prokaryote infecting viruses had the highest abundance, followed by plant viruses (Guerrero-Latorre et al. 2018). Interestingly, these Ecuadorian streams are known to be contaminated with untreated sewage (Guerrero-Latorr*e et al. 2*018), which may increase bacterial load (and therefore phage abundance). It is also possible that the high abundance of divergent viruses belonging to families in which the host is unknown (Figure 6) are in fact phage.

Metagenomics frequently identifies highly divergent viruses in environmental samples and in most such studies the host is unknown. We observed a similar pattern. The novel viruses identified often clustered into clades that fell in basal positions on family-level phylogenies and were most closely related to viral species found in other metagenomic studies (Zhang et al. 2018; Starr et al. 2019). As a case in point, many of the *Tombusviridae*-like viruses were highly divergent and fell into the ‘tombus-like’ virus clade. This is similar to the findings of Wolf et al. (2020), who identified 199 tombus-like viruses in seawater from the Yangshan Deep-Water Harbour, Shanghai, China. We also found many divergent viruses from the *Astroviridae* most closely related to viruses found using metagenomics from grassland soil, California, USA (Starr et al. 2019). As the *Astroviridae* is routinely associated with vertebrates it is surprising to find these viruses in soil and water. Hence, it is likely that these were shed from a vertebrate host although this clearly needs to be studied in greater detail.

As the sampling sites were all from the same river catchment, in some cases the water from one site would flow to other sampling sites downstream. For example, water from P1 and P2 would flow through F1 and U2. However, there did not appear to be a trend in the number of reads and number of viral hits when comparing upstream and downstream sites (Figure 3). In particular, very few viruses (n=17) were found at both the pristine sites and the downstream farming and urban sites, suggesting that viruses were generally not being carried downstream for long distances. Notably, for the pristine and farming sites, there was considerably more overlap in viral community at sites of the same land-use type: P1 and P2 were very similar to each other, as were and F1 and F2 (Figure 7). The exception to this was the urban site U2 (Palmerston North) that was more similar to the two farming sites than to U1 (Feilding). This difference appears to be driven by the high abundance and diversity of prokaryote infecting viruses in U1, and the lower abundance of animal- and plant-infecting viruses. Indeed, U1 was on a different tributary of the Manawatū River than the other sites (Figure 1), so the viral ecology in that part of the river may be different. Feilding (population 17,050) is also a much smaller urban centre than Palmerston North (population 81,500). In addition, while the sample sites from both urban sites were downstream of the urban area at the edge of the town/city, Feilding releases wastewater far downstream in a rural area, whereas Palmerston North releases wastewater upstream of our sampling site. However, although it might be expected that the presence of wastewater would increase the abundance of phage, Feilding had the higher phage abundance. Possibly changes in the bacterial diversity/structure in this urban site have resulted in differences in the phage diversity and structure.

Some novel viruses were found only in the sites adjacent to human land-use (i.e., farming and urban sites), including all the bastroviruses (*Astroviridae*), all of which were most closely related to a bastrovirus found in sewage (Dos Anjos et al. 2017). The pattern of presence/absence in our study and the link to sewage suggests the presence of these viruses are likely related to human land-use. Similarly, the presence of viruses from the *Genomoviridae* appears to be related to urban land-use: nine of the 14 viruses were most closely related to viruses found in sewage or faeces. Despite the generally large differences in viral community between the two urban sites, they both contained viruses from the *Genomoviridae*: 13 viral species were identified in U1 and four in U2 (including two viruses found in both sites), suggesting a link between human land-use and the presence of these viruses. Notably, however, we did not detect any human viruses. This contrasts with a PCR study that detected enteric viruses (adenovirus, norovirus, enterovirus, rotavirus, and hepatitis E virus) in two other rivers in New Zealand (Williamson et al. 2011), which may reflect the greater sensitivity of PCR assays specifically designed to detect these viruses. In general, more viruses were found in common between urban and farming sites. For example, 11 *Astroviridae*-like viruses were found in both urban and farming sites, but no *Astroviridae*-like viruses found in either of the pristine sites were identified in urban or farming sites. Hence, within viral families at the species level, the farming and urban sites had a more similar viral community to each other than to the pristine sites, which may be a result of human land-use adjacent to the river.

The only viruses we identified that were previously described were all found in the urban and farming sites, again indicative of an anthropogenic influence, including agriculture and introduced species. For example, we identified the Black beetle virus in all farming and urban sites. The host (*Heteronychus arator*) is a major invasive pasture pest species that was introduced from Africa in 1930s (Wilson et al. 2016), and the virus was first identified in 1975 in New Zealand (Longworth and Carey 1976). Similarly, at the Palmerston North urban site (U2) we detected Rhopalosiphum padi virus that infects the bird cherry-oat aphid, a pest of cereal crops that was first recorded in New Zealand in 1921 (Bulman et al. 2005).

Surprisingly, we found Norway luteo-like virus 4 with 96-100% amino acid identity in one farming and one urban site (F1 and U2), previously associated with the castor bean tick *Ixodes ricinus* (Pettersson et al. 2017). As this species of tick has not been described in New Zealand, this result suggest that this virus has a wider host range than is currently known.

Similarly, we identified Pepper mild mottle virus in U2. This virus has previously been proposed as a water quality indicator and an indicator of the presence of human faeces in freshwater, as it is the most abundant RNA virus in human faeces but rarely found in animal faecal matter (Kitajima et al. 2018; Rosario et al. 2009). Our ability to detect these viruses suggests this technique could be used for ongoing monitoring, including detecting the presence of pest species in the surrounding area and human-related viruses indicating contamination of faecal matter in the river. That none of these viruses were found in the pristine sites suggests these viruses are being introduced into the river from agricultural and urban run-off, and possibly also discharge of treated wastewater into the river. Indeed, at pristine sites we would generally expect viruses associated with New Zealand native plants and animals, most of which are yet to be described.

This study represents the first characterization of the virome of a New Zealand river. We observed a high abundance and diversity of viruses, including many that are both novel and highly divergent. Within the same river catchment, we identified viruses linked to agriculture and human presence (including possible links to sewage) in the farming and urban sites that were not present in the pristine sites. More broadly, this work provides the foundation for more detailed research on the impacts of human land-use on river viromes which will require larger sample sizes across multiple river systems. Our results show that human land-use impacted the viral community in the river, suggesting that further work is needed to reduce the impact of intensive farming and urbanization on the land and rivers.

## Acknowledgements

We thank Wendy Kay and Nigel French for their assistance in the field, and Anthony Pita for assistance with sample processing. We thank Ci-Xiu Li, Wei-Shan Chang, Jackie Mahar and Sabrina Sadiq for bioinformatics advice. We thank the Molecular Epidemiology and Public Health Laboratory at Massey University and the Behaviour, Ecology and Evolution Lab at the University of Sydney for use of their laboratory space. This research utilised the high-performance computing service, Artemis, provided by the Sydney Informatics Hub, Core Research Facility, University of Sydney.

## Data

Sequence data have been deposited in the Sequence Read Archive (SRA) under accession numbers SRR17234948-53. The trimmed alignment fasta files used to inder the phylogenetic trees are available at https://github.com/RKFrench/Viral-Diversity-NZ-River.

## References

1. Alegbeleye, O. O., and Sant’Ana, A. S. (2020) ‘Manure-borne pathogens as an important source of water contamination: An update on the dynamics of pathogen survival/transport as well as practical risk mitigation strategies’, International Journal of Hygiene and Environmental Health, 227: 113524.

2. Amvrosieva, T. et al. (2001) ‘Viral water contamination as the cause of aseptic meningitis outbreak in Belarus’, Central European Journal of Public Health 9: 154–7.

3. Baldisserotto, B. (2011) ‘Water pH and hardness affect growth of freshwater teleosts’, Brazilian Journal of Animal Science 40: 138–44.

4. Bolger, A.M., Lohse, M., and Usadel, B. (2014) ‘Trimmomatic: a flexible trimmer for Illumina sequence data’, Bioinformatics 30: 2114–20.

5. Buchfink, B., Reuter, and K., Drost, H.-G. (2021) ‘Sensitive protein alignments at tree-of-life scale using DIAMOND’, Nature Methods 18: 366–8.

6. Bulman, S. et al. (2005) ‘Rhopalosiphum aphids in New Zealand. I. RAPD markers reveal limited variability in lineages of *Rhopalosiphum padi**’*. New Zealand Journal of Zoology 32: 29–36.

7. Burridge, C. P., and Waters, J. M. (2020) ‘Does migration promote or inhibit diversification? A case study involving the dominant radiation of temperate Southern Hemisphere freshwater fishes’, Evolution 74: 1954–65.

8. Bushnell, B. (2018) ‘BBMap short-read aligner, and other bioinformatics tools’, University of California, Berkeley, CA.

9. Camacho, C. et al. (2009) ‘BLAST+: architecture and applications’, BMC Bioinformatics 10: 1–9.

10. Capella-Gutiérrez, S., Silla-Martínez, J. M., and Gabaldón, T. (2009) ‘trimAl: a tool for automated alignment trimming in large-scale phylogenetic analyses’, Bioinformatics 25: 1972–3.

11. Campbell, S. J. et al. (2020). ‘Red fox viromes in urban and rural landscapes’, Virus Evolution 6: veaa065.

12. Chapman, P. M., Bailey, H., and Canaria, E. (2000) ‘Toxicity of total dissolved solids associated with two mine effluents to chironomid larvae and early life stages of rainbow trout’, Environmental Toxicology and Chemistry 19: 210–14.

13. Chen, J. et al. (2019) ‘Detection and characterization of a novel marine birnavirus isolated from Asian seabass in Singapore’, Virology Journal 16: 1–10.

14. Chen, W. et al. (2018) ‘Aquatic bacterial communities associated with land use and environmental factors in agricultural landscapes using a metabarcoding approach’, Frontiers in Microbiology 9: 2301.

15. Clausen, P. T., Aarestrup, F.M., and Lund, O. (2018) ‘Rapid and precise alignment of raw reads against redundant databases with KMA’, BMC Bioinformatics 19: 1–8.

16. Da Costa, B. et al. (2003) ‘Blotched snakehead virus is a new aquatic birnavirus that is slightly more related to avibirnavirus than to aquabirnavirus’ Journal of Virology 77: 719-25.

17. Davies-Colley, R. J. (2013) ‘River water quality in New Zealand: an introduction and overview’, Ecosystem Services in New Zealand: Conditions and Trends. Manaaki Whenua Press, Lincoln: 432–447.

18. de Sousa, D. N. R. et al. (2014) ‘Electrical conductivity and emerging contaminant as markers of surface freshwater contamination by wastewater’, Science of the Total Environment 484: 19–26.

19. Department of Conservation (2021a). ‘History and culture’, https://www.doc.govt.nz/parks-and-recreation/places-to-go/manawatu-whanganui/places/ruahine-forest-park/history-and-culture/ (Accessed 1st June, 2021).

20. Department of Conservation (2021b). ‘Nature and conservation’, https://www.doc.govt.nz/parks-and-recreation/places-to-go/manawatu-whanganui/places/ruahine-forest-park/nature-and-conservation/ (Accessed 1st June, 2021).

21. Dos Anjos, K., Nagata, T., and de Melo, F. L. (2017) ‘Complete genome sequence of a novel bastrovirus isolated from raw sewage’, Genome Announcements 5: e01010–01017.

22. Dunn, N. R. et al. (2018) ‘Conservation status of New Zealand freshwater fishes, 2017’, Publishing Team, Department of Conservation.

23. Ferguson, C., et al. (2003) ‘Fate and transport of surface water pathogens in watersheds’ Critical Reviews in Environmental Science and Technology 33: 299–361.

24. French, R. K., and Holmes, E. C. (2020) ‘An ecosystems perspective on virus evolution and emergence’, Trends in Microbiology 28: 165–75.

25. Grabherr, M. G. et al. (2011) ‘Full-length transcriptome assembly from RNA-Seq data without a reference genome’, Nature Biotechnology 29: 644.

26. Guerrero-Latorre, L. et al. (2018) ‘Quito’s virome: Metagenomic analysis of viral diversity in urban streams of Ecuador’s capital city’, Science of the Total Environment 645: 1334–43.

27. Gu, X. et al. (2018) ‘Geospatial distribution of viromes in tropical freshwater ecosystems’, Water Research 137: 220–32.

28. Guerrero-Latorre, L. et al (2018) ‘Quito’s virome: Metagenomic analysis of viral diversity in urban streams of Ecuador’s capital city’, Science of the Total Environment 645: 1334–43.

29. Hamill, K. (2012) ‘Effects of Palmerston North City’s Wastewater Treatment Plant discharge on water quality and aquatic life in the Manawatu River’, Palmerston North City Council and Horizons Regional Council.

30. Hill, M. O. (1973) ‘Diversity and evenness: a unifying notation and its consequences’, Ecology 54: 427–32.

31. Jack, S., Bell, D., and Hewitt, J. (2013) ‘Norovirus contamination of a drinking water supply at a hotel resort’, New Zealand Medical Journal 126: 98–107.

32. Jones, T. H., et al. (2017) ‘Waterborne viruses and F-specific coliphages in mixed-use watersheds: microbial associations, host specificities, and affinities with environmental/land use factors’, Applied and Environmental Microbiology 83: e02763–02716.

33. Kapelinskaya, T. V., et al. (2011) ‘Expression strategy of densonucleosis virus from the German cockroach, *Blattella germanica*’, Journal of Virology 85: 11855–70.

34. Katoh, K., and Standley, D. M (2013) ‘MAFFT multiple sequence alignment software version 7: improvements in performance and usability’, Molecular Biology and Evolution 30: 772–80.

35. Kirs, M., et al. (2011) ‘Source tracking faecal contamination in an urbanised and a rural waterway in the Nelson-Tasman region, New Zealand’, New Zealand Journal of Marine and Freshwater Research 45: 43–58.

36. Kitajima, M., Sassi, H. P., and Torrey, J. R. (2018) ‘Pepper mild mottle virus as a water quality indicator’, NPJ Clean Water 1: 1–9.

37. Kraberger, S., et al. (2015) ‘Characterisation of a diverse range of circular replication-associated protein encoding DNA viruses recovered from a sewage treatment oxidation pond’, Infection, Genetics and Evolution 31: 73–86.

38. Kraberger, S., et al. (2013) ‘Discovery of Sclerotinia sclerotiorum hypovirulence-associated virus-1 in urban river sediments of Heathcote and Styx Rivers in Christchurch City, New Zealand’, Genome Announcements 1: e00559–00513.

39. Krumbholz, A. et al. (2017) ‘Genome sequence of a novel picorna-like RNA virus from feces of the Antarctic fur seal (*Arctocephalus gazella*)’, Genome Announcements 5: e01001–17.

40. Lamberto, I. et al. (2014) ‘Mycovirus-like DNA virus sequences from cattle serum and human brain and serum samples from multiple sclerosis patients’, Genome Announcements 2: e00848–00814.

41. Le Lay, C., et al. (2020) ‘Unmapped RNA virus diversity in termites and their symbionts’, Viruses 12: 1145.

42. Li, B., and Dewey, C. N. (2011) ‘RSEM: accurate transcript quantification from RNA-Seq data with or without a reference genome’, BMC Bioinformatics 12: 1–16.

43. Longworth, J., and Carey, G. (1976) ‘A small RNA virus with a divided genome from *Heteronychus arator* (F.)[*Coleoptera: Scarabaeidae*]’, Journal of General Virology 33: 31–40.

44. Lukashov, V. V., and Goudsmit, J. 2002. Evolutionary relationships among *Astroviridae*. Journal of General Virology 83: 1397–1405.

45. Marcelino, V. R., et al. (2020) ‘CCMetagen: comprehensive and accurate identification of eukaryotes and prokaryotes in metagenomic data’, Genome Biology 21: 1–15.

46. Martolia, H. C. S. et al. (2009) ‘An outbreak of hepatitis E tracked to a spring in the foothills of the Himalayas, India, 2005’, Indian Journal of Gastroenterology 28: 99–101.

47. Mahar, J. E., et al. (2020) ‘Comparative analysis of RNA virome composition in rabbits and associated ectoparasites’ Journal of Virology 94: e02119–02119.

48. McMurdie, P. J., and Holmes, S. (2013) ‘phyloseq: an R package for reproducible interactive analysis and graphics of microbiome census data’, PLoS One 8: e61217.

49. Mortimer, N. et al. (2019) ‘Late Cretaceous oceanic plate reorganization and the breakup of Zealandia and Gondwana’, Gondwana Research 65: 31–42.

50. Mukherjee, S. S. (2011) ‘Identification and characterization of tobamo and tombusviruses isolated from New Zealand waters’, PhD thesis, State University of New York College of Environmental Science and Forestry.

51. Mukherjee, S. S. et al. (2012) ‘New tombusviruses isolated from surface waters in New Zealand’, Australasian Plant Pathology 41: 79–84.

52. Nayfach, S. et al. (2021) ‘Metagenomic compendium of 189,680 DNA viruses from the human gut microbiome’, Nature Microbiology 6: 960–70.

53. Nguyen, L.-T. et al. (2015) ‘IQ-TREE: a fast and effective stochastic algorithm for estimating maximum-likelihood phylogenies’ Molecular Biology and Evolution 32: 268–74.

54. Paradis, E., and Schliep, K. (2019) ‘ape 5.0: an environment for modern phylogenetics and evolutionary analyses in R’ Bioinformatics 35: 526–28.

55. Pénzes, J. J. et al. (2020) ‘Reorganizing the family *Parvoviridae*: a revised taxonomy independent of the canonical approach based on host association’, Archives of Virology 165: 2133–2146.

56. Pettersson, J. H.-O. et al. (2017) ‘Characterizing the virome of *Ixodes ricinus* ticks from northern Europe’, Scientific Reports 7: 1–7.

57. Phiri, B. J. et al. (2020) ‘Does land use affect pathogen presence in New Zealand drinking water supplies?’, Water Research 185: 116229.

58. Phiri, B. J. et al (2021) ‘Microbial diversity in water and animal faeces: a metagenomic analysis to assess public health risk’, New Zealand Journal of Zoology 48: 188–201.

59. Porter, A. F. et al. (2021) ‘Metagenomic identification of viral sequences in laboratory reagents’, Viruses 13: 2122

60. Qiu, H. et al. (2020) ‘Metagenomic analysis revealed that the terrestrial pollutants override the effects of seasonal variation on microbiome in river sediments’, Bulletin of Environmental Contamination and Toxicology 105: 892–8.

61. R Core Team. (2021) ‘R: A language and environment for statistical computing’, R Foundation for Statistical Computing, Vienna, Austria.

62. Reuter, G. et al. (2015) ‘A highly divergent picornavirus in an amphibian, the smooth newt (*Lissotriton vulgaris*)’. Journal of General Virology 96: 2607–13.

63. Rice, P., Longden, I., and Bleasby, A. (2000) ‘EMBOSS: the European molecular biology open software suite’, Trends in Genetics 16: 276–7.

64. Riviere, C., and Rochon, D. (1990) ‘Nucleotide sequence and genomic organization of melon necrotic spot virus’, Journal of General Virology 71: 1887–96.

65. Roossinck, M. J. (2015) ‘Plants, viruses and the environment: ecology and mutualism’, Virology 479: 271–7.

66. Rosario, K. et al. (2009) ‘Pepper mild mottle virus as an indicator of fecal pollution’, Applied and Environmental Microbiology 75: 7261–7.

67. Roygard, J., McArthur, K., and Clark, M. (2012) ‘Diffuse contributions dominate over point sources of soluble nutrients in two sub-catchments of the Manawatu River, New Zealand’ New Zealand Journal of Marine and Freshwater Research 46: 219–41.

68. Sedyaningsih-Mamahit, E. et al. (2002) ‘First documented outbreak of hepatitis E virus transmission in Java, Indonesia’, Transactions of the Royal Society of Tropical Medicine and Hygiene 96: 398–404.

69. Sekwadi, P. et al. (2018) ‘Waterborne outbreak of gastroenteritis on the KwaZulu-natal coast, South Africa, December 2016/January 2017’, Epidemiology and Infection 146: 1318–25.

70. Sit, T. L., and Lommel, S. A. (2015) ‘Tombusviridae’ eLS: 1–9.

71. Skotnicki, M. et al. (1993) ‘The genomic sequence of cardamine chlorotic fleck carmovirus’, Journal of General Virology 74: 1933–7.

72. Sommers, P. et al. (2021) ‘Integrating viral metagenomics into an ecological framework’, Annual Review of Virology 8: 133–58.

73. Starr, E. P. et al. (2019) ‘Metatranscriptomic reconstruction reveals RNA viruses with the potential to shape carbon cycling in soil’, Proceedings of the National Academy of Sciences USA 116: 25900–8.

74. Steel, O. et al. (2016) ‘Circular replication-associated protein encoding DNA viruses identified in the faecal matter of various animals in New Zealand’, Infection, Genetics and Evolution 43: 151–64.

75. Thao, M. L. et al. (2001) ‘Genetic characterization of a putative Densovirus from the mealybug *Planococcus citri*’, Current Microbiology 43: 457–458.

76. Valles, S. et al. (2017) ‘ICTV virus taxonomy profile: *Dicistroviridae*’, Journal of General Virology 98: 355–6.

77. Van Rossum, T. et al. (2015) ‘Year-long metagenomic study of river microbiomes across land use and water quality’, Frontiers in Microbiology 6: 1405.

78. Varsani, A., and Krupovic, M. (2021) ‘Family *Genomoviridae*: 2021 taxonomy update’, Archives of Virology 166: 2911–26

79. Waldron, F.M., Stone, G. N., and Obbard, D. J. (2018) ‘Metagenomic sequencing suggests a diversity of RNA interference-like responses to viruses across multicellular eukaryotes’, PLoS Genetics 14: e1007533.

80. Walker, S., Monks, A. and Innes, J. G (2021) ‘Life history traits explain vulnerability of endemic forest birds and predict recovery after predator suppression’, New Zealand Journal of Ecology 45: 3447.

81. Wickham, H. (2016) ‘ggplot2: Elegant Graphics for Data Analysis’, Springer-Verlag New York.

82. Wilkinson, L. (2011) ‘venneuler: Venn and Euler Diagrams’, R package version 1.1–0. Available at http://CRAN.R-project.org/package=venneuler.

83. Wille, M. et al. (2020) ‘Sustained RNA virome diversity in Antarctic penguins and their ticks’, The ISME Journal 14: 1768–82.

84. Wille, M. et al. (2021) ‘RNA virome abundance and diversity is associated with host age in a bird species’, Virology 561: 98–106.

85. Williamson, W. et al. (2011) ‘Enteric viruses in New Zealand drinking-water sources’ Water Science and Technology 63: 1744–51.

86. Wilson, M. J. et al. (2016) ‘Developing a strategy for using entomopathogenic nematodes to control the African black beetle (*Heteronychus arator*) in New Zealand pastures and investigating temperature constraints’, Biological Control 93: 1–7.

87. Wolf, Y. I. et al. (2020) ‘Doubling of the known set of RNA viruses by metagenomic analysis of an aquatic virome’, Nature Microbiology 5: 1262–70.

88. Yu, G. (2020) ‘Using ggtree to visualize data on tree-like structures’, Current Protocols in Bioinformatics 69: e96.

89. Zhang, Y.-Z., Shi, M., and Holmes, E. C. (2018) ‘Using metagenomics to characterize an expanding virosphere’, Cell 172: 1168–72.

