## Supplementary Information for "Human land-use impacts viral diversity and abundance in a New Zealand river"

**Supplementary Table**

**Table S1**. Details of the sequence alignments used to infer the phylogenetic trees for each virus family. Percentage pairwise identity refers to the percentage of amino acid residues that are identical in the alignment, excluding gap-gap residues.

**Supplementary Figure Legends**

**Figure S1**. Phylogeny of the *Alphaflexiviridae* based on the RNA-dependent RNA polymerase sequence (alignment length of 1128 amino acids). Viruses obtained in this study are shown in blue and have a '+' after their name. Related viruses are shown in black. Virus abundance is expressed as the percentage of the total number of reads and represented by the size of each coloured symbol. The colour of each symbol refers to the site type. Black circles on nodes show bootstrap support values greater than 90%. Branches are scaled according to the number of amino acid substitutions per site, shown in the scale bar. The tree is midpoint rooted.

**Figure S2**. Phylogeny of the *Birnaviridae* based on the RNA-dependent RNA polymerase sequence (alignment length of 651 amino acids). Viruses obtained in this study are shown in blue and have a '+' after their name. Related viruses are shown in black. Virus abundance is expressed as the percentage of the total number of reads and represented by the size of each coloured symbol. The colour of each symbol refers to the site type. Black circles on nodes show bootstrap support values greater than 90%. Branches are scaled according to the number of amino acid substitutions per site, shown in the scale bar. The tree is midpoint rooted.

**Figure S3**. Phylogeny of the *Botourmiaviridae* based on the RNA-dependent RNA polymerase sequence (alignment length of 490 amino acids). Viruses obtained in this study are shown in blue and have a '+' after their name. Related viruses are shown in black. Virus abundance is expressed as the percentage of the total number of reads and represented by the size of each coloured symbol. The colour of each symbol refers to the site type. Black circles on nodes show bootstrap support values greater than 90%. Branches are scaled according to the number of amino acid substitutions per site, shown in the scale bar. The tree is midpoint rooted.

**Figure S4**. Phylogeny of the *Dicistroviridae* based on the non-structural polyprotein sequence (alignment length of 1123 amino acids). Viruses obtained in this study are shown in blue and have a '+' after their name. Related viruses are shown in black. Virus abundance is expressed as the percentage of the total number of reads and represented by the size of each coloured symbol. The colour of each symbol refers to the site type. Black circles on nodes show bootstrap support values greater than 90%. Branches are scaled according to the number of amino acid substitutions per site, shown in the scale bar. The tree is midpoint rooted.

**Figure S5**. Phylogeny of the *Genomoviridae* based on the replication-associated protein sequence (alignment length of 208 amino acids). Viruses obtained in this study are shown in blue and have a '+' after their name. Related viruses are shown in black. Virus abundance is expressed as the percentage of the total number of reads and represented by the size of each coloured symbol. The colour of each symbol refers to the site type. Black circles on nodes show bootstrap support values greater than 90%. Branches are scaled according to the number of amino acid substitutions per site, shown in the scale bar. The tree is midpoint rooted.

**Figure S6**. Phylogeny of the *Hepeviridae* based on the non-structural polyprotein sequence (alignment length of 817 amino acids). Viruses obtained in this study are shown in blue and have a '+' after their name. Related viruses are shown in black. Virus abundance is expressed as the percentage of the total number of reads and represented by the size of each coloured symbol. The colour of each symbol refers to the site type. Black circles on nodes show bootstrap support values greater than 90%. Branches are scaled according to the number of amino acid substitutions per site, shown in the scale bar. The tree is midpoint rooted.

**Figure S7**. Phylogeny of the *Iflaviridae* based on the polyprotein sequence (alignment length of 1423 amino acids). Viruses obtained in this study are shown in blue and have a '+' after their name. Related viruses are shown in black. Virus abundance is expressed as the percentage of the total number of reads and represented by the size of each coloured symbol. The colour of each symbol refers to the site type. Black circles on nodes show bootstrap support values greater than 90%. Branches are scaled according to the number of amino acid substitutions per site, shown in the scale bar. The tree is midpoint rooted.

**Figure S8**. Phylogeny of the *Leviviridae* based on the RNA-dependent RNA polymerase sequence (alignment length of 440 amino acids). Viruses obtained in this study are shown in blue and have a '+' after their name. Related viruses are shown in black. Virus abundance is expressed as the percentage of the total number of reads and represented by the size of each coloured symbol. The colour of each symbol refers to the site type. Black circles on nodes show bootstrap support values greater than 90%. Branches are scaled according to the number of amino acid substitutions per site, shown in the scale bar. The tree is midpoint rooted.

**Figure S9**. Phylogeny of the *Luteoviridae* based on the RNA-dependent RNA polymerase sequence (alignment length of 523 amino acids). Viruses obtained in this study are shown in blue and have a '+' after their name. Related viruses are shown in black. Virus abundance is expressed as the percentage of the total number of reads and represented by the size of each coloured symbol. The colour of each symbol refers to the site type. Black circles on nodes show bootstrap support values greater than 90%. Branches are scaled according to the number of amino acid substitutions per site, shown in the scale bar. The tree is midpoint rooted.

**Figure S10**. Phylogeny of the *Marnaviridae* based on the RNA-dependent RNA polymerase sequence (alignment length of 1156 amino acids). Viruses obtained in this study are shown in blue and have a '+' after their name. Related viruses are shown in black. Virus abundance is expressed as the percentage of the total number of reads and represented by the size of each coloured symbol. The colour of each symbol refers to the site type. Black circles on nodes show bootstrap support values greater than 90%. Branches are scaled according to the number of amino acid substitutions per site, shown in the scale bar. The tree is midpoint rooted.

**Figure S11**. Phylogeny of the *Microviridae* based on the replication associated protein sequence (alignment length of 355 amino acids). Viruses obtained in this study are shown in blue and have a '+' after their name. Related viruses are shown in black. Virus abundance is expressed as the percentage of the total number of reads and represented by the size of each coloured symbol. The colour of each symbol refers to the site type. Black circles on nodes show bootstrap support values greater than 90%. Branches are scaled according to the number of amino acid substitutions per site, shown in the scale bar. The tree is midpoint rooted.

**Figure S12**. Phylogeny of the *Myoviridae* based on the DNA polymerase sequence (alignment length of 440 amino acids). Viruses obtained in this study are shown in blue and have a '+' after their name. Related viruses are shown in black. Virus abundance is expressed as the percentage of the total number of reads and represented by the size of each coloured symbol. The colour of each symbol refers to the site type. Black circles on nodes show bootstrap support values greater than 90%. Branches are scaled according to the number of amino acid substitutions per site, shown in the scale bar. The tree is midpoint rooted.

**Figure S13**. Phylogeny of the *Narnaviridae* based on the RNA-dependent RNA polymerase sequence (alignment length of 1043 amino acids). Viruses obtained in this study are shown in blue and have a '+' after their name. Related viruses are shown in black. Virus abundance is expressed as the percentage of the total number of reads and represented by the size of each coloured symbol. The colour of each symbol refers to the site type. Black circles on nodes show bootstrap support values greater than 90%. Branches are scaled according to the number of amino acid substitutions per site, shown in the scale bar. The tree is midpoint rooted.

**Figure S14**. Phylogeny of the *Nodaviridae* based on the RNA-dependent RNA polymerase sequence (alignment length of 1232 amino acids). Viruses obtained in this study are shown in blue and have a '+' after their name. Related viruses are shown in black. Virus abundance is expressed as the percentage of the total number of reads and represented by the size of each coloured symbol. The colour of each symbol refers to the site type. Black circles on nodes show bootstrap support values greater than 90%. Branches are scaled according to the number of amino acid substitutions per site, shown in the scale bar. The tree is midpoint rooted.

**Figure S15**. Phylogeny of the *Partitiviridae* based on the RNA-dependent RNA polymerase sequence (alignment length of 407 amino acids). Viruses obtained in this study are shown in blue and have a '+' after their name. Related viruses are shown in black. Virus abundance is expressed as the percentage of the total number of reads and represented by the size of each coloured symbol. The colour of each symbol refers to the site type. Black circles on nodes show bootstrap support values greater than 90%. Branches are scaled according to the number of amino acid substitutions per site, shown in the scale bar. The tree is midpoint rooted.

**Figure S16**. Phylogeny of the *Peribunyaviridae* based on the RNA-dependent RNA polymerase sequence (alignment length of 1842 amino acids). Viruses obtained in this study are shown in blue and have a '+' after their name. Related viruses are shown in black. Virus abundance is expressed as the percentage of the total number of reads and represented by the size of each coloured symbol. The colour of each symbol refers to the site type. Black circles on nodes show bootstrap support values greater than 90%. Branches are scaled according to the number of amino acid substitutions per site, shown in the scale bar. The tree is midpoint rooted.

**Figure S17**. Phylogeny of the *Picornaviridae* based on the polyprotein sequence (alignment length of 1673 amino acids). Viruses obtained in this study are shown in blue and have a '+' after their name. Related viruses are shown in black. Virus abundance is expressed as the percentage of the total number of reads and represented by the size of each coloured symbol. The colour of each symbol refers to the site type. Black circles on nodes show bootstrap support values greater than 90%. Branches are scaled according to the number of amino acid substitutions per site, shown in the scale bar. The tree is midpoint rooted.

**Figure S18**. Phylogeny of the *Podoviridae* based on the DNA polymerase sequence (alignment length of 586 amino acids). Viruses obtained in this study are shown in blue and have a '+' after their name. Related viruses are shown in black. Virus abundance is expressed as the percentage of the total number of reads and represented by the size of each coloured symbol. The colour of each symbol refers to the site type. Black circles on nodes show bootstrap support values greater than 90%. Branches are scaled according to the number of amino acid substitutions per site, shown in the scale bar. The tree is midpoint rooted.

**Figure S19**. Phylogeny of the *Polycipiviridae* based on the RNA-dependent RNA polymerase sequence (alignment length of 1415 amino acids). Viruses obtained in this study are shown in blue and have a '+' after their name. Related viruses are shown in black. Virus abundance is expressed as the percentage of the total number of reads and represented by the size of each coloured symbol. The colour of each symbol refers to the site type. Black circles on nodes show bootstrap support values greater than 90%. Branches are scaled according to the number of amino acid substitutions per site, shown in the scale bar. The tree is midpoint rooted.

**Figure S20**. Phylogeny of the *Secoviridae* based on the RNA-dependent RNA polymerase sequence (alignment length of 1203 amino acids). Viruses obtained in this study are shown in blue and have a '+’ after their name. Related viruses are shown in black. Virus abundance is expressed as the percentage of the total number of reads and represented by the size of each coloured symbol. The colour of each symbol refers to the site type. Black circles on nodes show bootstrap support values greater than 90%. Branches are scaled according to the number of amino acid substitutions per site, shown in the scale bar. The tree is midpoint rooted.

**Figure S21**. Phylogeny of the *Siphoviridae* based on the DNA polymerase sequence (alignment length of 488 amino acids). Viruses obtained in this study are shown in blue and have a '+’ after their name. Related viruses are shown in black. Virus abundance is expressed as the percentage of the total number of reads and represented by the size of each coloured symbol. The colour of each symbol refers to the site type. Black circles on nodes show bootstrap support values greater than 90%. Branches are scaled according to the number of amino acid substitutions per site, shown in the scale bar. The tree is midpoint rooted.

**Figure S22**. Phylogeny of the *Solemoviridae* based on the RNA-dependent RNA polymerase sequence (alignment length of 736 amino acids). Viruses obtained in this study are shown in blue and have a '+’ after their name. Related viruses are shown in black. Virus abundance is expressed as the percentage of the total number of reads and represented by the size of each coloured symbol. The colour of each symbol refers to the site type. Black circles on nodes show bootstrap support values greater than 90%. Branches are scaled according to the number of amino acid substitutions per site, shown in the scale bar. The tree is midpoint rooted.

**Figure S23**. Phylogeny of the *Togaviridae* based on the RNA-dependent RNA polymerase sequence (alignment length of 940 amino acids). Viruses obtained in this study are shown in blue and have a '+’ after their name. Related viruses are shown in black. Virus abundance is expressed as the percentage of the total number of reads and represented by the size of each coloured symbol. The colour of each symbol refers to the site type. Black circles on nodes show bootstrap support values greater than 90%. Branches are scaled according to the number of amino acid substitutions per site, shown in the scale bar. The tree is midpoint rooted.

**Figure S24**. Phylogeny of the *Tombusviridae* based on the RNA-dependent RNA polymerase sequence (alignment length of 459 amino acids). Viruses obtained in this study are shown in blue and have a '+' after their name. Related viruses are shown in black. Virus abundance is expressed as the percentage of the total number of reads and represented by the size of each coloured symbol. The colour of each symbol refers to the site type. Black circles on nodes show bootstrap support values greater than 90%. Branches are scaled according to the number of amino acid substitutions per site, shown in the scale bar. The tree is midpoint rooted.

**Figure S25**. Phylogeny of the *Virgaviridae* based on the RNA-dependent RNA polymerase sequence (alignment length of 1081 amino acids). Viruses obtained in this study are shown in blue and have a '+' after their name. Related viruses are shown in black. Virus abundance is expressed as the percentage of the total number of reads and represented by the size of each coloured symbol. The colour of each symbol refers to the site type. Black circles on nodes show bootstrap support values greater than 90%. Branches are scaled according to the number of amino acid substitutions per site, shown in the scale bar. The tree is midpoint rooted.

| Viral family | Gene | Alignment length (AA) | Number of sequences | % Pairwise Identity |
|---|---|---|---|---|
| *Alphaflexiviridae* | RNA-dependent RNA polymerase | 1228 | 55 | 47 |
| *Astroviridae* | Non-structural polyprotein | 733 | 97 | 15.6 |
| *Birnaviridae* | RNA-dependent RNA polymerase | 651 | 23 | 41.9 |
| *Botourmiaviridae* | RNA-dependent RNA polymerase | 490 | 54 | 22.5 |
| *Dicistroviridae* | Non-structural polyprotein | 1123 | 151 | 20.9 |
| *Genomoviridae* | Replication-associated protein | 208 | 142 | 49 |
| *Hepeviridae* | Non-structural polyprotein | 817 | 58 | 28.9 |
| *Iflaviridae* | Polyprotein | 1423 | 61 | 22 |
| *Leviviridae* | RNA-dependent RNA polymerase | 440 | 21 | 34.9 |
| *Luteoviridae* | RNA-dependent RNA polymerase | 523 | 111 | 33.2 |
| *Marnaviridae* | RNA-dependent RNA polymerase | 1156 | 55 | 22.7 |
| *Microviridae* | Replication-associated protein | 355 | 173 | 23.9 |
| *Myoviridae* | DNA polymerase | 440 | 59 | 16.9 |
| *Narnaviridae* | RNA-dependent RNA polymerase | 1043 | 153 | 13.1 |
| *Nodaviridae* | RNA-dependent RNA polymerase | 1232 | 175 | 22.7 |
| *Partitiviridae* | RNA-dependent RNA polymerase | 407 | 81 | 27.5 |
| *Parvoviridae* | Non-structural protein | 452 | 155 | 20.7 |
| *Peribunyaviridae* | RNA-dependent RNA polymerase | 1842 | 235 | 41.5 |
| *Picornaviridae* | Polyprotein | 1673 | 275 | 26.3 |
| *Podoviridae* | DNA polymerase | 586 | 160 | 14.7 |
| *Polycipiviridae* | RNA-dependent RNA polymerase | 1415 | 18 | 29.1 |
| *Secoviridae* | RNA-dependent RNA polymerase | 1203 | 79 | 24.3 |
| *Siphoviridae* | DNA polymerase | 488 | 61 | 18.8 |
| *Solemoviridae* | RNA-dependent RNA polymerase | 736 | 100 | 22.6 |
| *Togaviridae* | RNA-dependent RNA polymerase | 940 | 57 | 41.3 |
| *Tombusviridae* | RNA-dependent RNA polymerase | 459 | 163 | 40.1 |
| *Virgaviridae* | RNA-dependent RNA polymerase | 1081 | 81 | 33.7 |

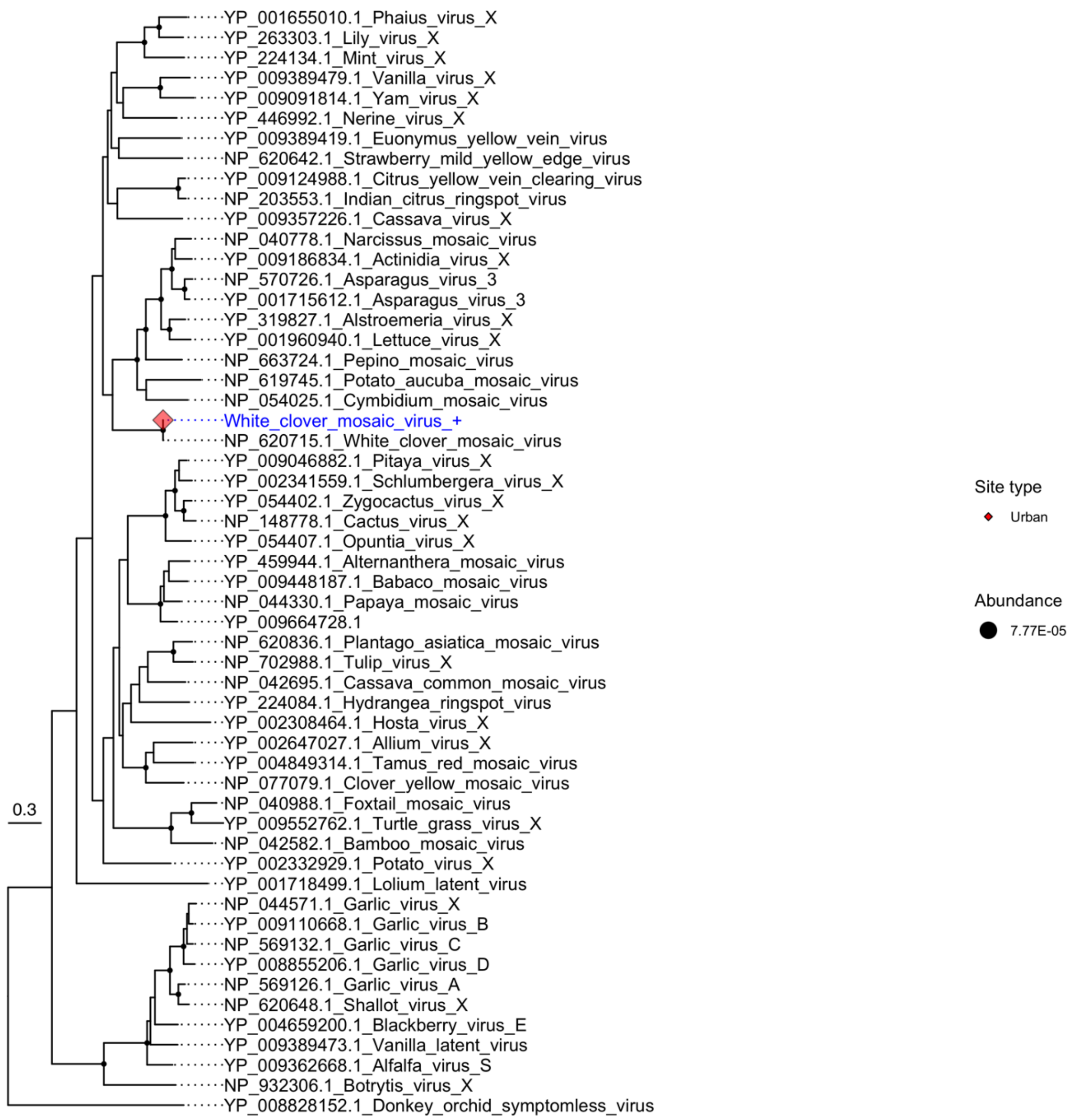

YP_001655010.1_Phaius_virus_X
YP_263303.1_Lily_virus_X
YP_224134.1_Mint_virus_X
YP_009389479.1_Vanilla_virus_X
YP_009091814.1_Yam_virus_X
YP_446992.1_Nerine_virus_X
YP_009389419.1_Euonymus_yellow_vein_virus
NP_620642.1_Strawberry_mild_yellow_edge_virus
YP_009124988.1_Citrus_yellow_vein_clearing_virus
NP_203553.1_Indian_citrus_ringspot_virus
YP_009357226.1_Cassava_virus_X
NP_040778.1_Narcissus_mosaic_virus
YP_009186834.1_Actinidia_virus_X
NP_570726.1_Asparagus_virus_3
YP_001715612.1_Asparagus_virus_3
YP_319827.1_Alstroemeria_virus_X
YP_001960940.1_Lettuce_virus_X
NP_663724.1_Pepino_mosaic_virus
NP_619745.1_Potato_aucuba_mosaic_virus
NP_054025.1_Cymbidium_mosaic_virus
White_clover_mosaic_virus_+
NP_620715.1_White_clover_mosaic_virus
YP_009046882.1_Pitaya_virus_X
YP_002341559.1_Schlumbergera_virus_X
YP_054402.1_Zygocactus_virus_X
NP_148778.1_Cactus_virus_X
YP_054407.1_Opuntia_virus_X
YP_459944.1_Alternanthera_mosaic_virus
YP_009448187.1_Babaco_mosaic_virus
NP_044330.1_Papaya_mosaic_virus
YP_009664728.1
NP_620836.1_Plantago_asiatica_mosaic_virus
NP_702988.1_Tulip_virus_X
NP_042695.1_Cassava_common_mosaic_virus
YP_224084.1_Hydrangea_ringspot_virus
YP_002308464.1_Hosta_virus_X
YP_002647027.1_Allium_virus_X
YP_004849314.1_Tamus_red_mosaic_virus
NP_077079.1_Clover_yellow_mosaic_virus
NP_040988.1_Foxtail_mosaic_virus
YP_009552762.1_Turtle_grass_virus_X
NP_042582.1_Bamboo_mosaic_virus
YP_002332929.1_Potato_virus_X
YP_001718499.1_Lolium_latent_virus
NP_044571.1_Garlic_virus_X
YP_009110668.1_Garlic_virus_B
NP_569132.1_Garlic_virus_C
YP_008855206.1_Garlic_virus_D
NP_569126.1_Garlic_virus_A
NP_620648.1_Shallot_virus_X
YP_004659200.1_Blackberry_virus_E
YP_009389473.1_Vanilla_latent_virus
YP_009362668.1_Alfalfa_virus_S
NP_932306.1_Botrytis_virus_X
YP_008828152.1_Donkey_orchid_symptomless_virus
0.3
Site type
Urban
Abundance
7.77E-05


**Figure S1**

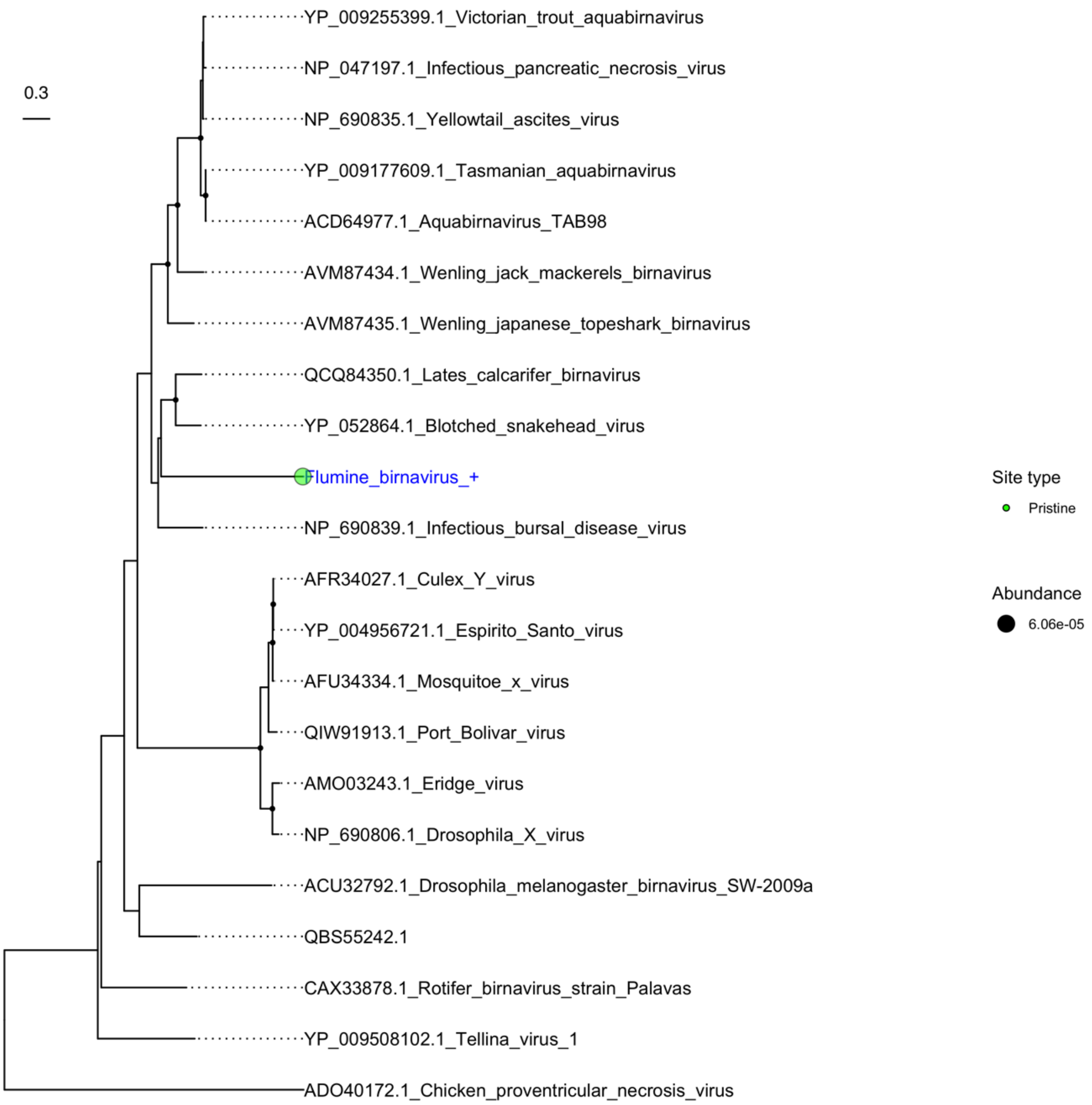
0.3
YP_009255399.1_Victorian_trout_aquabirnavirus
NP_047197.1_Infectious_pancreatic_necrosis_virus
NP_690835.1_Yellowtail_ascites_virus
YP_009177609.1_Tasmanian_aquabirnavirus
ACD64977.1_Aquabirnavirus_TAB98
AVM87434.1_Wenling_jack_mackerels_birnavirus
AVM87435.1_Wenling_japanese_topeshark_birnavirus
QCQ84350.1_Lates_calcarifer_birnavirus
YP_052864.1_Blotched_snakehead_virus
Flumine_birnavirus_+
NP_690839.1_Infectious_bursal_disease_virus
AFR34027.1_Culex_Y_virus
YP_004956721.1_Espirito_Santo_virus
AFU34334.1_Mosquitoe_x_virus
QIW91913.1_Port_Bolivar_virus
AMO03243.1_Eridge_virus
NP_690806.1_Drosophila_X_virus
ACU32792.1_Drosophila_melanogaster_birnavirus_SW-2009a
QBS55242.1
CAX33878.1_Rotifer_birnavirus_strain_Palavas
YP_009508102.1_Tellina_virus_1
ADO40172.1_Chicken_proventricular_necrosis_virus
Site type
Pristine
Abundance
6.06e-05

**Figure S2**

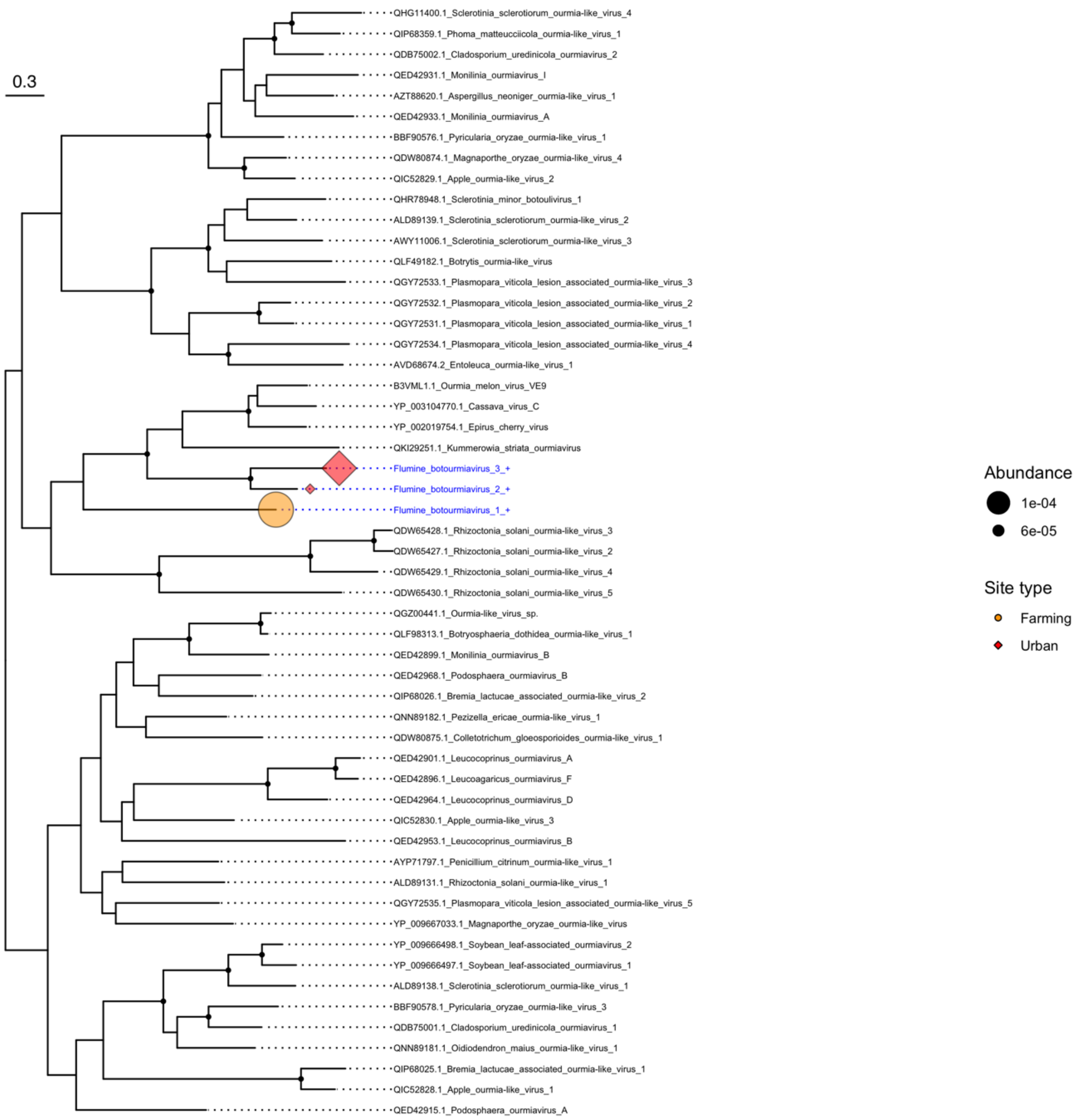
0.3
QHG11400.1_Sclerotinia_sclerotiorum_ourmia-like_virus_4
QIP68359.1_Phoma_matteucciicola_ourmia-like_virus_1
QDB75002.1_Cladosporium_uredinicola_ourmiavirus_2
QED42931.1_Monilinia_ourmiavirus_I
AZT88620.1_Aspergillus_neoniger_ourmia-like_virus_1
QED42933.1_Monilinia_ourmiavirus_A
BBF90576.1_Pyricularia_oryzae_ourmia-like_virus_1
QDW80874.1_Magnaporthe_oryzae_ourmia-like_virus_4
QIC52829.1_Apple_ourmia-like_virus_2
QHR78948.1_Sclerotinia_minor_botoulivirus_1
ALD89139.1_Sclerotinia_sclerotiorum_ourmia-like_virus_2
AWY11006.1_Sclerotinia_sclerotiorum_ourmia-like_virus_3
QLF49182.1_Botrytis_ourmia-like_virus
QGY72533.1_Plasmopara_viticola_lesion_associated_ourmia-like_virus_3
QGY72532.1_Plasmopara_viticola_lesion_associated_ourmia-like_virus_2
QGY72531.1_Plasmopara_viticola_lesion_associated_ourmia-like_virus_1
QGY72534.1_Plasmopara_viticola_lesion_associated_ourmia-like_virus_4
AVD68674.2_Entoleuca_ourmia-like_virus_1
B3VML1.1_Ourmia_melon_virus_VE9
YP_003104770.1_Cassava_virus_C
YP_002019754.1_Epirus_cherry_virus
QKI29251.1_Kummerowia_striata_ourmiavirus
Flumine_botourmiavirus_3_+
Flumine_botourmiavirus_2_+
Flumine_botourmiavirus_1_+
QDW65428.1_Rhizoctonia_solani_ourmia-like_virus_3
QDW65427.1_Rhizoctonia_solani_ourmia-like_virus_2
QDW65429.1_Rhizoctonia_solani_ourmia-like_virus_4
QDW65430.1_Rhizoctonia_solani_ourmia-like_virus_5
QGZ00441.1_Ourmia-like_virus_sp.
QLF98313.1_Botryosphaeria_dothidea_ourmia-like_virus_1
QED42899.1_Monilinia_ourmiavirus_B
QED42968.1_Podosphaera_ourmiavirus_B
QIP68026.1_Bremia_lactucae_associated_ourmia-like_virus_2
QNN89182.1_Pezizella_ericae_ourmia-like_virus_1
QDW80875.1_Colletotrichum_gloeosporioides_ourmia-like_virus_1
QED42901.1_Leucocoprinus_ourmiavirus_A
QED42896.1_Leucoagaricus_ourmiavirus_F
QED42964.1_Leucocoprinus_ourmiavirus_D
QIC52830.1_Apple_ourmia-like_virus_3
QED42953.1_Leucocoprinus_ourmiavirus_B
AYP71797.1_Penicillium_citrinum_ourmia-like_virus_1
ALD89131.1_Rhizoctonia_solani_ourmia-like_virus_1
QGY72535.1_Plasmopara_viticola_lesion_associated_ourmia-like_virus_5
YP_009667033.1_Magnaporthe_oryzae_ourmia-like_virus
YP_009666498.1_Soybean_leaf-associated_ourmiavirus_2
YP_009666497.1_Soybean_leaf-associated_ourmiavirus_1
ALD89138.1_Sclerotinia_sclerotiorum_ourmia-like_virus_1
BBF90578.1_Pyricularia_oryzae_ourmia-like_virus_3
QDB75001.1_Cladosporium_uredinicola_ourmiavirus_1
QNN89181.1_Oidiodendron_maius_ourmia-like_virus_1
QIP68025.1_Bremia_lactucae_associated_ourmia-like_virus_1
QIC52828.1_Apple_ourmia-like_virus_1
QED42915.1_Podosphaera_ourmiavirus_A
Abundance
1e-04
6e-05
Site type
Farming
Urban

**Figure S3**

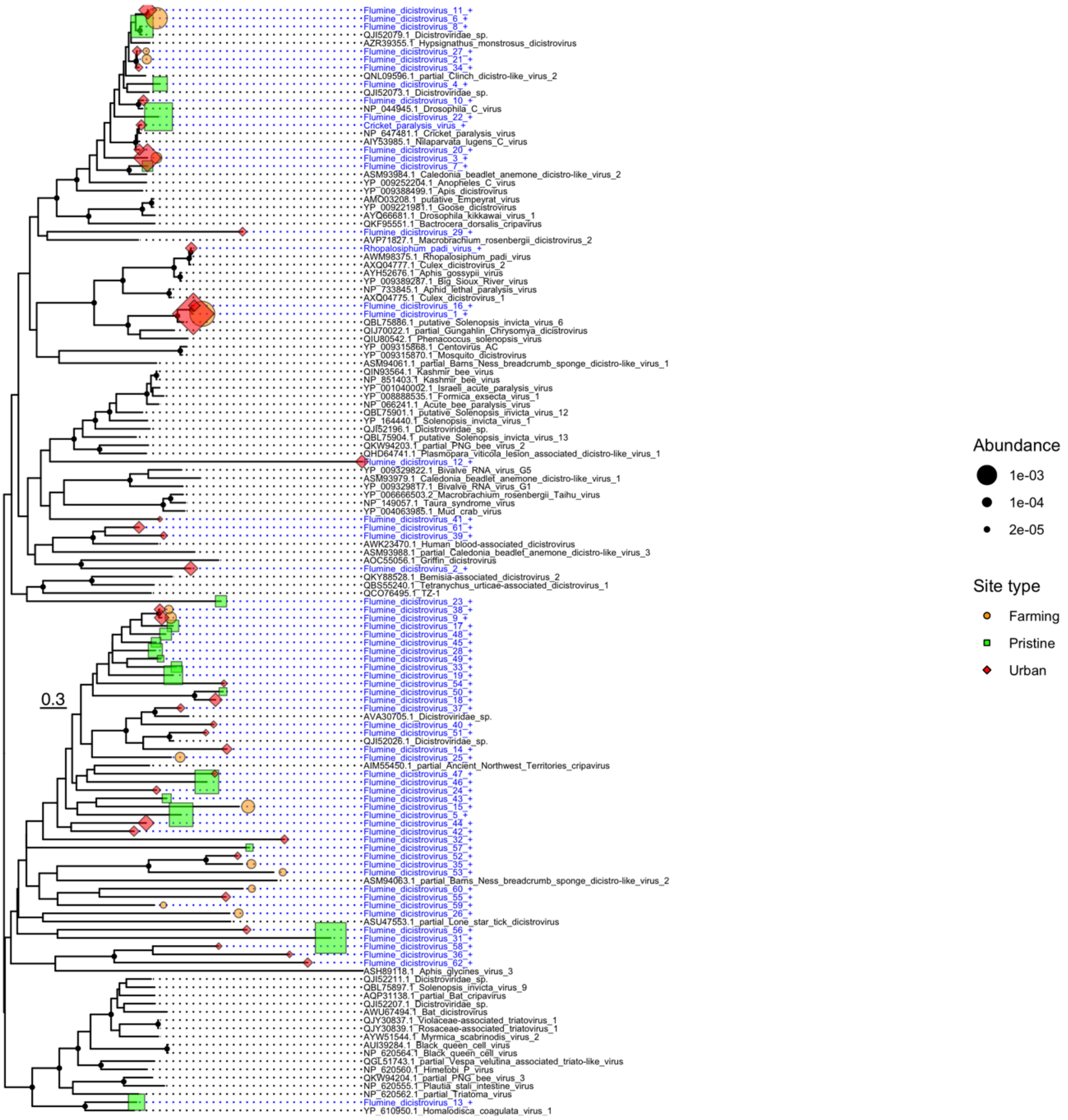
Flumine_dicistrovirus_11_+
Flumine_dicistrovirus_6_+
Flumine_dicistrovirus_8_+
QJI52079.1_Dicistroviridae_sp.
AZR39355.1_Hypsignathus_monstrosus_dicistrovirus
Flumine_dicistrovirus_27_+
Flumine_dicistrovirus_21_+
Flumine_dicistrovirus_34_+
QNL09596.1_partial_Clinch_dicistro-like_virus_2
Flumine_dicistrovirus_4_+
QJI52073.1_Dicistroviridae_sp.
Flumine_dicistrovirus_10_+
NP_044945.1_Drosophila_C_virus
Flumine_dicistrovirus_22_+
Cricket_paralysis_virus_+
NP_647481.1_Cricket_paralysis_virus
AIY53985.1_Nilaparvata_lugens_C_virus
Flumine_dicistrovirus_20_+
Flumine_dicistrovirus_3_+
Flumine_dicistrovirus_7_+
ASM93984.1_Caledonia_beadlet_anemone_dicistro-like_virus_2
YP_009252204.1_Anopheles_C_virus
YP_009388499.1_Apis_dicistrovirus
AMO03208.1_putative_Empeyrat_virus
YP_009221981.1_Goose_dicistrovirus
AYQ66681.1_Drosophila_kikkawai_virus_1
QKF95551.1_Bactrocera_dorsalis_cripavirus
Flumine_dicistrovirus_29_+
AVP71827.1_Macrobrachium_rosenbergii_dicistrovirus_2
Rhopalosiphum_padi_virus_+
AWM98375.1_Rhopalosiphum_padi_virus
AXQ04777.1_Culex_dicistrovirus_2
AYH52676.1_Aphis_gossypii_virus
YP_009389287.1_Big_Sioux_River_virus
NP_733845.1_Aphid_lethal_paralysis_virus
AXQ04775.1_Culex_dicistrovirus_1
Flumine_dicistrovirus_16_+
Flumine_dicistrovirus_1_+
QBL75886.1_putative_Solenopsis_invicta_virus_6
QIJ70022.1_partial_Gungahlin_Chrysomya_dicistrovirus
QIU80542.1_Phenacoccus_solenopsis_virus
YP_009315868.1_Centovirus_AC
YP_009315870.1_Mosquito_dicistrovirus
ASM94061.1_partial_Barns_Ness_breadcrumb_sponge_dicistro-like_virus_1
QIN93564.1_Kashmir_bee_virus
NP_851403.1_Kashmir_bee_virus
YP_001040002.1_Israeli_acute_paralysis_virus
YP_008888535.1_Formica_exsecta_virus_1
NP_066241.1_Acute_bee_paralysis_virus
QBL75901.1_putative_Solenopsis_invicta_virus_12
YP_164440.1_Solenopsis_invicta_virus_1
QJI52196.1_Dicistroviridae_sp.
QBL75904.1_putative_Solenopsis_invicta_virus_13
QKW94203.1_partial_PNG_bee_virus_2
QHD64741.1_Plasmopara_viticola_lesion_associated_dicistro-like_virus_1
Flumine_dicistrovirus_12_+
YP_009329822.1_Bivalve_RNA_virus_G5
ASM93979.1_Caledonia_beadlet_anemone_dicistro-like_virus_1
YP_009329817.1_Bivalve_RNA_virus_G1
YP_006666503.2_Macrobrachium_rosenbergii_Taihu_virus
NP_149057.1_Taura_syndrome_virus
YP_004063985.1_Mud_crab_virus
Flumine_dicistrovirus_41_+
Flumine_dicistrovirus_61_+
Flumine_dicistrovirus_39_+
AWK23470.1_Human_blood-associated_dicistrovirus
ASM93988.1_partial_Caledonia_beadlet_anemone_dicistro-like_virus_3
AOC55056.1_Griffin_dicistrovirus
Flumine_dicistrovirus_2_+
QKY88528.1_Bemisia-associated_dicistrovirus_2
QBS55240.1_Tetranychus_urticae-associated_dicistrovirus_1
QCO76495.1_TZ-1
Flumine_dicistrovirus_23_+
Flumine_dicistrovirus_38_+
Flumine_dicistrovirus_9_+
Flumine_dicistrovirus_17_+
Flumine_dicistrovirus_48_+
Flumine_dicistrovirus_45_+
Flumine_dicistrovirus_28_+
Flumine_dicistrovirus_49_+
Flumine_dicistrovirus_33_+
Flumine_dicistrovirus_19_+
Flumine_dicistrovirus_54_+
Flumine_dicistrovirus_50_+
Flumine_dicistrovirus_18_+
Flumine_dicistrovirus_37_+
AVA30705.1_Dicistroviridae_sp.
Flumine_dicistrovirus_40_+
Flumine_dicistrovirus_51_+
QJI52026.1_Dicistroviridae_sp.
Flumine_dicistrovirus_14_+
Flumine_dicistrovirus_25_+
AIM55450.1_partial_Ancient_Northwest_Territories_cripavirus
Flumine_dicistrovirus_47_+
Flumine_dicistrovirus_46_+
Flumine_dicistrovirus_24_+
Flumine_dicistrovirus_43_+
Flumine_dicistrovirus_15_+
Flumine_dicistrovirus_5_+
Flumine_dicistrovirus_44_+
Flumine_dicistrovirus_42_+
Flumine_dicistrovirus_32_+
Flumine_dicistrovirus_57_+
Flumine_dicistrovirus_52_+
Flumine_dicistrovirus_35_+
Flumine_dicistrovirus_53_+
ASM94063.1_partial_Barns_Ness_breadcrumb_sponge_dicistro-like_virus_2
Flumine_dicistrovirus_60_+
Flumine_dicistrovirus_55_+
Flumine_dicistrovirus_59_+
Flumine_dicistrovirus_26_+
ASU47553.1_partial_Lone_star_tick_dicistrovirus
Flumine_dicistrovirus_56_+
Flumine_dicistrovirus_31_+
Flumine_dicistrovirus_58_+
Flumine_dicistrovirus_36_+
Flumine_dicistrovirus_62_+
ASH89118.1_Aphis_glycines_virus_3
QJI52211.1_Dicistroviridae_sp.
QBL75897.1_Solenopsis_invicta_virus_9
AQP31138.1_partial_Bat_cripavirus
QJI52207.1_Dicistroviridae_sp.
AWU67494.1_Bat_dicistrovirus
QJY30837.1_Violaceae-associated_triatovirus_1
QJY30839.1_Rosaceae-associated_triatovirus_1
AYW51544.1_Myrmica_scabrinodis_virus_2
AUI39284.1_Black_queen_cell_virus
NP_620564.1_Black_queen_cell_virus
QGL51743.1_partial_Vespa_velutina_associated_triato-like_virus
NP_620560.1_Himetobi_P_virus
QKW94204.1_partial_PNG_bee_virus_3
NP_620555.1_Plautia_stali_intestine_virus
NP_620562.1_partial_Triatoma_virus
Flumine_dicistrovirus_13_+
YP_610950.1_Homalodisca_coagulata_virus_1
0.3
Abundance
1e-03
1e-04
2e-05
Site type
Farming
Pristine
Urban

**Figure S4**

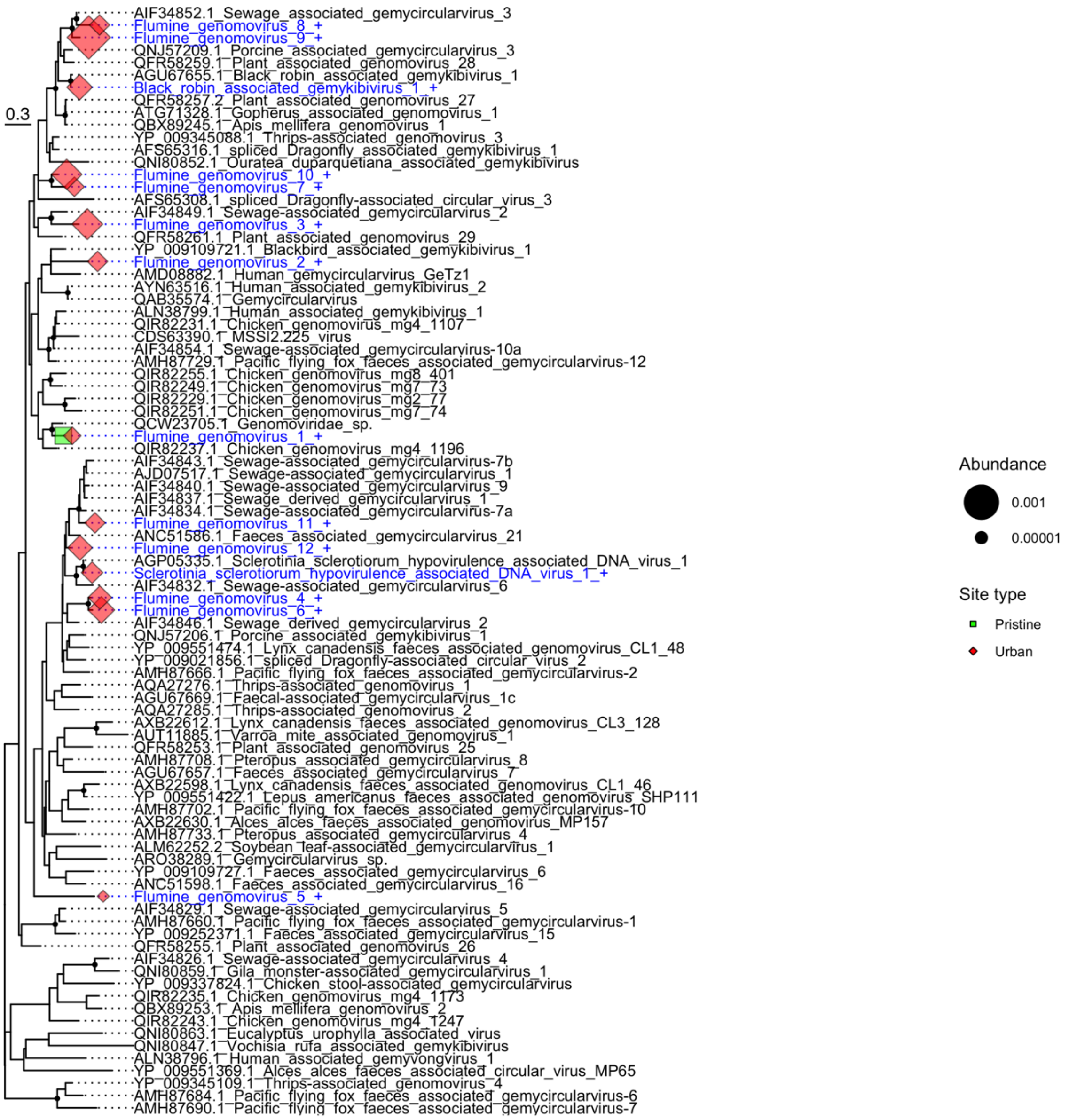

0.3
AIF34852.1_Sewage_associated_gemycircularvirus_3
Flumine_genomovirus_8_+
Flumine_genomovirus_9_+
QNJ57209.1_Porcine_associated_gemycircularvirus_3
QFR58259.1_Plant_associated_genomovirus_28
AGU67655.1_Black_robin_associated_gemykibivirus_1
Black_robin_associated_gemykibivirus_1_+
QFR58257.2_Plant_associated_genomovirus_27
ATG71328.1_Gopherus_associated_genomovirus_1
QBX89245.1_Apis_mellifera_genomovirus_1
YP_009345088.1_Thrips-associated_genomovirus_3
AFS65316.1_spliced_Dragonfly_associated_gemykibivirus_1
QNI80852.1_Ouratea_duparquetiana_associated_gemykibivirus
Flumine_genomovirus_10_+
Flumine_genomovirus_7_+
AFS65308.1_spliced_Dragonfly-associated_circular_virus_3
AIF34849.1_Sewage-associated_gemycircularvirus_2
Flumine_genomovirus_3_+
QFR58261.1_Plant_associated_genomovirus_29
YP_009109721.1_Blackbird_associated_gemykibivirus_1
Flumine_genomovirus_2_+
AMD08882.1_Human_gemycircularvirus_GeTz1
AYN63516.1_Human_associated_gemykibivirus_2
QAB35574.1_Gemycircularvirus
ALN38799.1_Human_associated_gemykibivirus_1
QIR82231.1_Chicken_genomovirus_mg4_1107
CDS63390.1_MSSI2.225_virus
AIF34854.1_Sewage-associated_gemycircularvirus-10a
AMH87729.1_Pacific_flying_fox_faeces_associated_gemycircularvirus-12
QIR82255.1_Chicken_genomovirus_mg8_401
QIR82249.1_Chicken_genomovirus_mg7_73
QIR82229.1_Chicken_genomovirus_mg2_77
QIR82251.1_Chicken_genomovirus_mg7_74
QCW23705.1_Genomoviridae_sp.
Flumine_genomovirus_1_+
QIR82237.1_Chicken_genomovirus_mg4_1196
AIF34843.1_Sewage-associated_gemycircularvirus-7b
AJD07517.1_Sewage-associated_gemycircularvirus_1
AIF34840.1_Sewage-associated_gemycircularvirus_9
AIF34837.1_Sewage_derived_gemycircularvirus_1
AIF34834.1_Sewage-associated_gemycircularvirus-7a
Flumine_genomovirus_11_+
ANC51586.1_Faeces_associated_gemycircularvirus_21
Flumine_genomovirus_12_+
AGP05335.1_Sclerotinia_sclerotiorum_hypovirulence_associated_DNA_virus_1
Sclerotinia_sclerotiorum_hypovirulence_associated_DNA_virus_1_+
AIF34832.1_Sewage-associated_gemycircularvirus_6
Flumine_genomovirus_4_+
Flumine_genomovirus_6_+
AIF34846.1_Sewage_derived_gemycircularvirus_2
QNJ57206.1_Porcine_associated_gemykibivirus_1
YP_009551474.1_Lynx_canadensis_faeces_associated_genomovirus_CL1_48
YP_009021856.1_spliced_Dragonfly-associated_circular_virus_2
AMH87666.1_Pacific_flying_fox_faeces_associated_gemycircularvirus-2
AQA27276.1_Thrips-associated_genomovirus_1
AGU67669.1_Faecal-associated_gemycircularvirus_1c
AQA27285.1_Thrips-associated_genomovirus_2
AXB22612.1_Lynx_canadensis_faeces_associated_genomovirus_CL3_128
AUT11885.1_Varroa_mite_associated_genomovirus_1
QFR58253.1_Plant_associated_genomovirus_25
AMH87708.1_Pteropus_associated_gemycircularvirus_8
AGU67657.1_Faeces_associated_gemycircularvirus_7
AXB22598.1_Lynx_canadensis_faeces_associated_genomovirus_CL1_46
YP_009551422.1_Lepus_americanus_faeces_associated_genomovirus_SHP111
AMH87702.1_Pacific_flying_fox_faeces_associated_gemycircularvirus-10
AXB22630.1_Alces_alces_faeces_associated_genomovirus_MP157
AMH87733.1_Pteropus_associated_gemycircularvirus_4
ALM62252.2_Soybean_leaf-associated_gemycircularvirus_1
ARO38289.1_Gemycircularvirus_sp.
YP_009109727.1_Faeces_associated_gemycircularvirus_6
ANC51598.1_Faeces_associated_gemycircularvirus_16
Flumine_genomovirus_5_+
AIF34829.1_Sewage-associated_gemycircularvirus_5
AMH87660.1_Pacific_flying_fox_faeces_associated_gemycircularvirus-1
YP_009252371.1_Faeces_associated_gemycircularvirus_15
QFR58255.1_Plant_associated_genomovirus_26
AIF34826.1_Sewage-associated_gemycircularvirus_4
QNI80859.1_Gila_monster-associated_gemycircularvirus_1
YP_009337824.1_Chicken_stool-associated_gemycircularvirus
QIR82235.1_Chicken_genomovirus_mg4_1173
QBX89253.1_Apis_mellifera_genomovirus_2
QIR82243.1_Chicken_genomovirus_mg4_1247
QNI80863.1_Eucalyptus_urophylla_associated_virus
QNI80847.1_Vochisia_rufa_associated_gemykibivirus
ALN38796.1_Human_associated_gemyvongvirus_1
YP_009551369.1_Alces_alces_faeces_associated_circular_virus_MP65
YP_009345109.1_Thrips-associated_genomovirus_4
AMH87684.1_Pacific_flying_fox_faeces_associated_gemycircularvirus-6
AMH87690.1_Pacific_flying_fox_faeces_associated_gemycircularvirus-7
Abundance
0.001
0.00001
Site type
Pristine
Urban


**Figure S5**

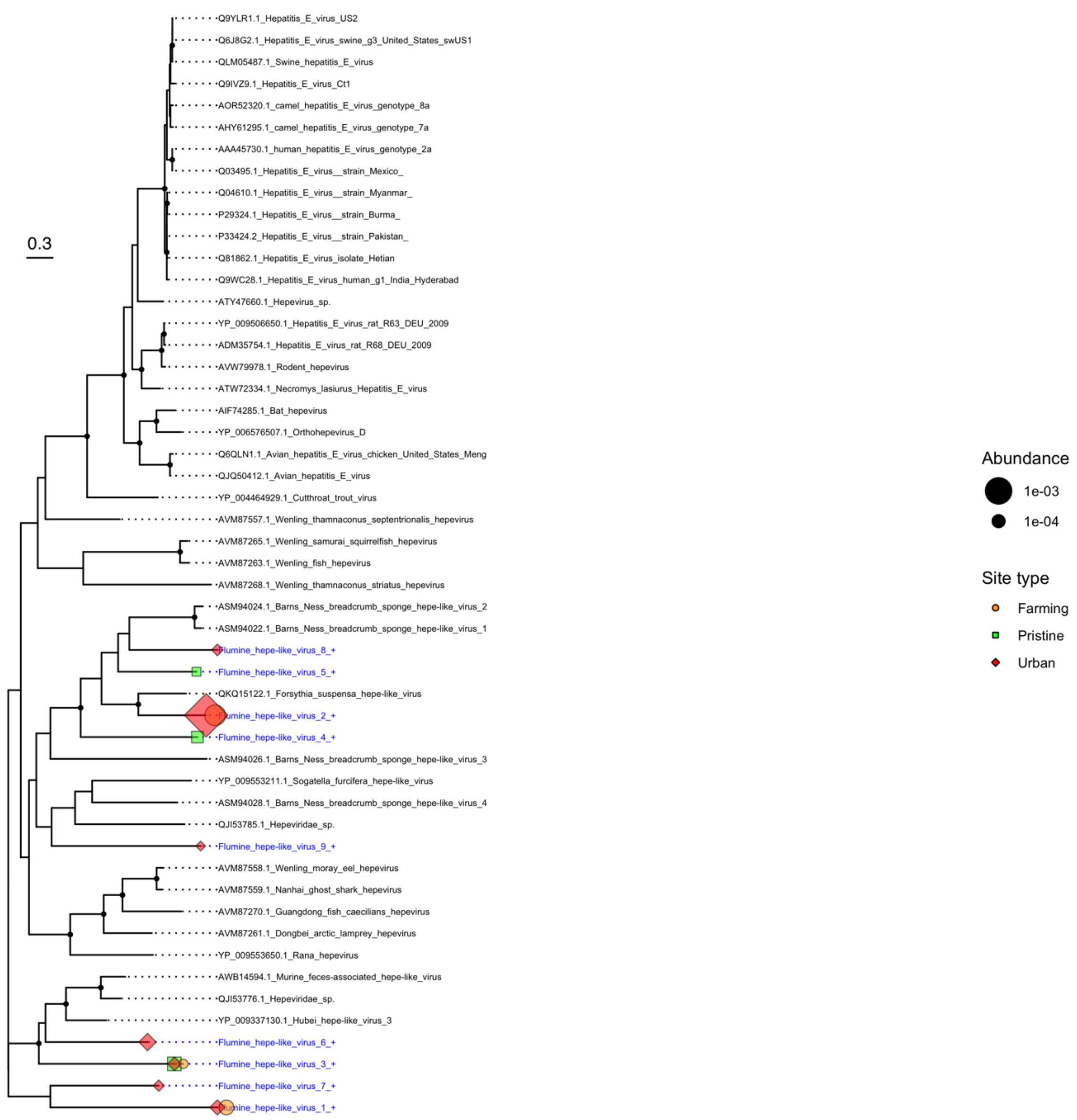
0.3
Q9YLR1.1_Hepatitis_E_virus_US2
Q6J8G2.1_Hepatitis_E_virus_swine_g3_United_States_swUS1
QLM05487.1_Swine_hepatitis_E_virus
Q9IVZ9.1_Hepatitis_E_virus_Ct1
AOR52320.1_camel_hepatitis_E_virus_genotype_8a
AHY61295.1_camel_hepatitis_E_virus_genotype_7a
AAA45730.1_human_hepatitis_E_virus_genotype_2a
Q03495.1_Hepatitis_E_virus__strain_Mexico_
Q04610.1_Hepatitis_E_virus__strain_Myanmar_
P29324.1_Hepatitis_E_virus__strain_Burma_
P33424.2_Hepatitis_E_virus__strain_Pakistan_
Q81862.1_Hepatitis_E_virus_isolate_Hetian
Q9WC28.1_Hepatitis_E_virus_human_g1_India_Hyderabad
ATY47660.1_Hepevirus_sp.
YP_009506650.1_Hepatitis_E_virus_rat_R63_DEU_2009
ADM35754.1_Hepatitis_E_virus_rat_R68_DEU_2009
AVW79978.1_Rodent_hepevirus
ATW72334.1_Necromys_lasiurus_Hepatitis_E_virus
AIF74285.1_Bat_hepevirus
YP_006576507.1_Orthohepevirus_D
Q6QLN1.1_Avian_hepatitis_E_virus_chicken_United_States_Meng
QJQ50412.1_Avian_hepatitis_E_virus
YP_004464929.1_Cutthroat_trout_virus
AVM87557.1_Wenling_thamnaconus_septentrionalis_hepevirus
AVM87265.1_Wenling_samurai_squirrelfish_hepevirus
AVM87263.1_Wenling_fish_hepevirus
AVM87268.1_Wenling_thamnaconus_striatus_hepevirus
ASM94024.1_Barns_Ness_breadcrumb_sponge_hepe-like_virus_2
ASM94022.1_Barns_Ness_breadcrumb_sponge_hepe-like_virus_1
Flumine_hepe-like_virus_8_+
Flumine_hepe-like_virus_5_+
QKQ15122.1_Forsythia_suspensa_hepe-like_virus
Flumine_hepe-like_virus_2_+
Flumine_hepe-like_virus_4_+
ASM94026.1_Barns_Ness_breadcrumb_sponge_hepe-like_virus_3
YP_009553211.1_Sogatella_furcifera_hepe-like_virus
ASM94028.1_Barns_Ness_breadcrumb_sponge_hepe-like_virus_4
QJI53785.1_Hepeviridae_sp.
Flumine_hepe-like_virus_9_+
AVM87558.1_Wenling_moray_eel_hepevirus
AVM87559.1_Nanhai_ghost_shark_hepevirus
AVM87270.1_Guangdong_fish_caecilians_hepevirus
AVM87261.1_Dongbei_arctic_lamprey_hepevirus
YP_009553650.1_Rana_hepevirus
AWB14594.1_Murine_feces-associated_hepe-like_virus
QJI53776.1_Hepeviridae_sp.
YP_009337130.1_Hubei_hepe-like_virus_3
Flumine_hepe-like_virus_6_+
Flumine_hepe-like_virus_3_+
Flumine_hepe-like_virus_7_+
Flumine_hepe-like_virus_1_+
Abundance
1e-03
1e-04
Site type
Farming
Pristine
Urban

**Figure S6**

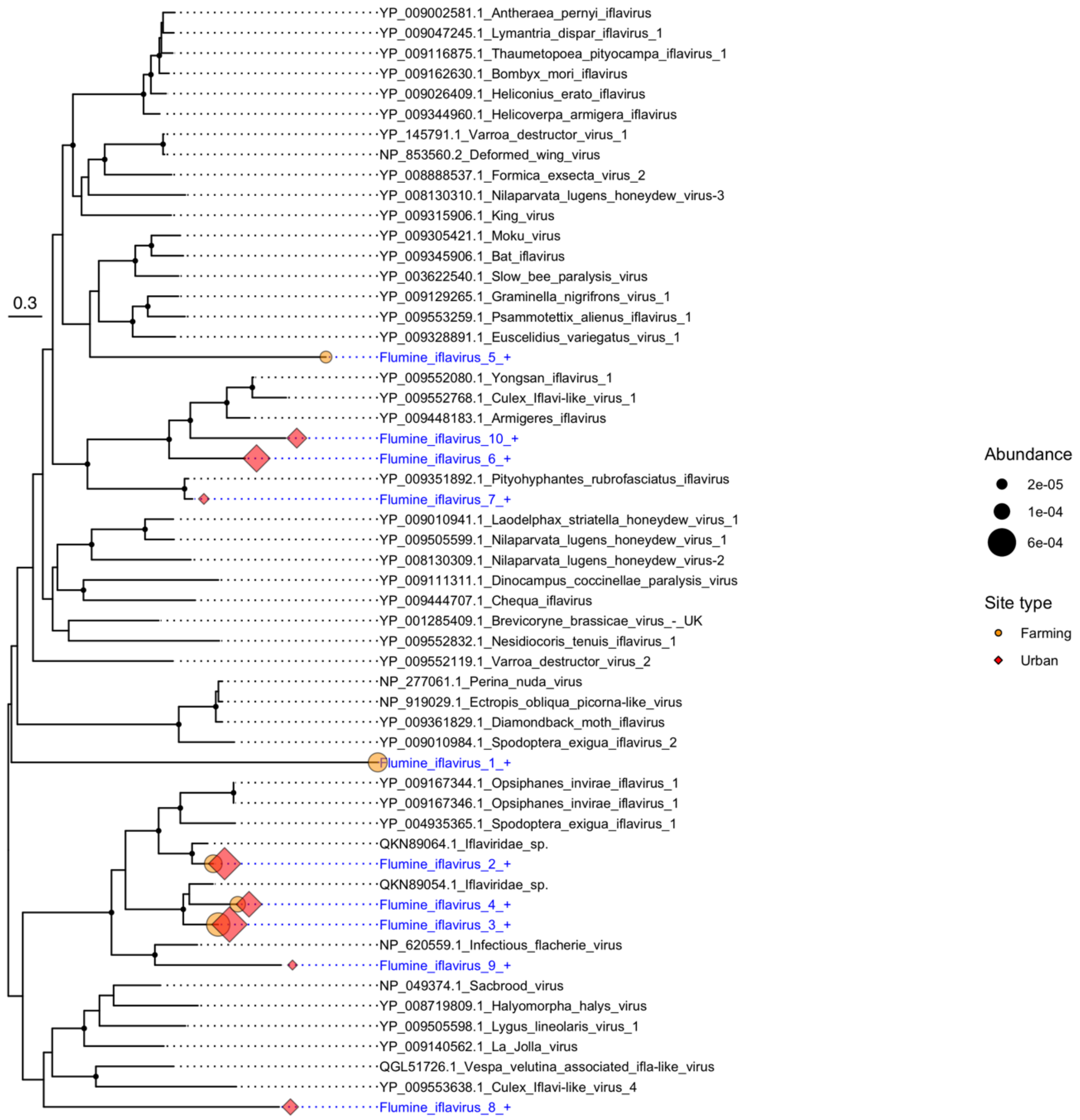
YP_009002581.1_Antheraea_pernyi_iflavirus
YP_009047245.1_Lymantria_dispar_iflavirus_1
YP_009116875.1_Thaumetopoea_pityocampa_iflavirus_1
YP_009162630.1_Bombyx_mori_iflavirus
YP_009026409.1_Heliconius_erato_iflavirus
YP_009344960.1_Helicoverpa_armigera_iflavirus
YP_145791.1_Varroa_destructor_virus_1
NP_853560.2_Deformed_wing_virus
YP_008888537.1_Formica_exsecta_virus_2
YP_008130310.1_Nilaparvata_lugens_honeydew_virus-3
YP_009315906.1_King_virus
YP_009305421.1_Moku_virus
YP_009345906.1_Bat_iflavirus
YP_003622540.1_Slow_bee_paralysis_virus
YP_009129265.1_Graminella_nigrifrons_virus_1
YP_009553259.1_Psammotettix_alienus_iflavirus_1
YP_009328891.1_Euscelidius_variegatus_virus_1
0.3
Flumine_iflavirus_5_+
YP_009552080.1_Yongsan_iflavirus_1
YP_009552768.1_Culex_Iflavi-like_virus_1
YP_009448183.1_Armigeres_iflavirus
Flumine_iflavirus_10_+
Flumine_iflavirus_6_+
YP_009351892.1_Pityohyphantes_rubrofasciatus_iflavirus
Flumine_iflavirus_7_+
YP_009010941.1_Laodelphax_striatella_honeydew_virus_1
YP_009505599.1_Nilaparvata_lugens_honeydew_virus_1
YP_008130309.1_Nilaparvata_lugens_honeydew_virus-2
YP_009111311.1_Dinocampus_coccinellae_paralysis_virus
YP_009444707.1_Chequa_iflavirus
YP_001285409.1_Brevicoryne_brassicae_virus_-_UK
YP_009552832.1_Nesidiocoris_tenuis_iflavirus_1
YP_009552119.1_Varroa_destructor_virus_2
NP_277061.1_Perina_nuda_virus
NP_919029.1_Ectropis_obliqua_picorna-like_virus
YP_009361829.1_Diamondback_moth_iflavirus
YP_009010984.1_Spodoptera_exigua_iflavirus_2
Flumine_iflavirus_1_+
YP_009167344.1_Opsiphanes_invirae_iflavirus_1
YP_009167346.1_Opsiphanes_invirae_iflavirus_1
YP_004935365.1_Spodoptera_exigua_iflavirus_1
QKN89064.1_Iflaviridae_sp.
Flumine_iflavirus_2_+
QKN89054.1_Iflaviridae_sp.
Flumine_iflavirus_4_+
Flumine_iflavirus_3_+
NP_620559.1_Infectious_flacherie_virus
Flumine_iflavirus_9_+
NP_049374.1_Sacbrood_virus
YP_008719809.1_Halyomorpha_halys_virus
YP_009505598.1_Lygus_lineolaris_virus_1
YP_009140562.1_La_Jolla_virus
QGL51726.1_Vespa_velutina_associated_ifla-like_virus
YP_009553638.1_Culex_Iflavi-like_virus_4
Flumine_iflavirus_8_+
Abundance
2e-05
1e-04
6e-04
Site type
Farming
Urban

**Figure S7**

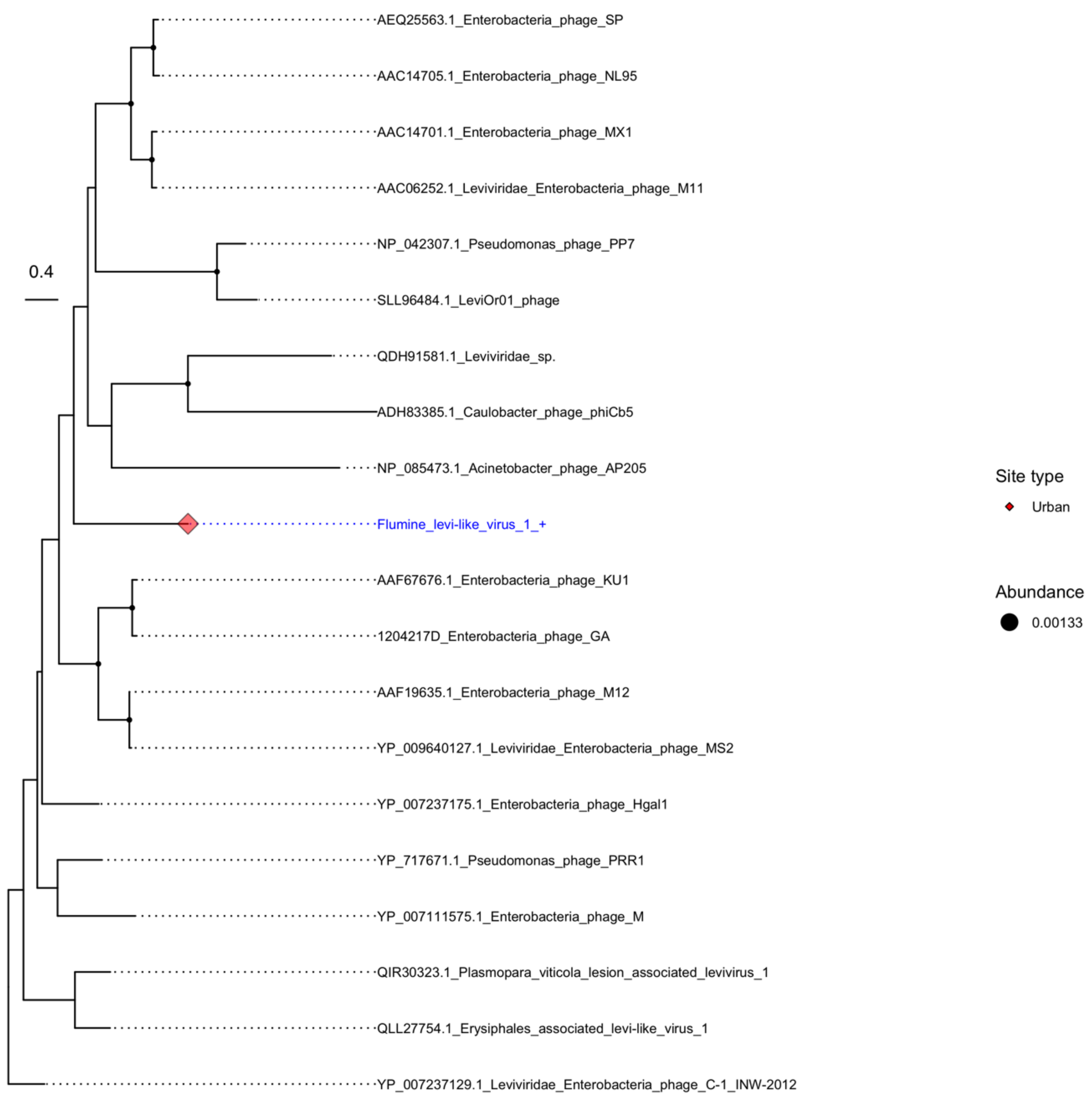
AEQ25563.1_Enterobacteria_phage_SP
AAC14705.1_Enterobacteria_phage_NL95
AAC14701.1_Enterobacteria_phage_MX1
AAC06252.1_Leviviridae_Enterobacteria_phage_M11
NP_042307.1_Pseudomonas_phage_PP7
0.4
SLL96484.1_LeviOr01_phage
QDH91581.1_Leviviridae_sp.
ADH83385.1_Caulobacter_phage_phiCb5
NP_085473.1_Acinetobacter_phage_AP205
Flumine_levi-like_virus_1_+
AAF67676.1_Enterobacteria_phage_KU1
1204217D_Enterobacteria_phage_GA
AAF19635.1_Enterobacteria_phage_M12
YP_009640127.1_Leviviridae_Enterobacteria_phage_MS2
YP_007237175.1_Enterobacteria_phage_Hgal1
YP_717671.1_Pseudomonas_phage_PRR1
YP_007111575.1_Enterobacteria_phage_M
QIR30323.1_Plasmopara_viticola_lesion_associated_levivirus_1
QLL27754.1_Erysiphales_associated_levi-like_virus_1
YP_007237129.1_Leviviridae_Enterobacteria_phage_C-1_INW-2012
Site type
Urban
Abundance
0.00133

**Figure S8**

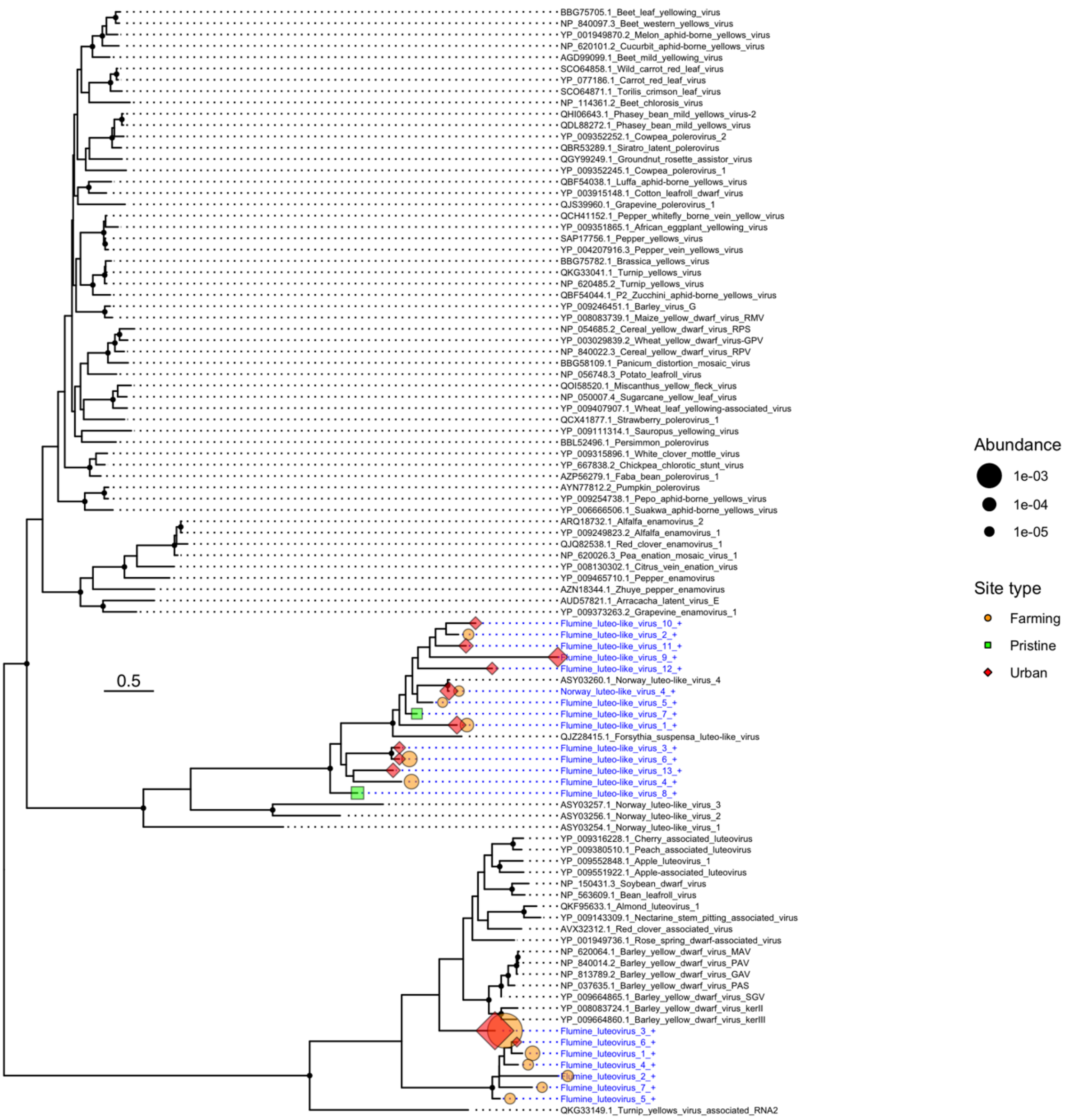
BBG75705.1_Beet_leaf_yellowing_virus
NP_840097.3_Beet_western_yellows_virus
YP_001949870.2_Melon_aphid-borne_yellows_virus
NP_620101.2_Cucurbit_aphid-borne_yellows_virus
AGD99099.1_Beet_mild_yellowing_virus
SCO64858.1_Wild_carrot_red_leaf_virus
YP_077186.1_Carrot_red_leaf_virus
SCO64871.1_Torilis_crimson_leaf_virus
NP_114361.2_Beet_chlorosis_virus
QHI06643.1_Phasey_bean_mild_yellows_virus-2
QDL88272.1_Phasey_bean_mild_yellows_virus
YP_009352252.1_Cowpea_polerovirus_2
QBR53289.1_Siratro_latent_polerovirus
QGY99249.1_Groundnut_rosette_assistor_virus
YP_009352245.1_Cowpea_polerovirus_1
QBF54038.1_Luffa_aphid-borne_yellows_virus
YP_003915148.1_Cotton_leafroll_dwarf_virus
QJS39960.1_Grapevine_polerovirus_1
QCH41152.1_Pepper_whitefly_borne_vein_yellow_virus
YP_009351865.1_African_eggplant_yellowing_virus
SAP17756.1_Pepper_yellows_virus
YP_004207916.3_Pepper_vein_yellows_virus
BBG75782.1_Brassica_yellows_virus
QKG33041.1_Turnip_yellows_virus
NP_620485.2_Turnip_yellows_virus
QBF54044.1_P2_Zucchini_aphid-borne_yellows_virus
YP_009246451.1_Barley_virus_G
YP_008083739.1_Maize_yellow_dwarf_virus_RMV
NP_054685.2_Cereal_yellow_dwarf_virus_RPS
YP_003029839.2_Wheat_yellow_dwarf_virus-GPV
NP_840022.3_Cereal_yellow_dwarf_virus_RPV
BBG58109.1_Panicum_distortion_mosaic_virus
NP_056748.3_Potato_leafroll_virus
QOI58520.1_Miscanthus_yellow_fleck_virus
NP_050007.4_Sugarcane_yellow_leaf_virus
YP_009407907.1_Wheat_leaf_yellowing-associated_virus
QCX41877.1_Strawberry_polerovirus_1
YP_009111314.1_Sauropus_yellowing_virus
BBL52496.1_Persimmon_polerovirus
YP_009315896.1_White_clover_mottle_virus
YP_667838.2_Chickpea_chlorotic_stunt_virus
AZP56279.1_Faba_bean_polerovirus_1
AYN77812.2_Pumpkin_polerovirus
YP_009254738.1_Pepo_aphid-borne_yellows_virus
YP_006666506.1_Suakwa_aphid-borne_yellows_virus
ARQ18732.1_Alfalfa_enamovirus_2
YP_009249823.2_Alfalfa_enamovirus_1
QJQ82538.1_Red_clover_enamovirus_1
NP_620026.3_Pea_enation_mosaic_virus_1
YP_008130302.1_Citrus_vein_enation_virus
YP_009465710.1_Pepper_enamovirus
AZN18344.1_Zhuye_pepper_enamovirus
AUD57821.1_Arracacha_latent_virus_E
YP_009373263.2_Grapevine_enamovirus_1
Flumine_luteo-like_virus_10_+
Flumine_luteo-like_virus_2_+
Flumine_luteo-like_virus_11_+
Flumine_luteo-like_virus_9_+
Flumine_luteo-like_virus_12_+
ASY03260.1_Norway_luteo-like_virus_4
Norway_luteo-like_virus_4_+
Flumine_luteo-like_virus_5_+
Flumine_luteo-like_virus_7_+
Flumine_luteo-like_virus_1_+
QJZ28415.1_Forsythia_suspensa_luteo-like_virus
Flumine_luteo-like_virus_3_+
Flumine_luteo-like_virus_6_+
Flumine_luteo-like_virus_13_+
Flumine_luteo-like_virus_4_+
Flumine_luteo-like_virus_8_+
ASY03257.1_Norway_luteo-like_virus_3
ASY03256.1_Norway_luteo-like_virus_2
ASY03254.1_Norway_luteo-like_virus_1
YP_009316228.1_Cherry_associated_luteovirus
YP_009380510.1_Peach_associated_luteovirus
YP_009552848.1_Apple_luteovirus_1
YP_009551922.1_Apple-associated_luteovirus
NP_150431.3_Soybean_dwarf_virus
NP_563609.1_Bean_leafroll_virus
QKF95633.1_Almond_luteovirus_1
YP_009143309.1_Nectarine_stem_pitting_associated_virus
AVX32312.1_Red_clover_associated_virus
YP_001949736.1_Rose_spring_dwarf-associated_virus
NP_620064.1_Barley_yellow_dwarf_virus_MAV
NP_840014.2_Barley_yellow_dwarf_virus_PAV
NP_813789.2_Barley_yellow_dwarf_virus_GAV
NP_037635.1_Barley_yellow_dwarf_virus_PAS
YP_009664865.1_Barley_yellow_dwarf_virus_SGV
YP_008083724.1_Barley_yellow_dwarf_virus_kerII
YP_009664860.1_Barley_yellow_dwarf_virus_kerIII
Flumine_luteovirus_3_+
Flumine_luteovirus_6_+
Flumine_luteovirus_1_+
Flumine_luteovirus_4_+
Flumine_luteovirus_2_+
Flumine_luteovirus_7_+
Flumine_luteovirus_5_+
QKG33149.1_Turnip_yellows_virus_associated_RNA2
0.5
Abundance
1e-03
1e-04
1e-05
Site type
Farming
Pristine
Urban

### Figure S9

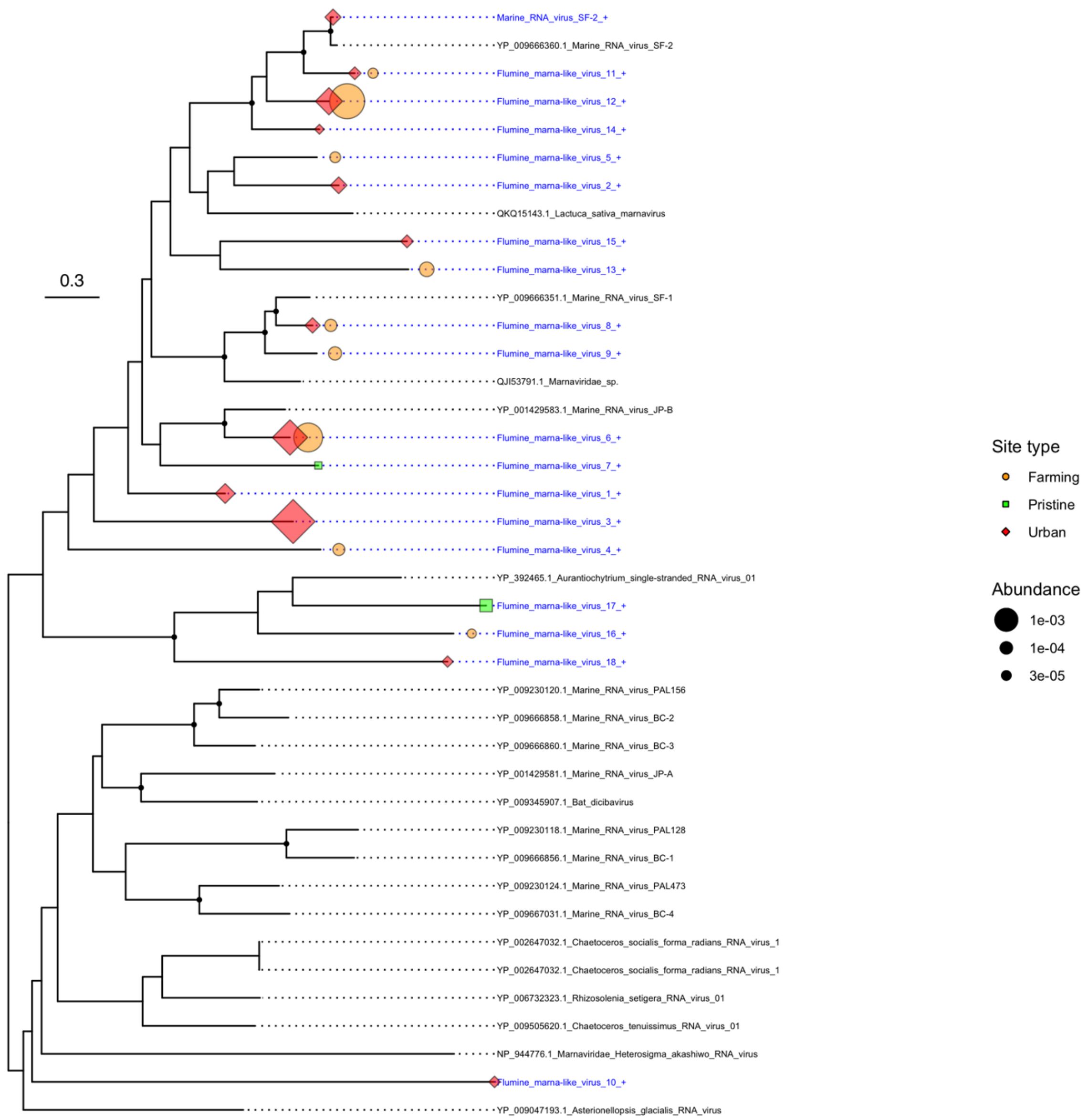
0.3
Marine_RNA_virus_SF-2_+
YP_009666360.1_Marine_RNA_virus_SF-2
Flumine_marna-like_virus_11_+
Flumine_marna-like_virus_12_+
Flumine_marna-like_virus_14_+
Flumine_marna-like_virus_5_+
Flumine_marna-like_virus_2_+
QKQ15143.1_Lactuca_sativa_marnavirus
Flumine_marna-like_virus_15_+
Flumine_marna-like_virus_13_+
YP_009666351.1_Marine_RNA_virus_SF-1
Flumine_marna-like_virus_8_+
Flumine_marna-like_virus_9_+
QJI53791.1_Marnaviridae_sp.
YP_001429583.1_Marine_RNA_virus_JP-B
Flumine_marna-like_virus_6_+
Flumine_marna-like_virus_7_+
Flumine_marna-like_virus_1_+
Flumine_marna-like_virus_3_+
Flumine_marna-like_virus_4_+
YP_392465.1_Aurantiochytrium_single-stranded_RNA_virus_01
Flumine_marna-like_virus_17_+
Flumine_marna-like_virus_16_+
Flumine_marna-like_virus_18_+
YP_009230120.1_Marine_RNA_virus_PAL156
YP_009666858.1_Marine_RNA_virus_BC-2
YP_009666860.1_Marine_RNA_virus_BC-3
YP_001429581.1_Marine_RNA_virus_JP-A
YP_009345907.1_Bat_dicibavirus
YP_009230118.1_Marine_RNA_virus_PAL128
YP_009666856.1_Marine_RNA_virus_BC-1
YP_009230124.1_Marine_RNA_virus_PAL473
YP_009667031.1_Marine_RNA_virus_BC-4
YP_002647032.1_Chaetoceros_socialis_forma_radians_RNA_virus_1
YP_002647032.1_Chaetoceros_socialis_forma_radians_RNA_virus_1
YP_006732323.1_Rhizosolenia_setigera_RNA_virus_01
YP_009505620.1_Chaetoceros_tenuissimus_RNA_virus_01
NP_944776.1_Marnaviridae_Heterosigma_akashiwo_RNA_virus
Flumine_marna-like_virus_10_+
YP_009047193.1_Asterionellopsis_glacialis_RNA_virus
Site type
Farming
Pristine
Urban
Abundance
1e-03
1e-04
3e-05

**Figure S10**

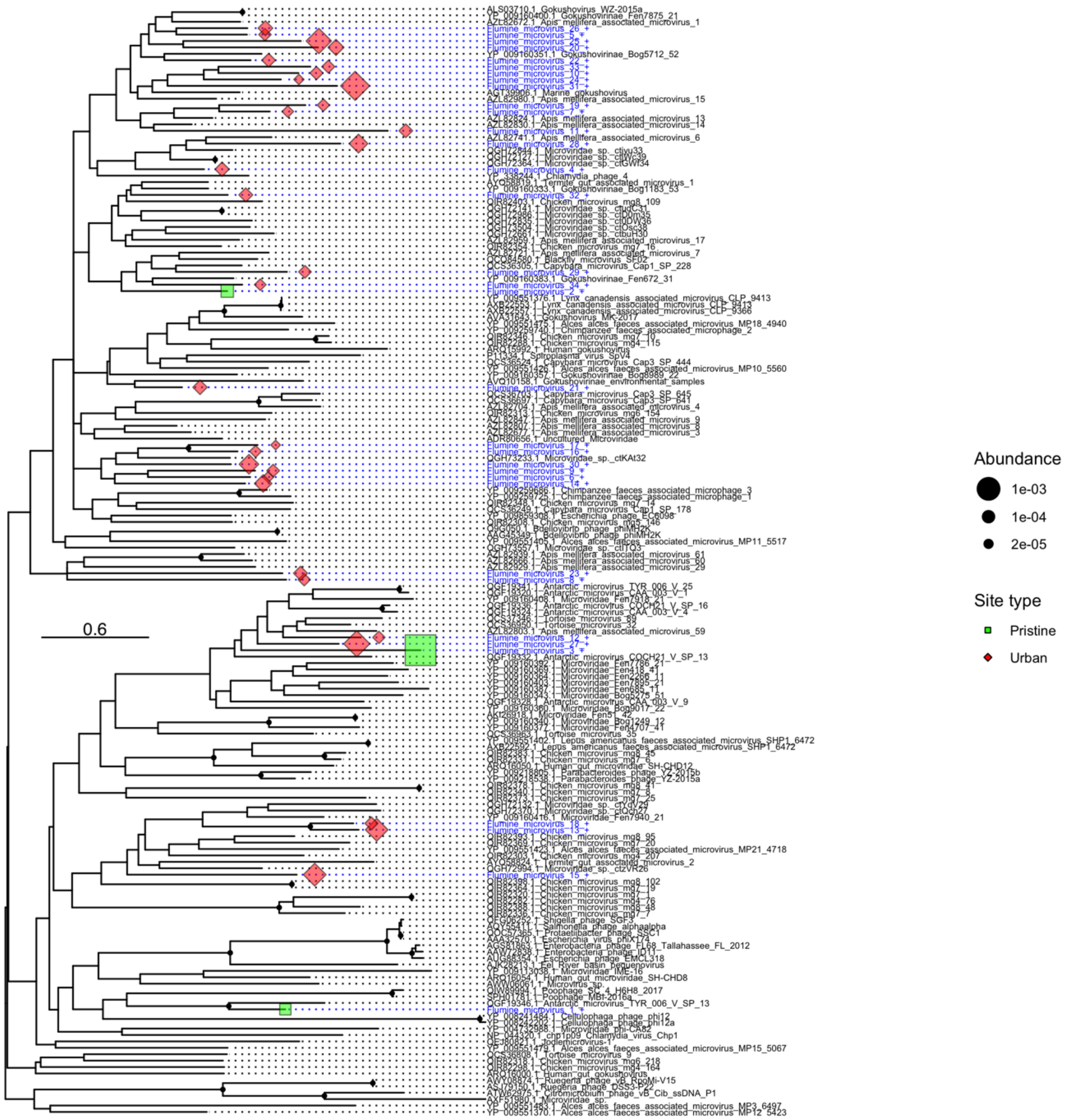


**Figure S11**

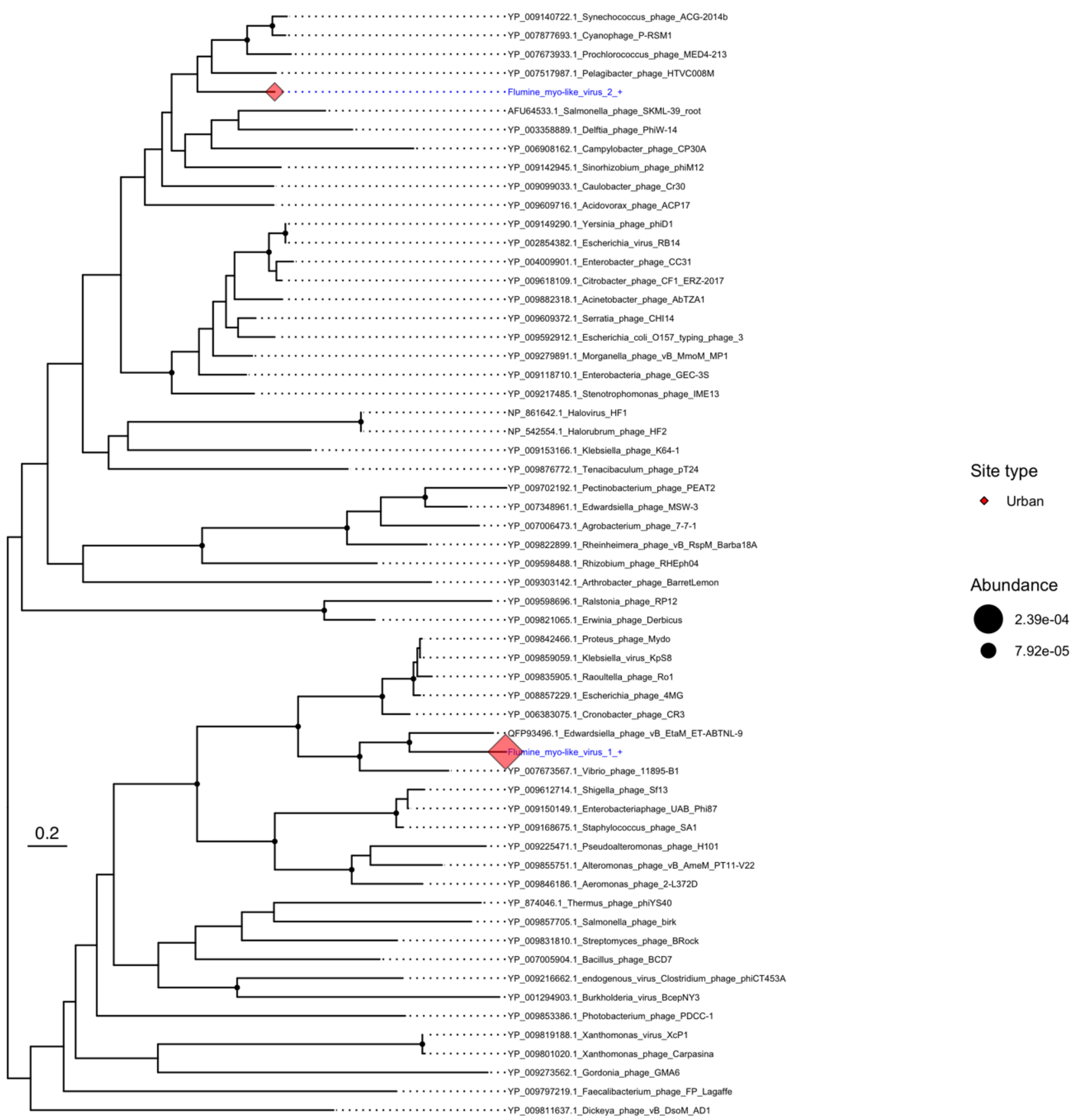
YP_009140722.1_Synechococcus_phage_ACG-2014b
YP_007877693.1_Cyanophage_P-RSM1
YP_007673933.1_Prochlorococcus_phage_MED4-213
YP_007517987.1_Pelagibacter_phage_HTVC008M
Flumine_myo-like_virus_2_+
AFU64533.1_Salmonella_phage_SKML-39_root
YP_003358889.1_Delftia_phage_PhiW-14
YP_006908162.1_Campylobacter_phage_CP30A
YP_009142945.1_Sinorhizobium_phage_phiM12
YP_009099033.1_Caulobacter_phage_Cr30
YP_009609716.1_Acidovorax_phage_ACP17
YP_009149290.1_Yersinia_phage_phiD1
YP_002854382.1_Escherichia_virus_RB14
YP_004009901.1_Enterobacter_phage_CC31
YP_009618109.1_Citrobacter_phage_CF1_ERZ-2017
YP_009882318.1_Acinetobacter_phage_AbTZA1
YP_009609372.1_Serratia_phage_CHI14
YP_009592912.1_Escherichia_coli_O157_typing_phage_3
YP_009279891.1_Morganella_phage_vB_MmoM_MP1
YP_009118710.1_Enterobacteria_phage_GEC-3S
YP_009217485.1_Stenotrophomonas_phage_IME13
NP_861642.1_Halovirus_HF1
NP_542554.1_Halorubrum_phage_HF2
YP_009153166.1_Klebsiella_phage_K64-1
YP_009876772.1_Tenacibaculum_phage_pT24
YP_009702192.1_Pectinobacterium_phage_PEAT2
YP_007348961.1_Edwardsiella_phage_MSW-3
YP_007006473.1_Agrobacterium_phage_7-7-1
YP_009822899.1_Rheinheimera_phage_vB_RspM_Barba18A
YP_009598488.1_Rhizobium_phage_RHEph04
YP_009303142.1_Arthrobacter_phage_BarretLemon
YP_009598696.1_Ralstonia_phage_RP12
YP_009821065.1_Erwinia_phage_Derbicus
YP_009842466.1_Proteus_phage_Mydo
YP_009859059.1_Klebsiella_virus_KpS8
YP_009835905.1_Raoultella_phage_Ro1
YP_008857229.1_Escherichia_phage_4MG
YP_006383075.1_Cronobacter_phage_CR3
QFP93496.1_Edwardsiella_phage_vB_EtaM_ET-ABTNL-9
Flumine_myo-like_virus_1_+
YP_007673567.1_Vibrio_phage_11895-B1
YP_009612714.1_Shigella_phage_Sf13
YP_009150149.1_Enterobacteriaphage_UAB_Phi87
YP_009168675.1_Staphylococcus_phage_SA1
YP_009225471.1_Pseudoalteromonas_phage_H101
YP_009855751.1_Alteromonas_phage_vB_AmeM_PT11-V22
YP_009846186.1_Aeromonas_phage_2-L372D
YP_874046.1_Thermus_phage_phiYS40
YP_009857705.1_Salmonella_phage_birk
YP_009831810.1_Streptomyces_phage_BRock
YP_007005904.1_Bacillus_phage_BCD7
YP_009216662.1_endogenous_virus_Clostridium_phage_phiCT453A
YP_001294903.1_Burkholderia_virus_BcepNY3
YP_009853386.1_Photobacterium_phage_PDCC-1
YP_009819188.1_Xanthomonas_virus_XcP1
YP_009801020.1_Xanthomonas_phage_Carpasina
YP_009273562.1_Gordonia_phage_GMA6
YP_009797219.1_Faecalibacterium_phage_FP_Lagaffe
YP_009811637.1_Dickeya_phage_vB_DsoM_AD1
0.2
Site type
Urban
Abundance
2.39e-04
7.92e-05

**Figure S12**

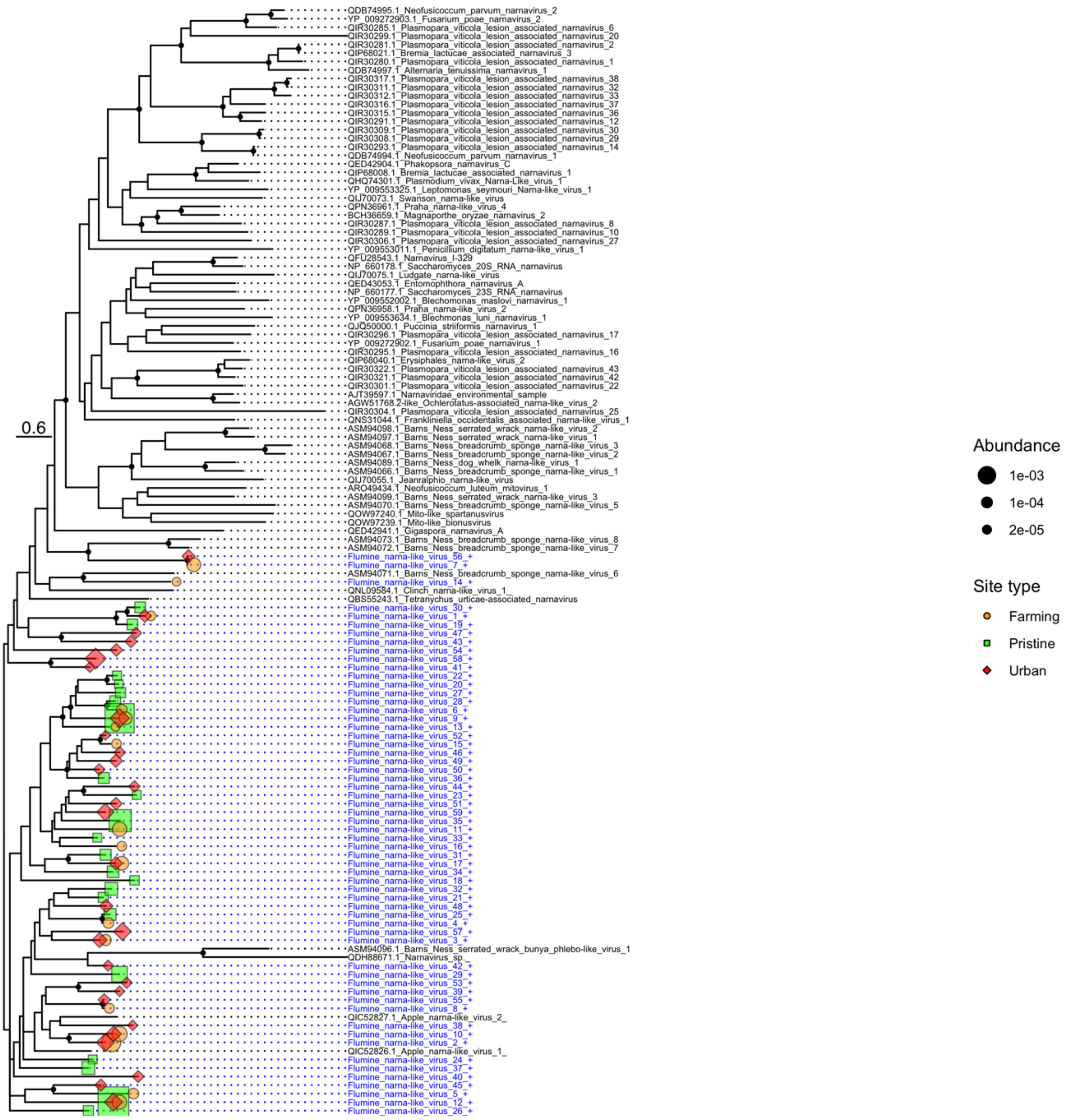
QDB74995.1_Neofusicoccum_parvum_narnavirus_2
YP_009272903.1_Fusarium_poae_narnavirus_2
QIR30285.1_Plasmopara_viticola_lesion_associated_narnavirus_6
QIR30299.1_Plasmopara_viticola_lesion_associated_narnavirus_20
QIR30281.1_Plasmopara_viticola_lesion_associated_narnavirus_2
QIP68021.1_Bremia_lactucae_associated_narnavirus_3
QIR30280.1_Plasmopara_viticola_lesion_associated_narnavirus_1
QDB74997.1_Alternaria_tenuissima_narnavirus_1
QIR30317.1_Plasmopara_viticola_lesion_associated_narnavirus_38
QIR30311.1_Plasmopara_viticola_lesion_associated_narnavirus_32
QIR30312.1_Plasmopara_viticola_lesion_associated_narnavirus_33
QIR30316.1_Plasmopara_viticola_lesion_associated_narnavirus_37
QIR30315.1_Plasmopara_viticola_lesion_associated_narnavirus_36
QIR30291.1_Plasmopara_viticola_lesion_associated_narnavirus_12
QIR30309.1_Plasmopara_viticola_lesion_associated_narnavirus_30
QIR30308.1_Plasmopara_viticola_lesion_associated_narnavirus_29
QIR30293.1_Plasmopara_viticola_lesion_associated_narnavirus_14
QDB74994.1_Neofusicoccum_parvum_narnavirus_1
QED42904.1_Phakopsora_narnavirus_C
QIP68008.1_Bremia_lactucae_associated_narnavirus_1
QHQ74301.1_Plasmodium_vivax_Narna-Like_virus_1
YP_009553325.1_Leptomonas_seymouri_Narna-like_virus_1
QIJ70073.1_Swanson_narna-like_virus
QPN36961.1_Praha_narna-like_virus_4
BCH36659.1_Magnaporthe_oryzae_narnavirus_2
QIR30287.1_Plasmopara_viticola_lesion_associated_narnavirus_8
QIR30289.1_Plasmopara_viticola_lesion_associated_narnavirus_10
QIR30306.1_Plasmopara_viticola_lesion_associated_narnavirus_27
YP_009553011.1_Penicillium_digitatum_narna-like_virus_1
QFU28543.1_Narnavirus_I-329
NP_660178.1_Saccharomyces_20S_RNA_narnavirus
QIJ70075.1_Ludgate_narna-like_virus
QED43053.1_Entomophthora_narnavirus_A
NP_660177.1_Saccharomyces_23S_RNA_narnavirus
YP_009552002.1_Blechomonas_maslovi_narnavirus_1
QPN36958.1_Praha_narna-like_virus_2
YP_009553634.1_Blechmonas_luni_narnavirus_1
QJQ50000.1_Puccinia_striiformis_narnavirus_1
QIR30296.1_Plasmopara_viticola_lesion_associated_narnavirus_17
YP_009272902.1_Fusarium_poae_narnavirus_1
QIR30295.1_Plasmopara_viticola_lesion_associated_narnavirus_16
QIP68040.1_Erysiphales_narna-like_virus_2
QIR30322.1_Plasmopara_viticola_lesion_associated_narnavirus_43
QIR30321.1_Plasmopara_viticola_lesion_associated_narnavirus_42
QIR30301.1_Plasmopara_viticola_lesion_associated_narnavirus_22
AJT39597.1_Narnaviridae_environmental_sample
AGW51768.2-like_Ochlerotatus-associated_narna-like_virus_2
QIR30304.1_Plasmopara_viticola_lesion_associated_narnavirus_25
QNS31044.1_Frankliniella_occidentalis_associated_narna-like_virus_1
ASM94098.1_Barns_Ness_serrated_wrack_narna-like_virus_2
ASM94097.1_Barns_Ness_serrated_wrack_narna-like_virus_1
ASM94068.1_Barns_Ness_breadcrumb_sponge_narna-like_virus_3
ASM94067.1_Barns_Ness_breadcrumb_sponge_narna-like_virus_2
ASM94089.1_Barns_Ness_dog_whelk_narna-like_virus_1
ASM94066.1_Barns_Ness_breadcrumb_sponge_narna-like_virus_1
QIJ70055.1_Jeanralphio_narna-like_virus
ARO49434.1_Neofusicoccum_luteum_mitovirus_1
ASM94099.1_Barns_Ness_serrated_wrack_narna-like_virus_3
ASM94070.1_Barns_Ness_breadcrumb_sponge_narna-like_virus_5
QOW97240.1_Mito-like_spartanusvirus
QOW97239.1_Mito-like_bionusvirus
QED42941.1_Gigaspora_narnavirus_A
ASM94073.1_Barns_Ness_breadcrumb_sponge_narna-like_virus_8
ASM94072.1_Barns_Ness_breadcrumb_sponge_narna-like_virus_7
Flumine_narna-like_virus_56_+
Flumine_narna-like_virus_7_+
ASM94071.1_Barns_Ness_breadcrumb_sponge_narna-like_virus_6
Flumine_narna-like_virus_14_+
QNL09584.1_Clinch_narna-like_virus_1_
QBS55243.1_Tetranychus_urticae-associated_narnavirus
Flumine_narna-like_virus_30_+
Flumine_narna-like_virus_1_+
Flumine_narna-like_virus_19_+
Flumine_narna-like_virus_47_+
Flumine_narna-like_virus_43_+
Flumine_narna-like_virus_54_+
Flumine_narna-like_virus_58_+
Flumine_narna-like_virus_41_+
Flumine_narna-like_virus_22_+
Flumine_narna-like_virus_20_+
Flumine_narna-like_virus_27_+
Flumine_narna-like_virus_28_+
Flumine_narna-like_virus_6_+
Flumine_narna-like_virus_9_+
Flumine_narna-like_virus_13_+
Flumine_narna-like_virus_52_+
Flumine_narna-like_virus_15_+
Flumine_narna-like_virus_46_+
Flumine_narna-like_virus_49_+
Flumine_narna-like_virus_50_+
Flumine_narna-like_virus_36_+
Flumine_narna-like_virus_44_+
Flumine_narna-like_virus_23_+
Flumine_narna-like_virus_51_+
Flumine_narna-like_virus_59_+
Flumine_narna-like_virus_35_+
Flumine_narna-like_virus_11_+
Flumine_narna-like_virus_33_+
Flumine_narna-like_virus_16_+
Flumine_narna-like_virus_31_+
Flumine_narna-like_virus_17_+
Flumine_narna-like_virus_34_+
Flumine_narna-like_virus_18_+
Flumine_narna-like_virus_32_+
Flumine_narna-like_virus_21_+
Flumine_narna-like_virus_48_+
Flumine_narna-like_virus_25_+
Flumine_narna-like_virus_4_+
Flumine_narna-like_virus_57_+
Flumine_narna-like_virus_3_+
ASM94096.1_Barns_Ness_serrated_wrack_bunya_phlebo-like_virus_1
QDH88671.1_Narnavirus_sp._
Flumine_narna-like_virus_42_+
Flumine_narna-like_virus_29_+
Flumine_narna-like_virus_53_+
Flumine_narna-like_virus_39_+
Flumine_narna-like_virus_55_+
Flumine_narna-like_virus_8_+
QIC52827.1_Apple_narna-like_virus_2_
Flumine_narna-like_virus_38_+
Flumine_narna-like_virus_10_+
Flumine_narna-like_virus_2_+
QIC52826.1_Apple_narna-like_virus_1_
Flumine_narna-like_virus_24_+
Flumine_narna-like_virus_37_+
Flumine_narna-like_virus_40_+
Flumine_narna-like_virus_45_+
Flumine_narna-like_virus_5_+
Flumine_narna-like_virus_12_+
Flumine_narna-like_virus_26_+
0.6
Abundance
1e-03
1e-04
2e-05
Site type
Farming
Pristine
Urban

**Figure S13**

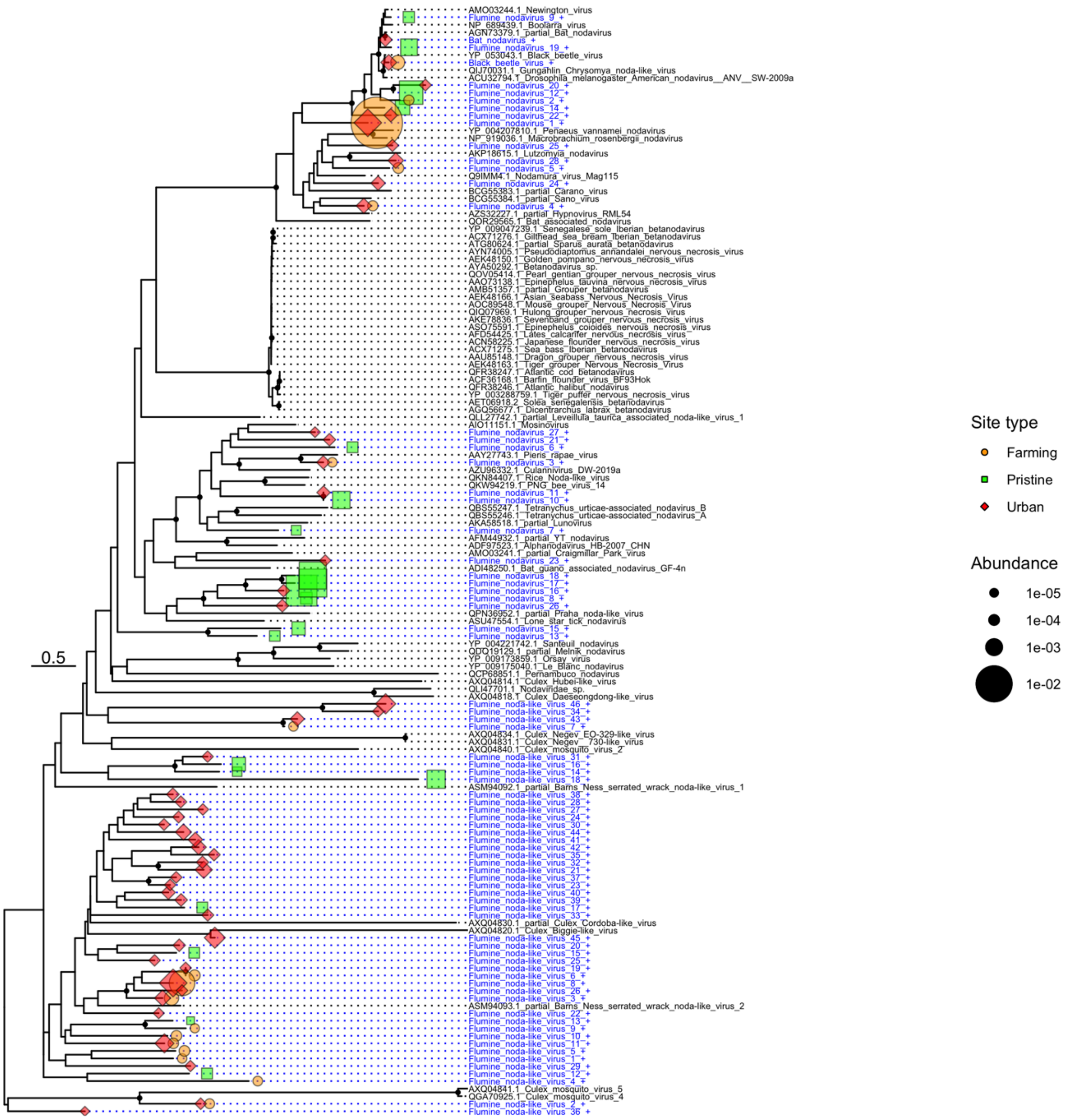
0.5
Site type
Farming
Pristine
Urban
Abundance
1e-05
1e-04
1e-03
1e-02

**Figure S14**

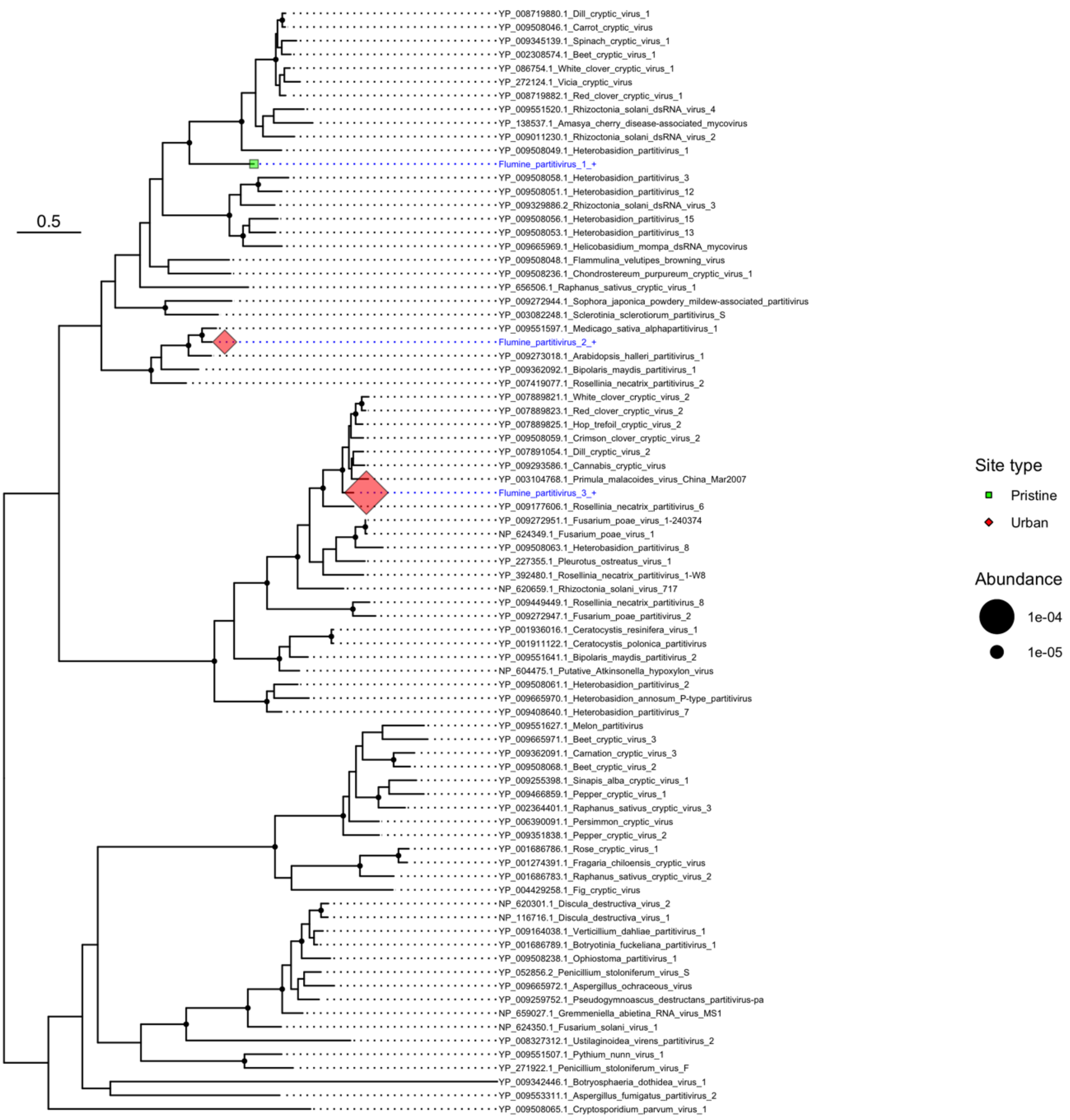

0.5
YP_008719880.1_Dill_cryptic_virus_1
YP_009508046.1_Carrot_cryptic_virus
YP_009345139.1_Spinach_cryptic_virus_1
YP_002308574.1_Beet_cryptic_virus_1
YP_086754.1_White_clover_cryptic_virus_1
YP_272124.1_Vicia_cryptic_virus
YP_008719882.1_Red_clover_cryptic_virus_1
YP_009551520.1_Rhizoctonia_solani_dsRNA_virus_4
YP_138537.1_Amasya_cherry_disease-associated_mycovirus
YP_009011230.1_Rhizoctonia_solani_dsRNA_virus_2
YP_009508049.1_Heterobasidion_partitivirus_1
Flumine_partitivirus_1_+
YP_009508058.1_Heterobasidion_partitivirus_3
YP_009508051.1_Heterobasidion_partitivirus_12
YP_009329886.2_Rhizoctonia_solani_dsRNA_virus_3
YP_009508056.1_Heterobasidion_partitivirus_15
YP_009508053.1_Heterobasidion_partitivirus_13
YP_009665969.1_Helicobasidium_mompa_dsRNA_mycovirus
YP_009508048.1_Flammulina_velutipes_browning_virus
YP_009508236.1_Chondrostereum_purpureum_cryptic_virus_1
YP_656506.1_Raphanus_sativus_cryptic_virus_1
YP_009272944.1_Sophora_japonica_powdery_mildew-associated_partitivirus
YP_003082248.1_Sclerotinia_sclerotiorum_partitivirus_S
YP_009551597.1_Medicago_sativa_alphapartitivirus_1
Flumine_partitivirus_2_+
YP_009273018.1_Arabidopsis_halleri_partitivirus_1
YP_009362092.1_Bipolaris_maydis_partitivirus_1
YP_007419077.1_Rosellinia_necatrix_partitivirus_2
YP_007889821.1_White_clover_cryptic_virus_2
YP_007889823.1_Red_clover_cryptic_virus_2
YP_007889825.1_Hop_trefoil_cryptic_virus_2
YP_009508059.1_Crimson_clover_cryptic_virus_2
YP_007891054.1_Dill_cryptic_virus_2
YP_009293586.1_Cannabis_cryptic_virus
YP_003104768.1_Primula_malacoides_virus_China_Mar2007
Flumine_partitivirus_3_+
YP_009177606.1_Rosellinia_necatrix_partitivirus_6
YP_009272951.1_Fusarium_poae_virus_1-240374
NP_624349.1_Fusarium_poae_virus_1
YP_009508063.1_Heterobasidion_partitivirus_8
YP_227355.1_Pleurotus_ostreatus_virus_1
YP_392480.1_Rosellinia_necatrix_partitivirus_1-W8
NP_620659.1_Rhizoctonia_solani_virus_717
YP_009449449.1_Rosellinia_necatrix_partitivirus_8
YP_009272947.1_Fusarium_poae_partitivirus_2
YP_001936016.1_Ceratocystis_resinifera_virus_1
YP_001911122.1_Ceratocystis_polonica_partitivirus
YP_009551641.1_Bipolaris_maydis_partitivirus_2
NP_604475.1_Putative_Atkinsonella_hypoxylon_virus
YP_009508061.1_Heterobasidion_partitivirus_2
YP_009665970.1_Heterobasidion_annosum_P-type_partitivirus
YP_009408640.1_Heterobasidion_partitivirus_7
YP_009551627.1_Melon_partitivirus
YP_009665971.1_Beet_cryptic_virus_3
YP_009362091.1_Carnation_cryptic_virus_3
YP_009508068.1_Beet_cryptic_virus_2
YP_009255398.1_Sinapis_alba_cryptic_virus_1
YP_009466859.1_Pepper_cryptic_virus_1
YP_002364401.1_Raphanus_sativus_cryptic_virus_3
YP_006390091.1_Persimmon_cryptic_virus
YP_009351838.1_Pepper_cryptic_virus_2
YP_001686786.1_Rose_cryptic_virus_1
YP_001274391.1_Fragaria_chiloensis_cryptic_virus
YP_001686783.1_Raphanus_sativus_cryptic_virus_2
YP_004429258.1_Fig_cryptic_virus
NP_620301.1_Discula_destructiva_virus_2
NP_116716.1_Discula_destructiva_virus_1
YP_009164038.1_Verticillium_dahliae_partitivirus_1
YP_001686789.1_Botryotinia_fuckeliana_partitivirus_1
YP_009508238.1_Ophiostoma_partitivirus_1
YP_052856.2_Penicillium_stoloniferum_virus_S
YP_009665972.1_Aspergillus_ochraceous_virus
YP_009259752.1_Pseudogymnoascus_destructans_partitivirus-pa
NP_659027.1_Gremmeniella_abietina_RNA_virus_MS1
NP_624350.1_Fusarium_solani_virus_1
YP_008327312.1_Ustilaginoidea_virens_partitivirus_2
YP_009551507.1_Pythium_nunn_virus_1
YP_271922.1_Penicillium_stoloniferum_virus_F
YP_009342446.1_Botryosphaeria_dothidea_virus_1
YP_009553311.1_Aspergillus_fumigatus_partitivirus_2
YP_009508065.1_Cryptosporidium_parvum_virus_1
Site type
Pristine
Urban
Abundance
1e-04
1e-05


**Figure S15**

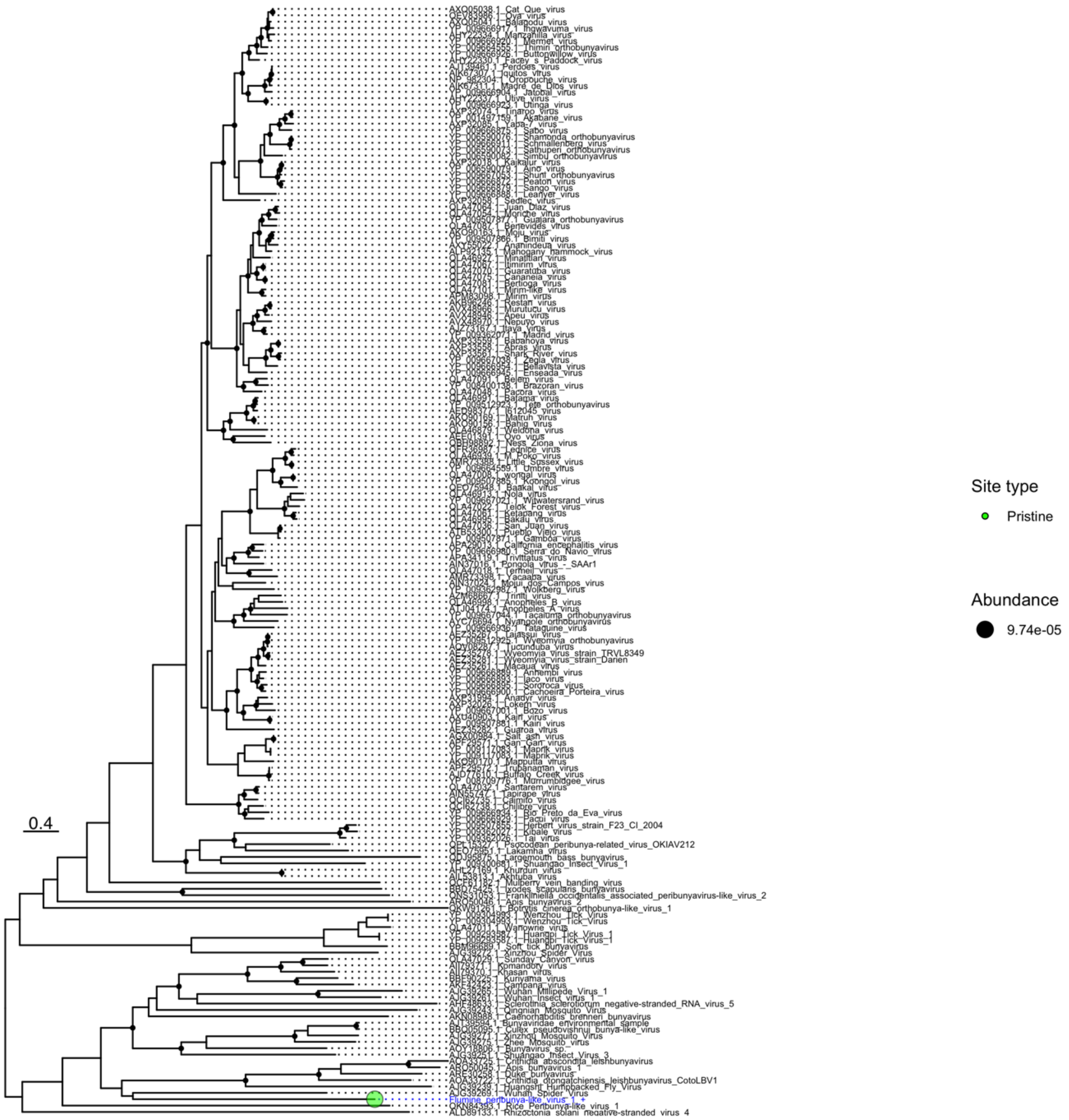


**Figure S16**

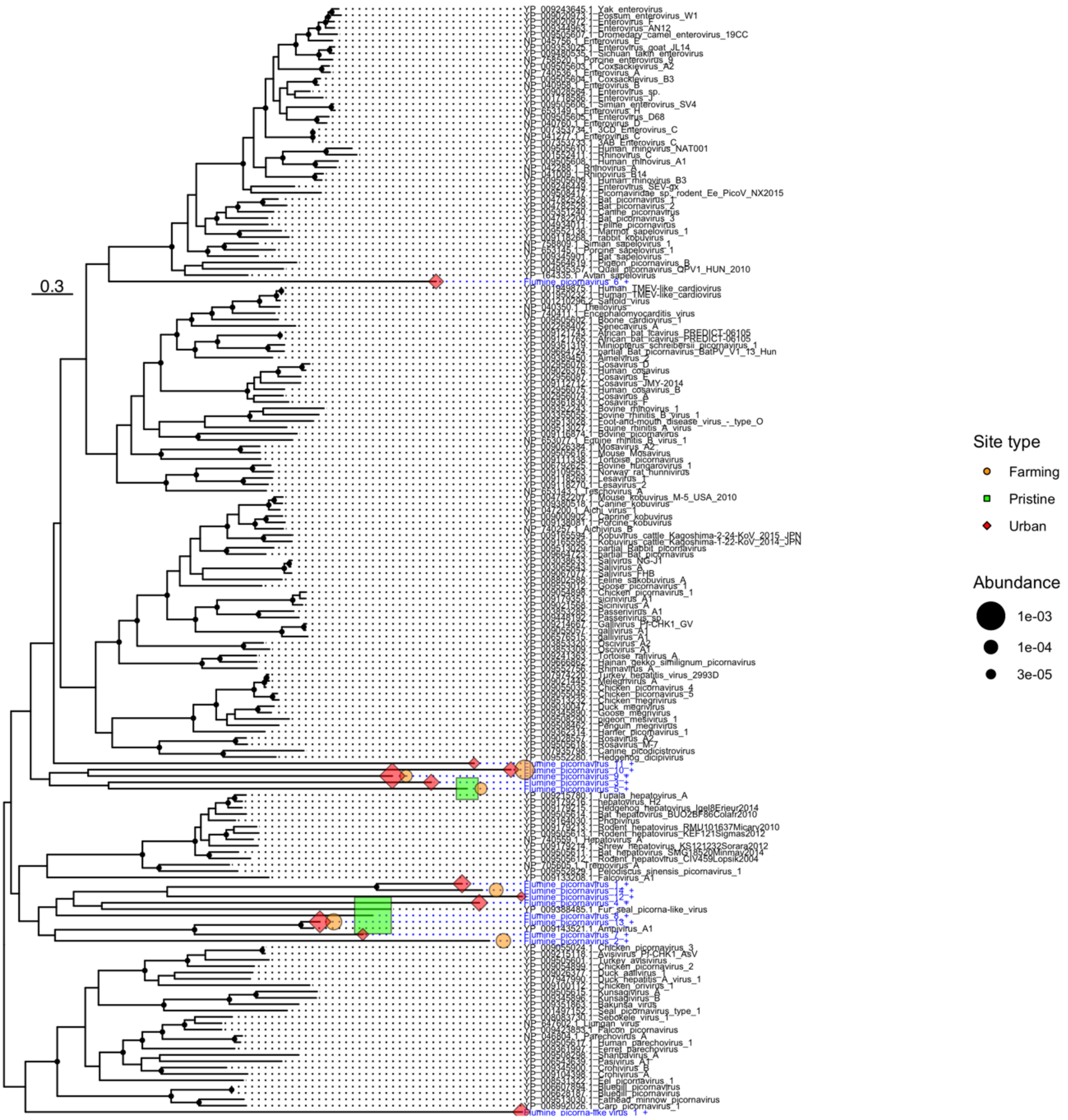
0.3
Site type
Farming
Pristine
Urban
Abundance
1e-03
1e-04
3e-05

**Figure S17**

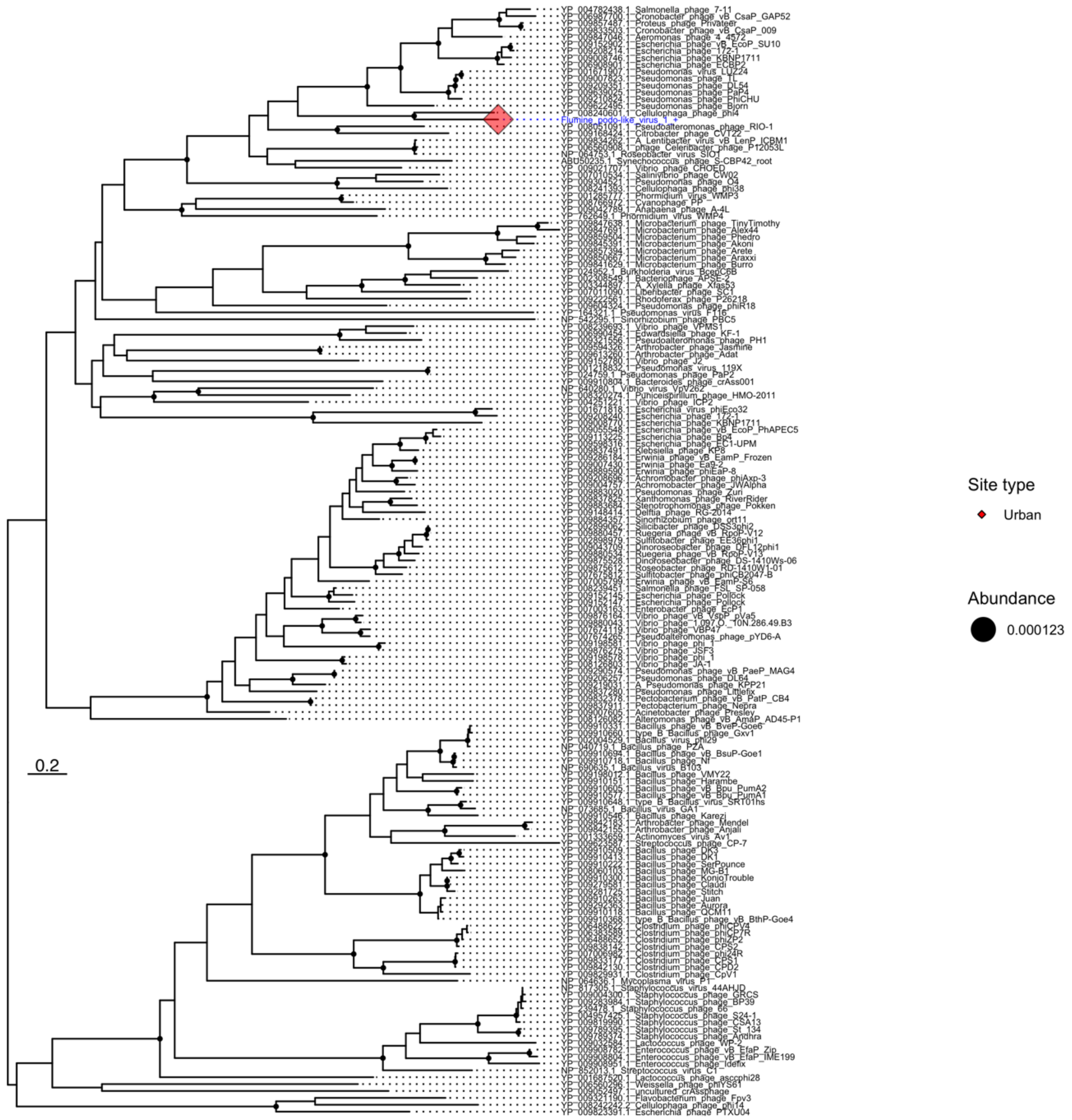


**Figure S18**

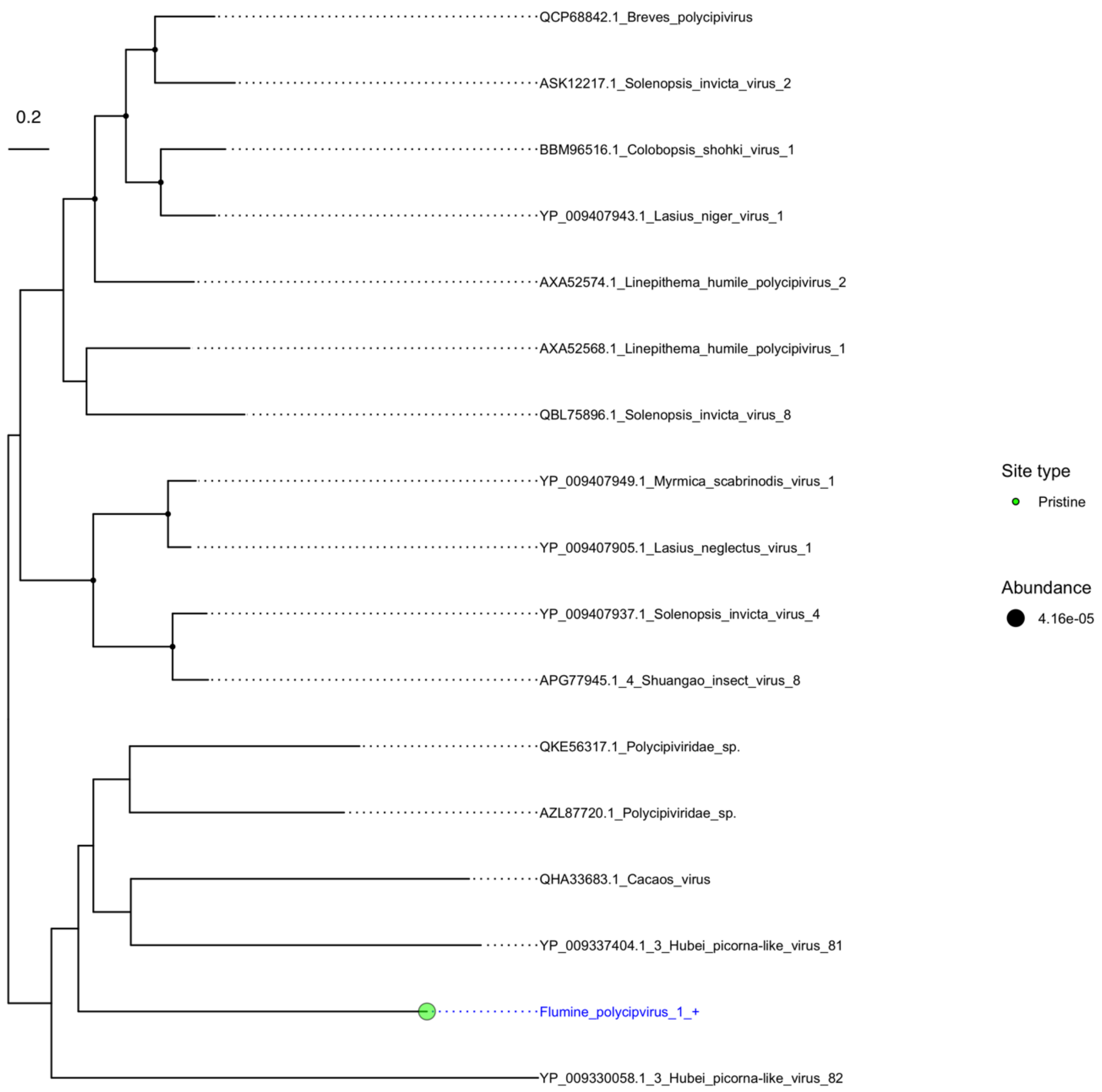

0.2
QCP68842.1_Breves_polycipivirus
ASK12217.1_Solenopsis_invicta_virus_2
BBM96516.1_Colobopsis_shohki_virus_1
YP_009407943.1_Lasius_niger_virus_1
AXA52574.1_Linepithema_humile_polycipivirus_2
AXA52568.1_Linepithema_humile_polycipivirus_1
QBL75896.1_Solenopsis_invicta_virus_8
YP_009407949.1_Myrmica_scabrinodis_virus_1
YP_009407905.1_Lasius_neglectus_virus_1
YP_009407937.1_Solenopsis_invicta_virus_4
APG77945.1_4_Shuangao_insect_virus_8
QKE56317.1_Polycipiviridae_sp.
AZL87720.1_Polycipiviridae_sp.
QHA33683.1_Cacaos_virus
YP_009337404.1_3_Hubei_picorna-like_virus_81
Flumine_polycipvirus_1_+
YP_009330058.1_3_Hubei_picorna-like_virus_82
Site type
Pristine
Abundance
4.16e-05


**Figure S19**

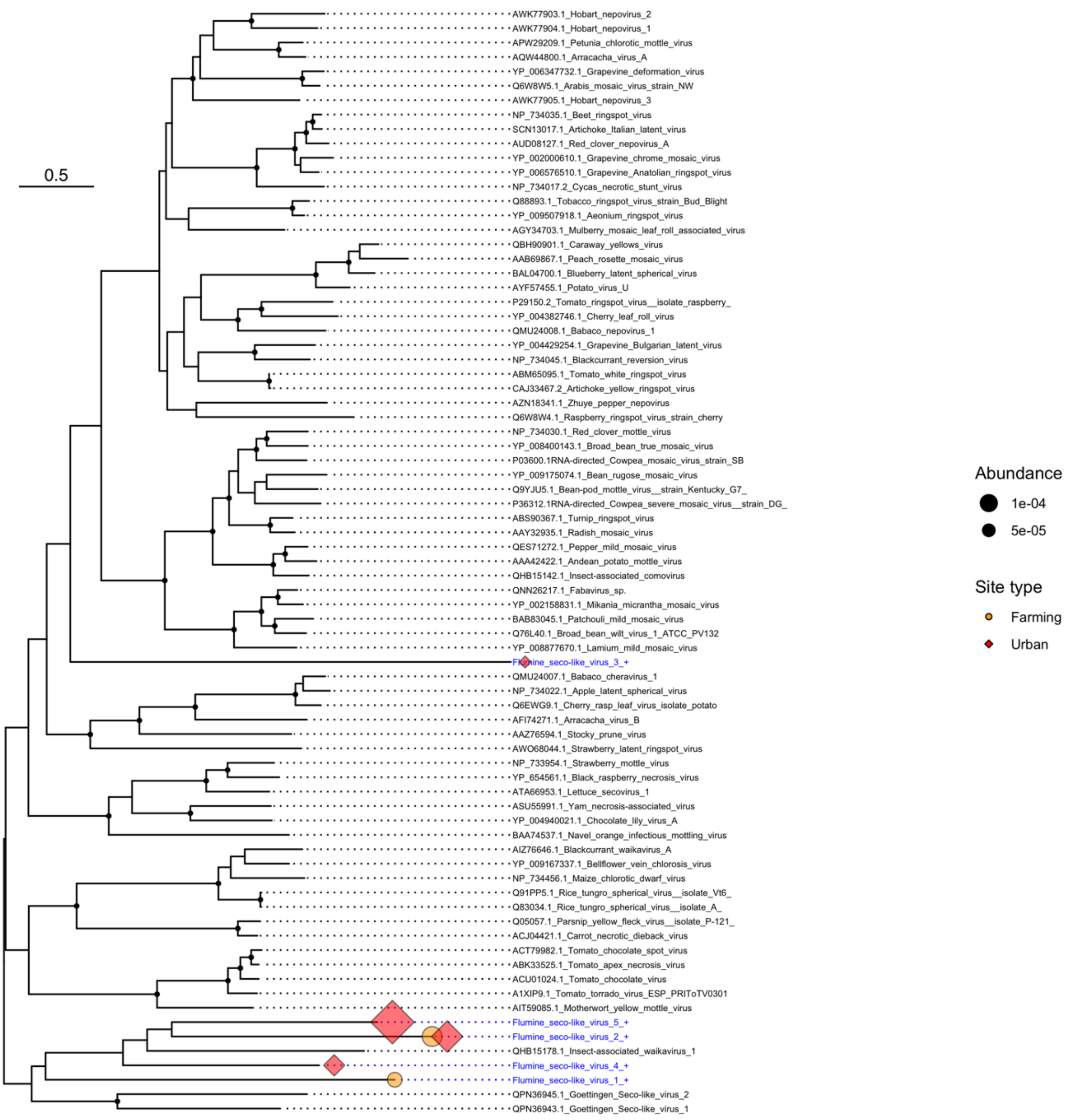

0.5
AWK77903.1_Hobart_nepovirus_2
AWK77904.1_Hobart_nepovirus_1
APW29209.1_Petunia_chlorotic_mottle_virus
AQW44800.1_Arracacha_virus_A
YP_006347732.1_Grapevine_deformation_virus
Q6W8W5.1_Arabis_mosaic_virus_strain_NW
AWK77905.1_Hobart_nepovirus_3
NP_734035.1_Beet_ringspot_virus
SCN13017.1_Artichoke_Italian_latent_virus
AUD08127.1_Red_clover_nepovirus_A
YP_002000610.1_Grapevine_chrome_mosaic_virus
YP_006576510.1_Grapevine_Anatolian_ringspot_virus
NP_734017.2_Cycas_necrotic_stunt_virus
Q88893.1_Tobacco_ringspot_virus_strain_Bud_Blight
YP_009507918.1_Aeonium_ringspot_virus
AGY34703.1_Mulberry_mosaic_leaf_roll_associated_virus
QBH90901.1_Caraway_yellows_virus
AAB69867.1_Peach_rosette_mosaic_virus
BAL04700.1_Blueberry_latent_spherical_virus
AYF57455.1_Potato_virus_U
P29150.2_Tomato_ringspot_virus__isolate_raspberry_
YP_004382746.1_Cherry_leaf_roll_virus
QMU24008.1_Babaco_nepovirus_1
YP_004429254.1_Grapevine_Bulgarian_latent_virus
NP_734045.1_Blackcurrant_reversion_virus
ABM65095.1_Tomato_white_ringspot_virus
CAJ33467.2_Artichoke_yellow_ringspot_virus
AZN18341.1_Zhuye_pepper_nepovirus
Q6W8W4.1_Raspberry_ringspot_virus_strain_cherry
NP_734030.1_Red_clover_mottle_virus
YP_008400143.1_Broad_bean_true_mosaic_virus
P03600.1RNA-directed_Cowpea_mosaic_virus_strain_SB
YP_009175074.1_Bean_rugose_mosaic_virus
Q9YJU5.1_Bean-pod_mottle_virus__strain_Kentucky_G7_
P36312.1RNA-directed_Cowpea_severe_mosaic_virus__strain_DG_
ABS90367.1_Turnip_ringspot_virus
AAY32935.1_Radish_mosaic_virus
QES71272.1_Pepper_mild_mosaic_virus
AAA42422.1_Andean_potato_mottle_virus
QHB15142.1_Insect-associated_comovirus
QNN26217.1_Fabavirus_sp.
YP_002158831.1_Mikania_micrantha_mosaic_virus
BAB83045.1_Patchouli_mild_mosaic_virus
Q76L40.1_Broad_bean_wilt_virus_1_ATCC_PV132
YP_008877670.1_Lamium_mild_mosaic_virus
Flumine_seco-like_virus_3_+
QMU24007.1_Babaco_cheravirus_1
NP_734022.1_Apple_latent_spherical_virus
Q6EWG9.1_Cherry_rasp_leaf_virus_isolate_potato
AFI74271.1_Arracacha_virus_B
AAZ76594.1_Stocky_prune_virus
AWO68044.1_Strawberry_latent_ringspot_virus
NP_733954.1_Strawberry_mottle_virus
YP_654561.1_Black_raspberry_necrosis_virus
ATA66953.1_Lettuce_secovirus_1
ASU55991.1_Yam_necrosis-associated_virus
YP_004940021.1_Chocolate_lily_virus_A
BAA74537.1_Navel_orange_infectious_mottling_virus
AIZ76646.1_Blackcurrant_waikavirus_A
YP_009167337.1_Bellflower_vein_chlorosis_virus
NP_734456.1_Maize_chlorotic_dwarf_virus
Q91PP5.1_Rice_tungro_spherical_virus__isolate_Vt6_
Q83034.1_Rice_tungro_spherical_virus__isolate_A_
Q05057.1_Parsnip_yellow_fleck_virus__isolate_P-121_
ACJ04421.1_Carrot_necrotic_dieback_virus
ACT79982.1_Tomato_chocolate_spot_virus
ABK33525.1_Tomato_apex_necrosis_virus
ACU01024.1_Tomato_chocolate_virus
A1XIP9.1_Tomato_torrado_virus_ESP_PRIToTV0301
AIT59085.1_Motherwort_yellow_mottle_virus
Flumine_seco-like_virus_5_+
Flumine_seco-like_virus_2_+
QHB15178.1_Insect-associated_waikavirus_1
Flumine_seco-like_virus_4_+
Flumine_seco-like_virus_1_+
QPN36945.1_Goettingen_Seco-like_virus_2
QPN36943.1_Goettingen_Seco-like_virus_1
Abundance
1e-04
5e-05
Site type
Farming
Urban


### Figure S20

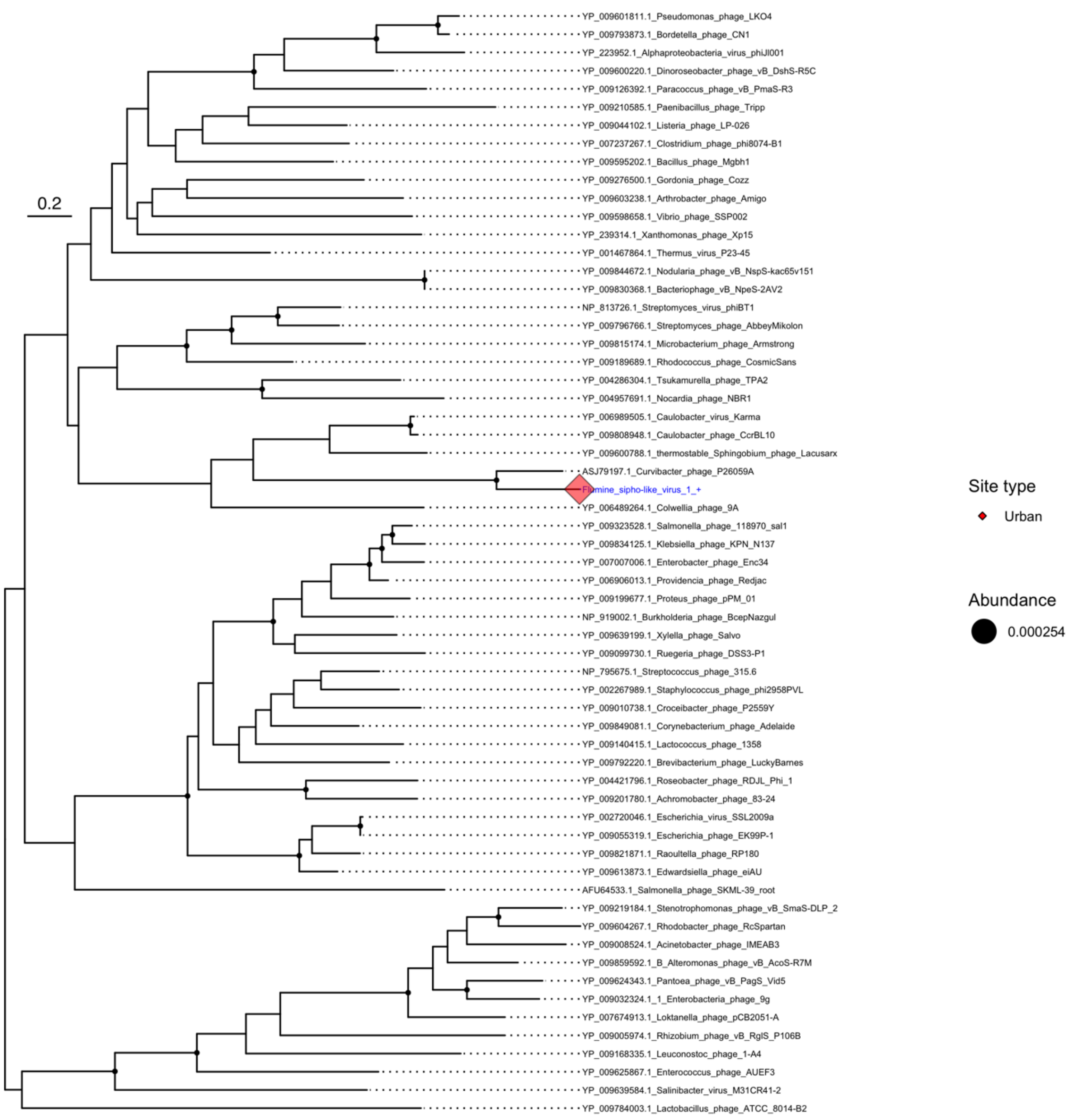

0.2
YP_009601811.1_Pseudomonas_phage_LKO4
YP_009793873.1_Bordetella_phage_CN1
YP_223952.1_Alphaproteobacteria_virus_phiJl001
YP_009600220.1_Dinoroseobacter_phage_vB_DshS-R5C
YP_009126392.1_Paracoccus_phage_vB_PmaS-R3
YP_009210585.1_Paenibacillus_phage_Tripp
YP_009044102.1_Listeria_phage_LP-026
YP_007237267.1_Clostridium_phage_phi8074-B1
YP_009595202.1_Bacillus_phage_Mgbh1
YP_009276500.1_Gordonia_phage_Cozz
YP_009603238.1_Arthrobacter_phage_Amigo
YP_009598658.1_Vibrio_phage_SSP002
YP_239314.1_Xanthomonas_phage_Xp15
YP_001467864.1_Thermus_virus_P23-45
YP_009844672.1_Nodularia_phage_vB_NspS-kac65v151
YP_009830368.1_Bacteriophage_vB_NpeS-2AV2
NP_813726.1_Streptomyces_virus_phiBT1
YP_009796766.1_Streptomyces_phage_AbbeyMikolon
YP_009815174.1_Microbacterium_phage_Armstrong
YP_009189689.1_Rhodococcus_phage_CosmicSans
YP_004286304.1_Tsukamurella_phage_TPA2
YP_004957691.1_Nocardia_phage_NBR1
YP_006989505.1_Caulobacter_virus_Karma
YP_009808948.1_Caulobacter_phage_CcrBL10
YP_009600788.1_thermostable_Sphingobium_phage_Lacusarx
ASJ79197.1_Curvibacter_phage_P26059A
Flumine_sipho-like_virus_1_+
YP_006489264.1_Colwellia_phage_9A
YP_009323528.1_Salmonella_phage_118970_sal1
YP_009834125.1_Klebsiella_phage_KPN_N137
YP_007007006.1_Enterobacter_phage_Enc34
YP_006906013.1_Providencia_phage_Redjac
YP_009199677.1_Proteus_phage_pPM_01
NP_919002.1_Burkholderia_phage_BcepNazgul
YP_009639199.1_Xylella_phage_Salvo
YP_009099730.1_Ruegeria_phage_DSS3-P1
NP_795675.1_Streptococcus_phage_315.6
YP_002267989.1_Staphylococcus_phage_phi2958PVL
YP_009010738.1_Croceibacter_phage_P2559Y
YP_009849081.1_Corynebacterium_phage_Adelaide
YP_009140415.1_Lactococcus_phage_1358
YP_009792220.1_Brevibacterium_phage_LuckyBarnes
YP_004421796.1_Roseobacter_phage_RDJL_Phi_1
YP_009201780.1_Achromobacter_phage_83-24
YP_002720046.1_Escherichia_virus_SSL2009a
YP_009055319.1_Escherichia_phage_EK99P-1
YP_009821871.1_Raoultella_phage_RP180
YP_009613873.1_Edwardsiella_phage_eiAU
AFU64533.1_Salmonella_phage_SKML-39_root
YP_009219184.1_Stenotrophomonas_phage_vB_SmaS-DLP_2
YP_009604267.1_Rhodobacter_phage_RcSpartan
YP_009008524.1_Acinetobacter_phage_IMEAB3
YP_009859592.1_B_Alteromonas_phage_vB_AcoS-R7M
YP_009624343.1_Pantoea_phage_vB_PagS_Vid5
YP_009032324.1_1_Enterobacteria_phage_9g
YP_007674913.1_Loktanella_phage_pCB2051-A
YP_009005974.1_Rhizobium_phage_vB_RglS_P106B
YP_009168335.1_Leuconostoc_phage_1-A4
YP_009625867.1_Enterococcus_phage_AUEF3
YP_009639584.1_Salinibacter_virus_M31CR41-2
YP_009784003.1_Lactobacillus_phage_ATCC_8014-B2
Site type
Urban
Abundance
0.000254


**Figure S21**

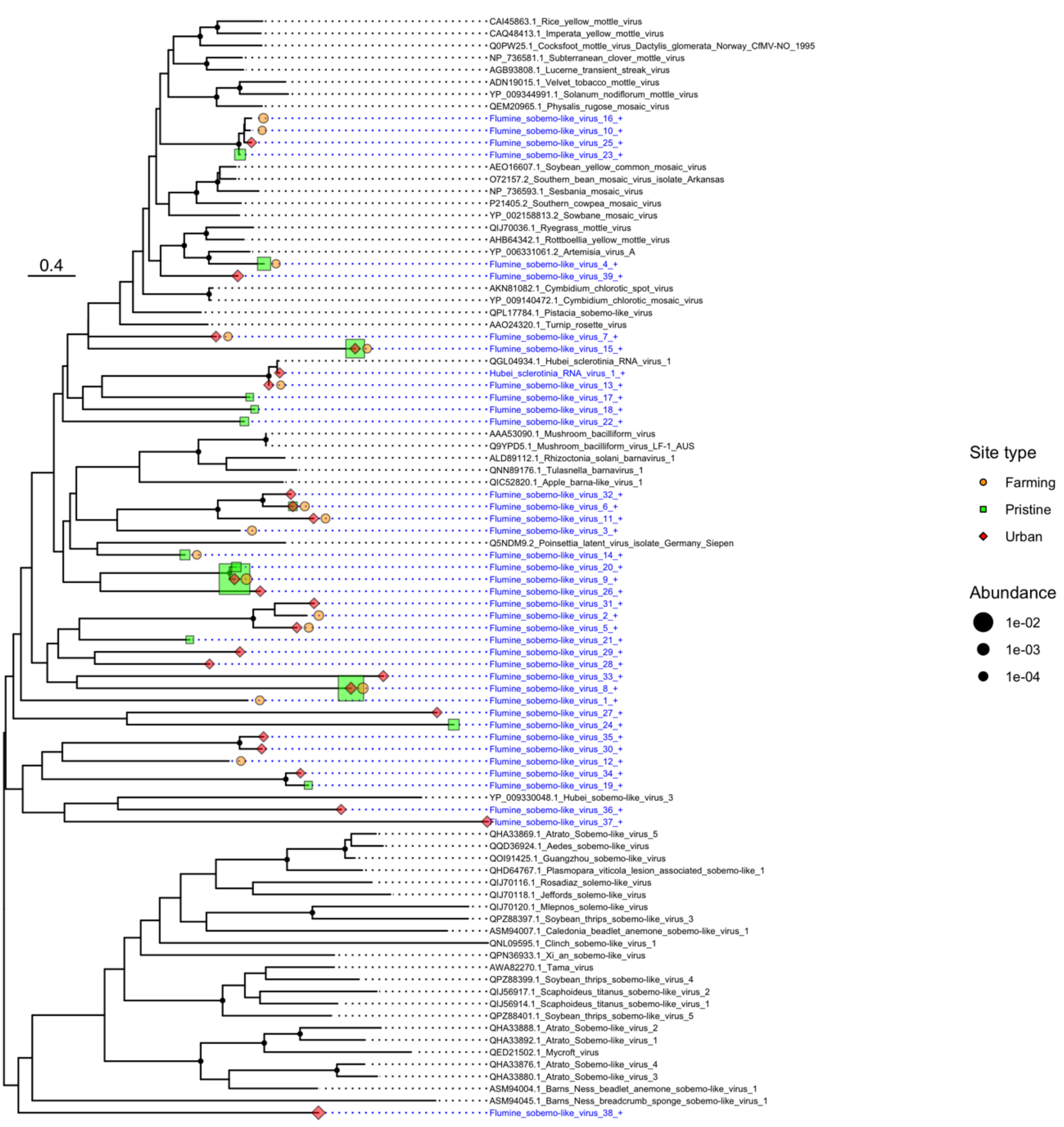
0.4
CAI45863.1_Rice_yellow_mottle_virus
CAQ48413.1_Imperata_yellow_mottle_virus
Q0PW25.1_Cocksfoot_mottle_virus_Dactylis_glomerata_Norway_CfMV-NO_1995
NP_736581.1_Subterranean_clover_mottle_virus
AGB93808.1_Lucerne_transient_streak_virus
ADN19015.1_Velvet_tobacco_mottle_virus
YP_009344991.1_Solanum_nodiflorum_mottle_virus
QEM20965.1_Physalis_rugose_mosaic_virus
Flumine_sobemo-like_virus_16_+
Flumine_sobemo-like_virus_10_+
Flumine_sobemo-like_virus_25_+
Flumine_sobemo-like_virus_23_+
AEO16607.1_Soybean_yellow_common_mosaic_virus
O72157.2_Southern_bean_mosaic_virus_isolate_Arkansas
NP_736593.1_Sesbania_mosaic_virus
P21405.2_Southern_cowpea_mosaic_virus
YP_002158813.2_Sowbane_mosaic_virus
QIJ70036.1_Ryegrass_mottle_virus
AHB64342.1_Rottboellia_yellow_mottle_virus
YP_006331061.2_Artemisia_virus_A
Flumine_sobemo-like_virus_4_+
Flumine_sobemo-like_virus_39_+
AKN81082.1_Cymbidium_chlorotic_spot_virus
YP_009140472.1_Cymbidium_chlorotic_mosaic_virus
QPL17784.1_Pistacia_sobemo-like_virus
AAO24320.1_Turnip_rosette_virus
Flumine_sobemo-like_virus_7_+
Flumine_sobemo-like_virus_15_+
QGL04934.1_Hubei_sclerotinia_RNA_virus_1
Hubei_sclerotinia_RNA_virus_1_+
Flumine_sobemo-like_virus_13_+
Flumine_sobemo-like_virus_17_+
Flumine_sobemo-like_virus_18_+
Flumine_sobemo-like_virus_22_+
AAA53090.1_Mushroom_bacilliform_virus
Q9YPD5.1_Mushroom_bacilliform_virus_LF-1_AUS
ALD89112.1_Rhizoctonia_solani_barnavirus_1
QNN89176.1_Tulasnella_barnavirus_1
QIC52820.1_Apple_barna-like_virus_1
Flumine_sobemo-like_virus_32_+
Flumine_sobemo-like_virus_6_+
Flumine_sobemo-like_virus_11_+
Flumine_sobemo-like_virus_3_+
Q5NDM9.2_Poinsettia_latent_virus_isolate_Germany_Siepen
Flumine_sobemo-like_virus_14_+
Flumine_sobemo-like_virus_20_+
Flumine_sobemo-like_virus_9_+
Flumine_sobemo-like_virus_26_+
Flumine_sobemo-like_virus_31_+
Flumine_sobemo-like_virus_2_+
Flumine_sobemo-like_virus_5_+
Flumine_sobemo-like_virus_21_+
Flumine_sobemo-like_virus_29_+
Flumine_sobemo-like_virus_28_+
Flumine_sobemo-like_virus_33_+
Flumine_sobemo-like_virus_8_+
Flumine_sobemo-like_virus_1_+
Flumine_sobemo-like_virus_27_+
Flumine_sobemo-like_virus_24_+
Flumine_sobemo-like_virus_35_+
Flumine_sobemo-like_virus_30_+
Flumine_sobemo-like_virus_12_+
Flumine_sobemo-like_virus_34_+
Flumine_sobemo-like_virus_19_+
YP_009330048.1_Hubei_sobemo-like_virus_3
Flumine_sobemo-like_virus_36_+
Flumine_sobemo-like_virus_37_+
QHA33869.1_Atrato_Sobemo-like_virus_5
QQD36924.1_Aedes_sobemo-like_virus
QOI91425.1_Guangzhou_sobemo-like_virus
QHD64767.1_Plasmopara_viticola_lesion_associated_sobemo-like_1
QIJ70116.1_Rosadiaz_solemo-like_virus
QIJ70118.1_Jeffords_solemo-like_virus
QIJ70120.1_Mlepnos_solemo-like_virus
QPZ88397.1_Soybean_thrips_sobemo-like_virus_3
ASM94007.1_Caledonia_beadlet_anemone_sobemo-like_virus_1
QNL09595.1_Clinch_sobemo-like_virus_1
QPN36933.1_Xi_an_sobemo-like_virus
AWA82270.1_Tama_virus
QPZ88399.1_Soybean_thrips_sobemo-like_virus_4
QIJ56917.1_Scaphoideus_titanus_sobemo-like_virus_2
QIJ56914.1_Scaphoideus_titanus_sobemo-like_virus_1
QPZ88401.1_Soybean_thrips_sobemo-like_virus_5
QHA33888.1_Atrato_Sobemo-like_virus_2
QHA33892.1_Atrato_Sobemo-like_virus_1
QED21502.1_Mycroft_virus
QHA33876.1_Atrato_Sobemo-like_virus_4
QHA33880.1_Atrato_Sobemo-like_virus_3
ASM94004.1_Barns_Ness_beadlet_anemone_sobemo-like_virus_1
ASM94045.1_Barns_Ness_breadcrumb_sponge_sobemo-like_virus_1
Flumine_sobemo-like_virus_38_+
Site type
Farming
Pristine
Urban
Abundance
1e-02
1e-03
1e-04

**Figure S22**

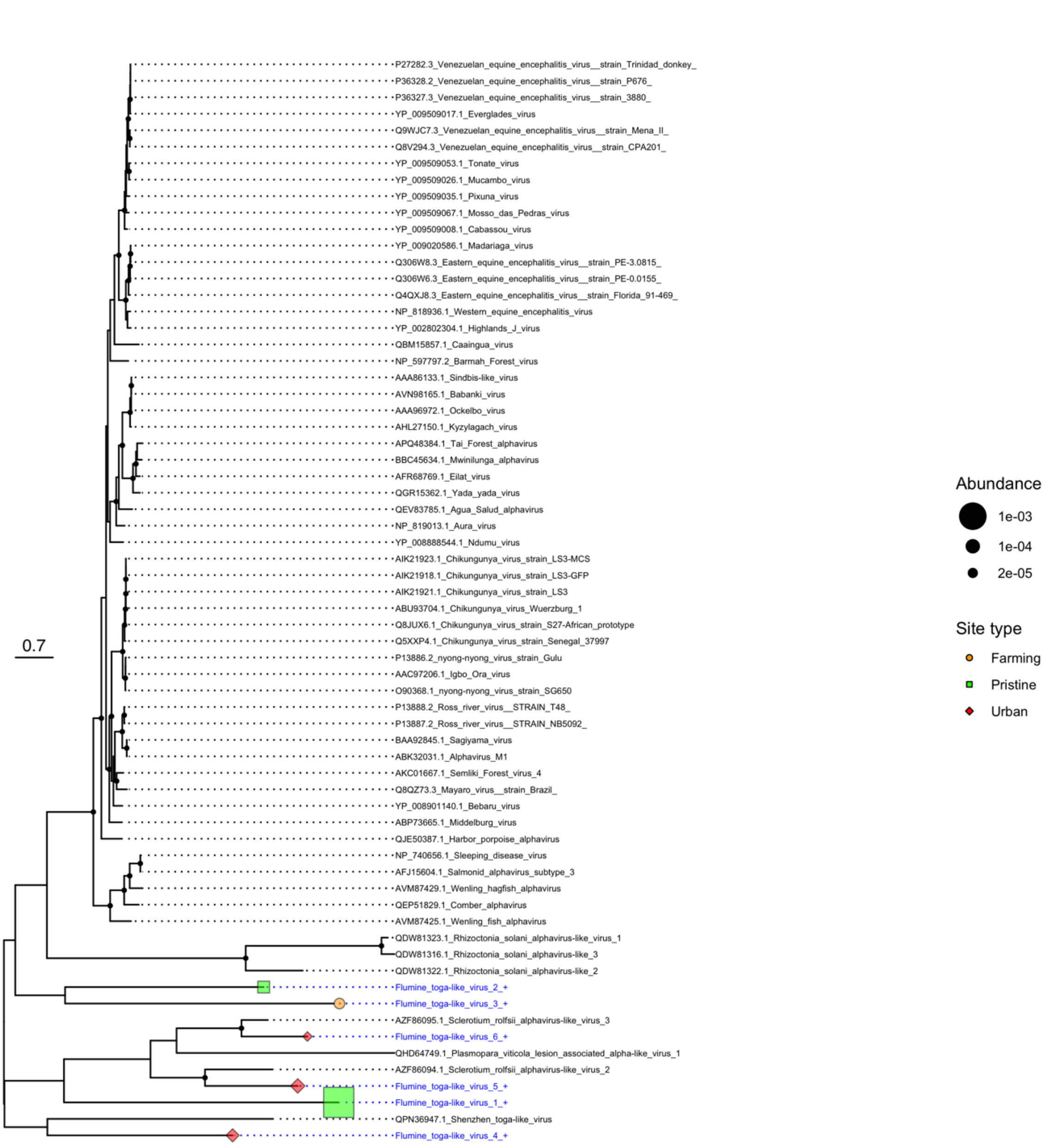
P27282.3_Venezuelan_equine_encephalitis_virus__strain_Trinidad_donkey_
P36328.2_Venezuelan_equine_encephalitis_virus__strain_P676_
P36327.3_Venezuelan_equine_encephalitis_virus__strain_3880_
YP_009509017.1_Everglades_virus
Q9WJC7.3_Venezuelan_equine_encephalitis_virus__strain_Mena_II_
Q8V294.3_Venezuelan_equine_encephalitis_virus__strain_CPA201_
YP_009509053.1_Tonate_virus
YP_009509026.1_Mucambo_virus
YP_009509035.1_Pixuna_virus
YP_009509067.1_Mosso_das_Pedras_virus
YP_009509008.1_Cabassou_virus
YP_009020586.1_Madariaga_virus
Q306W8.3_Eastern_equine_encephalitis_virus__strain_PE-3.0815_
Q306W6.3_Eastern_equine_encephalitis_virus__strain_PE-0.0155_
Q4QXJ8.3_Eastern_equine_encephalitis_virus__strain_Florida_91-469_
NP_818936.1_Western_equine_encephalitis_virus
YP_002802304.1_Highlands_J_virus
QBM15857.1_Caaingua_virus
NP_597797.2_Barmah_Forest_virus
AAA86133.1_Sindbis-like_virus
AVN98165.1_Babanki_virus
AAA96972.1_Ockelbo_virus
AHL27150.1_Kyzylagach_virus
APQ48384.1_Tai_Forest_alphavirus
BBC45634.1_Mwinilunga_alphavirus
AFR68769.1_Eilat_virus
QGR15362.1_Yada_yada_virus
QEV83785.1_Agua_Salud_alphavirus
NP_819013.1_Aura_virus
YP_008888544.1_Ndumu_virus
AIK21923.1_Chikungunya_virus_strain_LS3-MCS
AIK21918.1_Chikungunya_virus_strain_LS3-GFP
AIK21921.1_Chikungunya_virus_strain_LS3
ABU93704.1_Chikungunya_virus_Wuerzburg_1
Q8JUX6.1_Chikungunya_virus_strain_S27-African_prototype
Q5XXP4.1_Chikungunya_virus_strain_Senegal_37997
P13886.2_nyong-nyong_virus_strain_Gulu
AAC97206.1_Igbo_Ora_virus
O90368.1_nyong-nyong_virus_strain_SG650
P13888.2_Ross_river_virus__STRAIN_T48_
P13887.2_Ross_river_virus__STRAIN_NB5092_
BAA92845.1_Sagiyama_virus
ABK32031.1_Alphavirus_M1
AKC01667.1_Semliki_Forest_virus_4
Q8QZ73.3_Mayaro_virus__strain_Brazil_
YP_008901140.1_Bebaru_virus
ABP73665.1_Middelburg_virus
QJE50387.1_Harbor_porpoise_alphavirus
NP_740656.1_Sleeping_disease_virus
AFJ15604.1_Salmonid_alphavirus_subtype_3
AVM87429.1_Wenling_hagfish_alphavirus
QEP51829.1_Comber_alphavirus
AVM87425.1_Wenling_fish_alphavirus
QDW81323.1_Rhizoctonia_solani_alphavirus-like_virus_1
QDW81316.1_Rhizoctonia_solani_alphavirus-like_3
QDW81322.1_Rhizoctonia_solani_alphavirus-like_2
Flumine_toga-like_virus_2_+
Flumine_toga-like_virus_3_+
AZF86095.1_Sclerotium_rolfsii_alphavirus-like_virus_3
Flumine_toga-like_virus_6_+
QHD64749.1_Plasmopara_viticola_lesion_associated_alpha-like_virus_1
AZF86094.1_Sclerotium_rolfsii_alphavirus-like_virus_2
Flumine_toga-like_virus_5_+
Flumine_toga-like_virus_1_+
QPN36947.1_Shenzhen_toga-like_virus
Flumine_toga-like_virus_4_+
0.7
Abundance
1e-03
1e-04
2e-05
Site type
Farming
Pristine
Urban

**Figure S23**

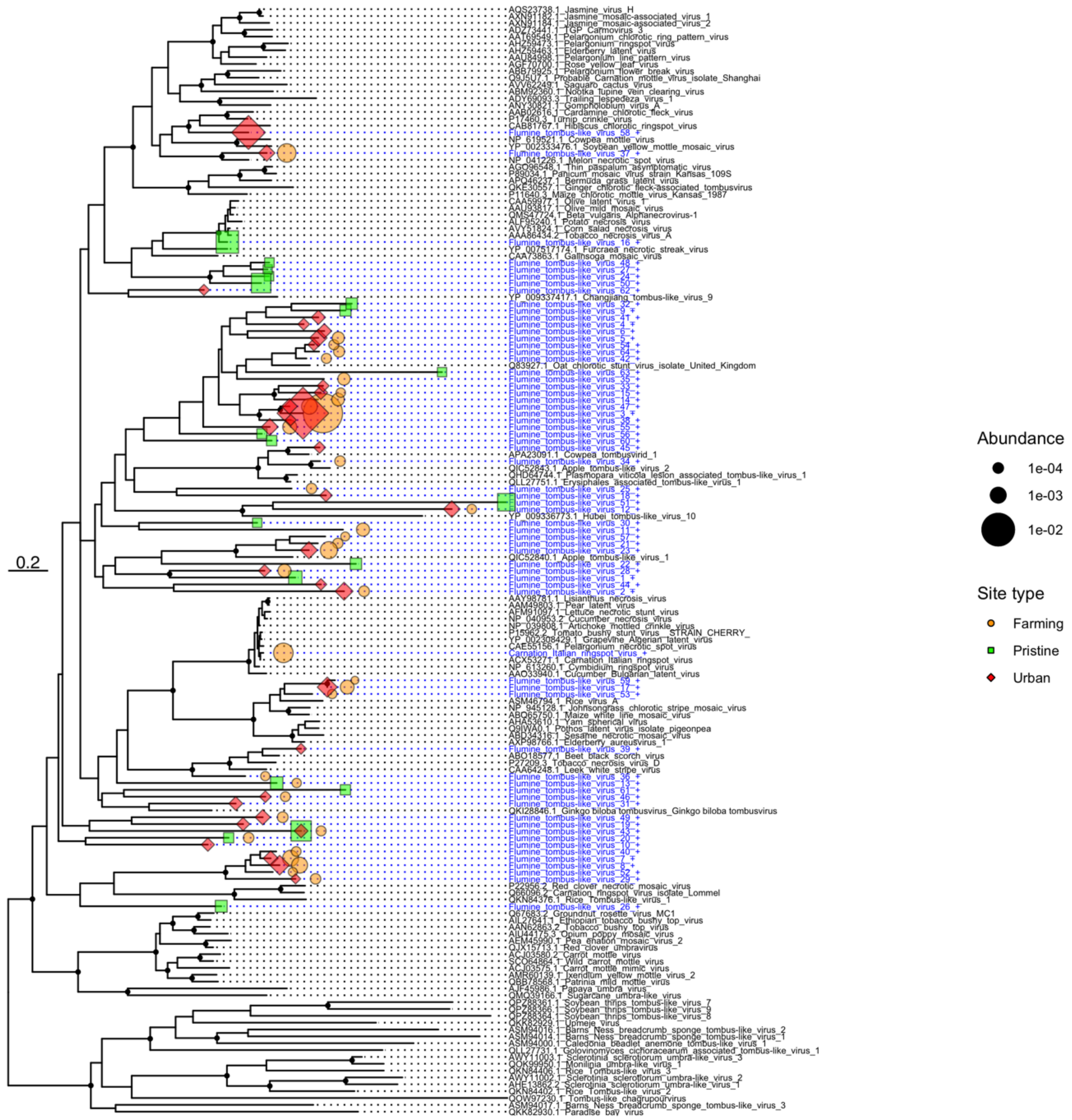
0.2
Abundance
1e-04
1e-03
1e-02
Site type
Farming
Pristine
Urban

**Figure S24**

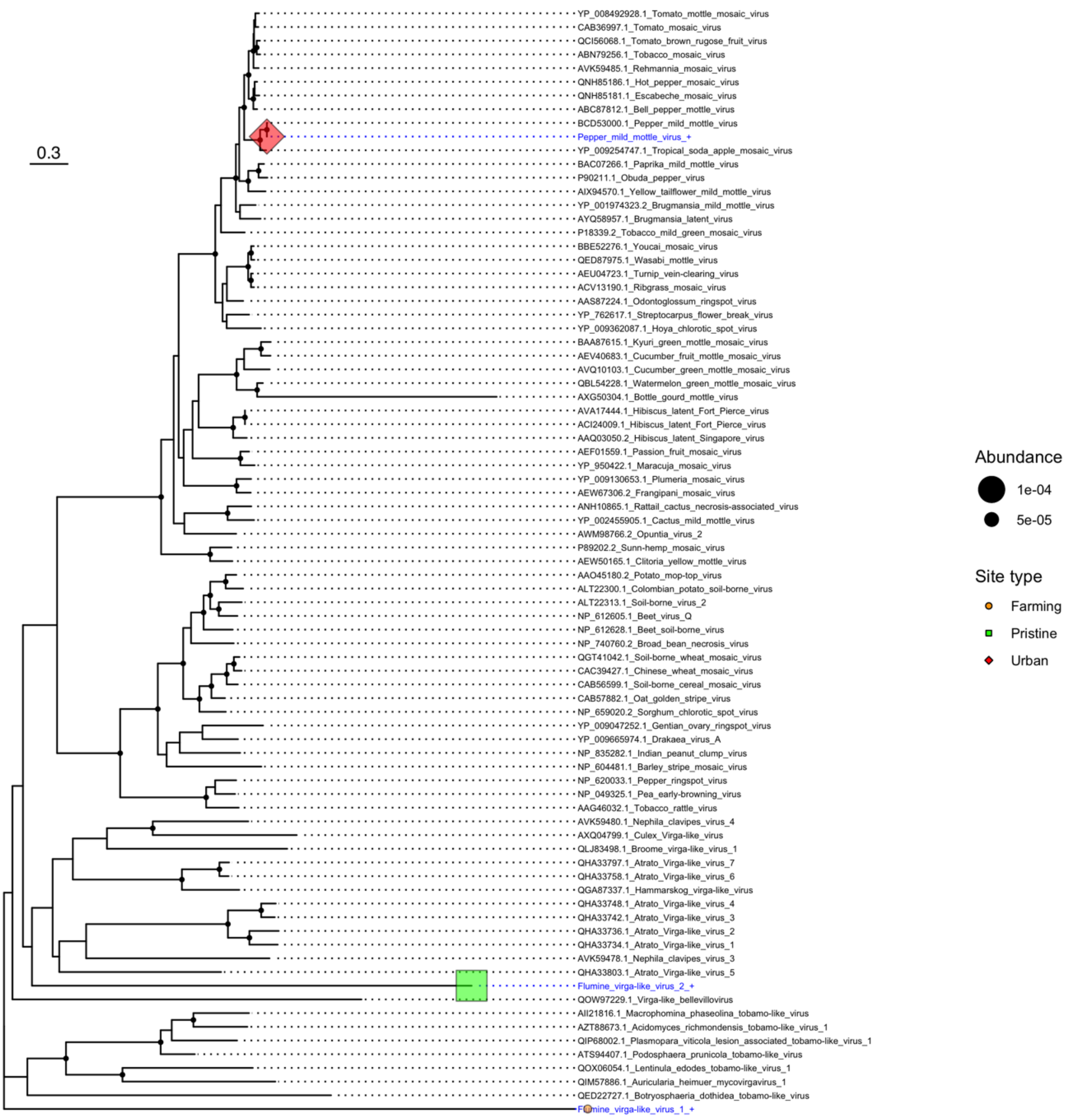
0.3
YP_008492928.1_Tomato_mottle_mosaic_virus
CAB36997.1_Tomato_mosaic_virus
QCI56068.1_Tomato_brown_rugose_fruit_virus
ABN79256.1_Tobacco_mosaic_virus
AVK59485.1_Rehmannia_mosaic_virus
QNH85186.1_Hot_pepper_mosaic_virus
QNH85181.1_Escabeche_mosaic_virus
ABC87812.1_Bell_pepper_mottle_virus
BCD53000.1_Pepper_mild_mottle_virus
Pepper_mild_mottle_virus_+
YP_009254747.1_Tropical_soda_apple_mosaic_virus
BAC07266.1_Paprika_mild_mottle_virus
P90211.1_Obuda_pepper_virus
AIX94570.1_Yellow_tailflower_mild_mottle_virus
YP_001974323.2_Brugmansia_mild_mottle_virus
AYQ58957.1_Brugmansia_latent_virus
P18339.2_Tobacco_mild_green_mosaic_virus
BBE52276.1_Youcai_mosaic_virus
QED87975.1_Wasabi_mottle_virus
AEU04723.1_Turnip_vein-clearing_virus
ACV13190.1_Ribgrass_mosaic_virus
AAS87224.1_Odontoglossum_ringspot_virus
YP_762617.1_Streptocarpus_flower_break_virus
YP_009362087.1_Hoya_chlorotic_spot_virus
BAA87615.1_Kyuri_green_mottle_mosaic_virus
AEV40683.1_Cucumber_fruit_mottle_mosaic_virus
AVQ10103.1_Cucumber_green_mottle_mosaic_virus
QBL54228.1_Watermelon_green_mottle_mosaic_virus
AXG50304.1_Bottle_gourd_mottle_virus
AVA17444.1_Hibiscus_latent_Fort_Pierce_virus
ACI24009.1_Hibiscus_latent_Fort_Pierce_virus
AAQ03050.2_Hibiscus_latent_Singapore_virus
AEF01559.1_Passion_fruit_mosaic_virus
YP_950422.1_Maracuja_mosaic_virus
YP_009130653.1_Plumeria_mosaic_virus
AEW67306.2_Frangipani_mosaic_virus
ANH10865.1_Rattail_cactus_necrosis-associated_virus
YP_002455905.1_Cactus_mild_mottle_virus
AWM98766.2_Opuntia_virus_2
P89202.2_Sunn-hemp_mosaic_virus
AEW50165.1_Clitoria_yellow_mottle_virus
AAO45180.2_Potato_mop-top_virus
ALT22300.1_Colombian_potato_soil-borne_virus
ALT22313.1_Soil-borne_virus_2
NP_612605.1_Beet_virus_Q
NP_612628.1_Beet_soil-borne_virus
NP_740760.2_Broad_bean_necrosis_virus
QGT41042.1_Soil-borne_wheat_mosaic_virus
CAC39427.1_Chinese_wheat_mosaic_virus
CAB56599.1_Soil-borne_cereal_mosaic_virus
CAB57882.1_Oat_golden_stripe_virus
NP_659020.2_Sorghum_chlorotic_spot_virus
YP_009047252.1_Gentian_ovary_ringspot_virus
YP_009665974.1_Drakaea_virus_A
NP_835282.1_Indian_peanut_clump_virus
NP_604481.1_Barley_stripe_mosaic_virus
NP_620033.1_Pepper_ringspot_virus
NP_049325.1_Pea_early-browning_virus
AAG46032.1_Tobacco_rattle_virus
AVK59480.1_Nephila_clavipes_virus_4
AXQ04799.1_Culex_Virga-like_virus
QLJ83498.1_Broome_virga-like_virus_1
QHA33797.1_Atrato_Virga-like_virus_7
QHA33758.1_Atrato_Virga-like_virus_6
QGA87337.1_Hammarskog_virga-like_virus
QHA33748.1_Atrato_Virga-like_virus_4
QHA33742.1_Atrato_Virga-like_virus_3
QHA33736.1_Atrato_Virga-like_virus_2
QHA33734.1_Atrato_Virga-like_virus_1
AVK59478.1_Nephila_clavipes_virus_3
QHA33803.1_Atrato_Virga-like_virus_5
Flumine_virga-like_virus_2_+
QOW97229.1_Virga-like_bellevillovirus
AII21816.1_Macrophomina_phaseolina_tobamo-like_virus
AZT88673.1_Acidomyces_richmondensis_tobamo-like_virus_1
QIP68002.1_Plasmopara_viticola_lesion_associated_tobamo-like_virus_1
ATS94407.1_Podosphaera_prunicola_tobamo-like_virus
QOX06054.1_Lentinula_edodes_tobamo-like_virus_1
QIM57886.1_Auricularia_heimuer_mycovirgavirus_1
QED22727.1_Botryosphaeria_dothidea_tobamo-like_virus
Flumine_virga-like_virus_1_+
Abundance
1e-04
5e-05
Site type
Farming
Pristine
Urban

### Figure S25
